# The phylogenetic affinities of Chaetognathifera, with considerations of systematic error and the robusticity of macrosyntenic results

**DOI:** 10.64898/2026.06.08.730799

**Authors:** James F. Fleming, Nickellaus G. Roberts, Holger Herlyn, Wilko Ahrlichs, Kevin M. Kocot, Torsten H. Struck

## Abstract

Chaetognathifera, a superphylum comprising Syndermata (Rotifera including Acanthocephala), Micrognathozoa, Gnathostomulida and Chaetognatha, is a complex grouping generally recovered as the sister to all other Lophotrochozoa. However, phylogenetic relationships within this group are controversial, in part due to poor sampling, resulting in two key questions. The first is whether Gnathostomulida or Chaetognatha represent the sister group to Syndermata+Micrognathozoa. The second is the phylogenetic position of the former phylum Acanthocephala within Syndermata. Here, we present the first study of the phylogenetic affinities of Chaetognathifera with genomic representation from all major phyla, and explore the potential of macrosynteny to better understand these relationships. For this latter aspect, we also developed a new jackknifing procedure to assess the robustness of linkage groups inferred by macrosyntenic analyses. We show that the phylogenetic relationships between these clades are corroborated through a variety of gene selection and analysis methodologies. This provides clear evidence of Acanthocephala as a derived clade within Syndermata as sister to Seisonidea, and that Gnathostomulida is sister to Syndermata+Micrognathozoa, with Chaetognatha as the earliest diverging clade within Chaetognathifera. On the other hand, we found that macrosyntenic patterns cannot resolve this question. Moreover, almost all possible linkage groups involving chaetognathiferan species lack robusticity and hence, should not be considered reliable. As a consequence so far, in Chaetognathifera none of the bilaterian ancestral linkage groups can be reliably found and independent massive chromosomal rearrangements occurred. We therefore strongly suggest that studies of macrosynteny should not only assess the significance of possible linkage groups, but also the robusticity of these linkage group inferences. Furthermore, we also present a script for this purpose, which can be found at: https://github.com/JFFleming/MacrosyntenicJackknife

## Introduction

Chaetognathifera is a superphylum within Lophotrochozoa (also known as Spiralia), comprised of four phyla – Syndermata (“Rotifera” including Acanthocephala), Micrognathozoa, Gnathostomulida and Chaetognatha. Chaetognathifera was first strongly recovered based on phylogenomic data^1–4^ and is supported by interpretations of the fossil record^5–7^. Species in Chaetognathifera are usually less than 1cm in size, but some can be as large as 12cm in Chaetognatha and 70cm in Acanthocephala^8,9^. Moreover, almost all members larger than 1mm belong to either Chaetognatha or Acanthocephala^8,10^. These phyla are highly variable and range from poorly described and recently discovered, to incredibly diverse. Except for Acanthocephala, they are united by chitinous structures used for feeding. In Chaetognatha, these structures take the form of spines that are used to grasp prey items using ambush hunting, and their distinct body shape grants the phylum its common name: arrow worms. In rotifers, gnathostomulids and micrognathozoans, however, these structures take the form of “jaws” - complex, articulated and highly variable structures that can be used for grazing, grasping, grinding, or piercing of smaller sized prey, algae or bacteria^8,10^. These phyla have been grouped together as Gnathifera due to the strong similarities between their jaw apparatuses and later molecular data^11,12^. These “jaws” reach incredible levels of complexity in rotifers and micrognathozoans. In all cases, these jaws are used in combination with a muscular pharynx allowing for complex movements.^13^ In contrast, despite - or perhaps due to - their parasitic lifestyle, Acanthocephala lack a jaw. While they have a range of hosts across both gnathostome vertebrates and mandibulate arthropods, and their life cycle involves inhabiting two distinct hosts^14,15^, they are most notable as economically significant parasites of fish^8,16,17^.

The phylogenetic relationships among these four groups, however, are unclear (Figure 1)^1–3,18,19^. Understanding these relationships - and the stability of these branches - builds a firm foundation for a broader understanding of the evolution including the ancestral body plan of Chaetognathifera^7^. This could help us understand more about the early evolution of Lophotrochozoa as a whole. Lophotrochozoa comprises more than 50% of the known animal phyla^18^. Besides the four already mentioned, these are diverse phyla like Entoprocta, Mollusca, Gastrotricha, Bryozoa, Annelida, Nemertea and Platyhelminthes. While some progress - such as the sister group relationship of Chaetognathifera to Platytrochozoa - has been achieved using modern phylogenomic analyses, many phylogenetic controversies remain, as these studies also recover an array of diverse arrangements of the phyla it comprises^18,19^. Accordingly, a firm understanding of the evolution of Chaetognathifera will allow us to root and polarize evolutionary trajectories of morphological, developmental and other character traits within its sister group Platytrochozoa in future studies.

**Figure 1:**
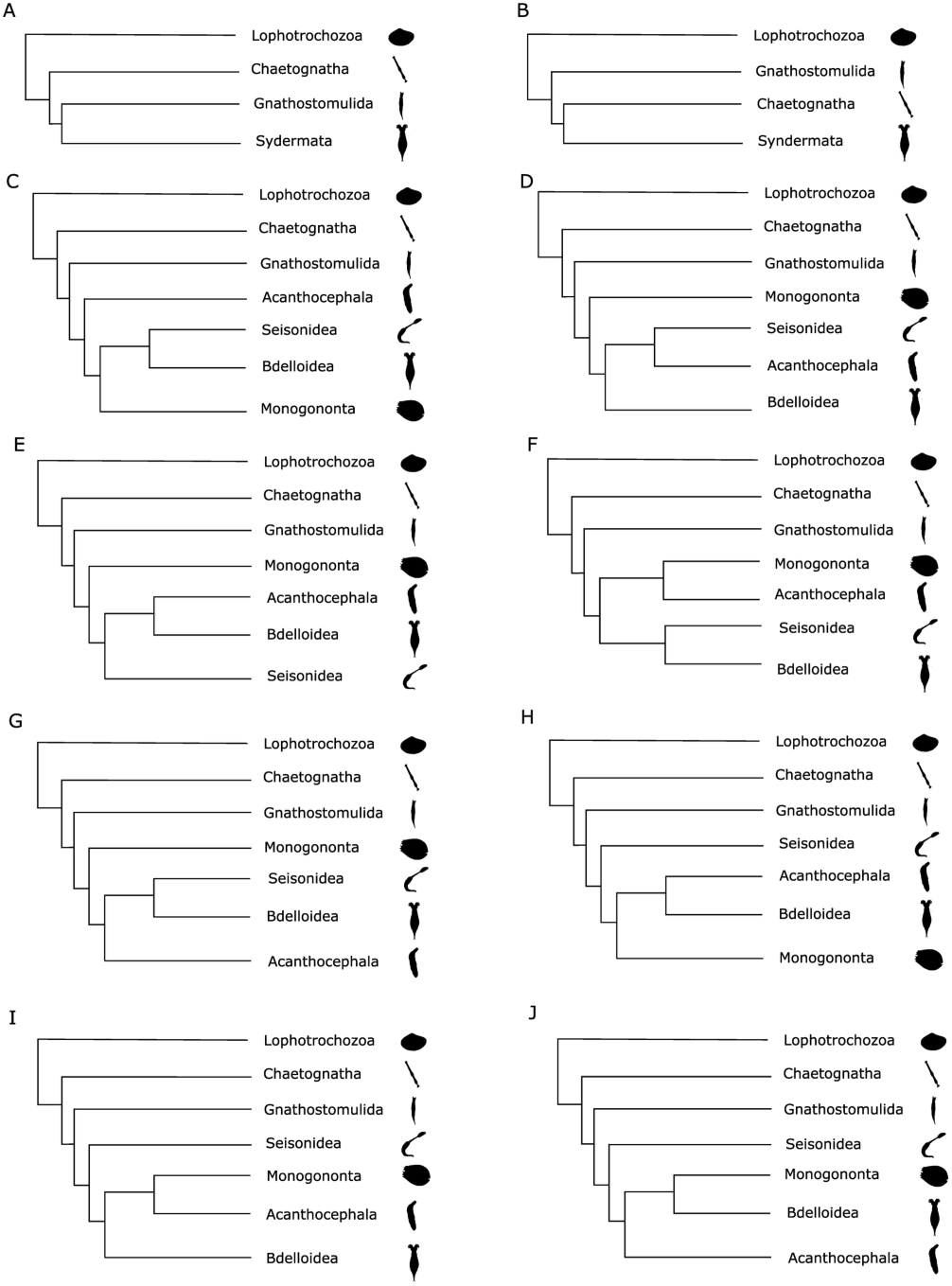
10 cladograms depicting current controversies within Chaetognathifera. Panels a and b display hypotheses related to the earliest diverging group within Chaetognathifera, whilst c through g display hypotheses related to the internal phylogeny of Syndermata with respect to Acanthocephala and panels h through i display hypothesis related to the internal phylogeny of Syndermata with respect to Seisonidea. Panel a) shows the Chaetognatha-first hypothesis. Panel b) shows the Gnathostomulida-first hypothesis. Panel c) shows traditional “Rotifera” as monophyletic with Acanthocephala as its sister. Panel d) shows Pararotatoria, where Seisonidea is sister to Acanthocephala. Panel e) shows Lemniscea, where Acanthocephala is sister to Bdelloidea. Panel f) shows Monogononta+Acanthocephala. Panel g) shows Hemirotifera with Acanthocephala is sister to Seisonidea+Bdelloidea. Panel h) shows Seisonidea as the sister group to the remaining Syndermata, with Acanthocephala and Bdelloidea as sisters (Seisonidea-AB). Panel i) shows Seisonidea as the sister group to the remaining Syndermata, with Acanthocephala and Monogononta as sisters (Seisonidea-AM). Panel j) shows Seisonidea as the sister group to the remaining Syndermata, with Bdelloidea and Monogononta as sisters (Seisonidea-BM).

When considering the evolution of Chaetognathifera, two pertinent questions have yet to be confidently resolved. The first concerns the relationships among the four phyla in the group: Syndermata, Micrognathozoa, Chaetognatha and Gnathostomulida. The monophyly of Syndermata and Micrognathozoa is consistently supported by molecular studies containing both phyla^18^, confirming the morphological conclusions based on the shared possession of apical aggregations of fibrous proteins in the epidermis^11,12,20,21^. In contrast, based on three phylogenomic studies, the relationship of these two phyla to Chaetognatha and Gnathostomulida remains disputed, presenting two plausible scenarios: “Chaetognatha-first” or “Gnathostomulida-first”^1–3^ (Figure 1A & B). Establishing the sister group of the remaining Chaetognathifera has considerable ramifications for understanding the broader history of the superphylum. One key question remains particularly relevant - did the complex jaw of Gnathifera, and the musculature that controls it, evolve once or twice convergently? In a Chaetognatha-first scenario, the gnathiferan jaw evolved only once, whilst in the Gnathostomula-first scenario, the gnathiferan jaw is either a synapomorphy of Chaetognathifera that has been independently lost in Chaetognatha, or has evolved twice independently, once in Gnathostomulida and once in the shared ancestor of Syndermata and Micrognathozoa. Resolving this question has been frustrated by the lack of a high-quality gnathostomulid genome, and topologies have varied significantly between studies. Only three phylogenomic studies to date have included transcriptomic data from both Chaetognatha and Gnathostomulida: Kocot et al (2017)^2^, Marletaz et al. (2019)^3^ and Laumer et al (2019)^1^. Kocot et al and Laumer et al both recovered Chaetognatha-first topologies, whilst Marletaz et al recovered Gnathostomulida-first, suggesting discordance in sampling and modelling considerations^22,23^.

The second pertinent phylogenetic question concerns the position of Acanthocephala, a former phylum, within Syndermata. All known acanthocephalans are obligate parasites. When the group was first named, in 1771, they were placed within the helminths. As systematic revisions progressed throughout the centuries, Acanthocephala found themselves morphologically placed as a phylum within Gnathifera at large - and Syndermata in specific^8,11,12,24^. Syndermata is also often called Rotifera in a broader understanding. A close relationship of Acanthocephala and “Rotifera” is supported among others by the syncytial epidermis with an apical intracellular aggregation of fibrous proteins, also called lamina, and intrusions of the outer epidermal cell membrane with bulbs through the lamina. Molecular analyses have since confirmed that Acanthocephala represents a highly derived group within Syndermata^25–27^. However, the phylogenetic position of Acanthocephala within Syndermata is matter of ongoing debate. “Rotifera” is divided into three classes: Seisonidea, Monogononta and Bdelloidea. A stricter Rotifera hypothesis presents Rotifera and Acanthocephala as two separate, monophyletic phyla sharing a common ancestor, though this has not been recovered by molecular data (Figure 1C). However, three alternative hypotheses present Acanthocephala within Syndermata as more closely related to Seisonidea and/or Bdelloidea (Figure 1D, E & G). The Pararotatoria hypothesis places Acanthocephala as sister to Seisonidea, the Lemniscea hypothesis places Acanthocephala as sister to Bdelloidea, and the Hemirotifera hypothesis places Monogononta as the sister to a clade comprising Acanthocephala, Bdelloidea and Seisonidea. This means that the latter is congruent with the two former hypotheses but also to a scenario where Acanthocephala are sister to Bdelloidea plus Seisonidea (which is how this term is used within the remainder of this study, as this specific hypothesis otherwise lacks a name). Finally, it has also been suggested that Seisonidea is sister to all other syndermatans and accordingly Acanthocephala more closely related to either Bdelloidea (congruent with the Lemniscea hypothesis), Monogonota or both together (Figure 1H-J)^28^.

This discordance is potentially driven by the highly specialised nature of Acanthocephala themselves. As obligate parasites, they are both morphologically and molecularly highly divergent from other Syndermata^8^. This presents the possibility that this discordance is driven by long branch attraction or compositional heterogeneity, two key forms of methodological incongruence that can frustrate phylogenetic analyses. Recent work, focused on selecting genes less prone to methodological incongruence, has suggested that Pararotatoria or Hemirotifera may be favoured over the other hypotheses, and mounting evidence from both genetic and morphological data now suggests that Acanthocephala represent a particularly divergent lineage within Syndermata as sister to Seisonidea^25,26,29^. However, prior genomic studies have been hindered by issues of outgroup selection. Many clades within wider Lophotrochozoa that would otherwise make ideal outgroups to resolve this problem are themselves particularly prone to methodological incongruence, and outgroup selection has been shown to have a significant effect on topological recovery in phylogenetic analysis^18,30^. In this respect, a dataset containing genomic data from both Chaetognatha and Gnathostomulida serves to reinforce the inferences that can be made about Syndermata.

Besides understanding morphological evolution, a better understanding of relationships within Chaetognathifera as well as genomic information for all major phyla will help us to better understand genomic evolution within Chaetognathifera but also in Lophotrochozoa in general. Macrosyntenic analyses may be a way to better understand genomic events, possible evolutionary synapomorphies and to assess the plausibility of certain phylogenetic hypotheses^31,32^. Here, fusions and fissions of entire chromosomes, ancestral linkage groups (ALG), and fusion-with-mixing events can provide new perspectives on how genomes evolve^4,32–38^. For example, with respect to Lophotrochozoa it has been suggested that four such fusion-with-mixing events could support the monophyly of Lophotrochozoa^4,33,35,36,38^. In a recent publication of a chaetognath genome^39^, it was suggested that at least two of these events could also be found in the chaetognath genome of *Paraspadella gotoi*, while the other two could neither be confirmed nor rejected. On the other hand, a closer examination of the detection of the ALGs assignments revealed that the inference of fusion with mixing events was highly dependent on the species used in the pairwise comparison to *P. gotoi*^18^. This resulted in strong incongruencies regarding which ALG should be assigned to which chromosome. Hence, it is still uncertain if bilaterian ancestral linkage groups and specific lophotrochozoan events can be detected in Chaetognathifera.

Here, we present a comprehensive analysis of both critical systematic questions, pursuing multiple lines of evidence to establish the identity of the sister group to the remaining Chaetognathifera, and the position of Acanthocephala within the group. Guided by comprehensive pre-analysis gene selection, and using a combination of the Canary Sequence methodology^40^, CAT Posterior Mean Site Frequency^41^, and gene tree approaches^42^, we clearly show that Acanthocephala should be considered a clade nested within Syndermata, as sister to the Seisonidea, and that Chaetognatha represent the earliest diverging group of the broader Chaetognathifera. The macrosyntenic approach^43^ was inconclusive in the phylogenetic respect and it rather strongly suggests the occurrence of massive fusion-with-mixing events independently in all chaetognathiferan phyla examined in this study - and a lack of detectable bilaterian ancestral linkage groups. In contrast, our study shows the importance of assessing and confirming the robusticity of detected pairwise linkage groups in addition to significance determination. Only the latter is routinely conducted to date for macrosyntenic studies, while the former is lacking. We present such a jackknifing approach, alongside a script to implement it using MacrosyntR^43^, available at the Github repository associated with this study: https://github.com/JFFleming/MacrosyntenicJackknife

## Methods

### Sample collec9on and DNA sequencing

We collected *Gnathostomula armata* from the German North Sea island Sylt (55.026227, 8.434112). The specimen (Gn60, Biosample number SAMN60462428) was extracted from sediment of the oxygenated top layer (1 cm) in an area with *Arenicola marina* tubes. The monogonontan rotifer *Euchlanis meneta* (D05_EMe, Biosample number SAMN60533003) was collected from the lake Kleiner Bornhorster See in Germany (53.1837094057668, 8.269197107622258). *Spadella aff. kappae* (KK3082.1L, Biosample number 60546929) was collected from sediment samples near Davis Reef in the Florida Keys (24.9285, - 80.5070). One female specimens of *Pomphorhynchus laevis* (*sensu lato*)^44^ (Plaevis2, Biosample number SAMN60532144) was collected in August 2019 from the intestine of a naturally infected leuciscid fish (*<u>Squalius cephalus</u>*, Leuciscidae), caught and sacrificed by an authorized person (game fisher) in the river Fulda near suburb Bonaforth of the town Hannoversch Münden (Germany). For reasons of brevity, the latter two will referred to as *Spadella kappae* and *Pomphorhynchus laevis* in the following.

Upon collection, all samples were flash frozen (*P. laevis*, *G. armata*, *E. meneta*) or preserved in the REPLI-g storage buffer and then flash frozen (*S. kappae*). DNA of *P. laevis* was first extracted using the Nanobind Tissue Big DNA kit (Circulomics) and whole-genome amplification was performed using the REPLI-g Single Cell kit (Qiagen) with about 5 ng of extracted DNA. For the other three species, the whole genome was directly amplified from tissue using REPLI-g Advanced Single Cell or REPLI-g Single Cell kit. The quality and concentration of DNA was checked at each step using Nanodrop (Thermo Fisher Scientific), Qubit (Thermo Fisher Scientific) and Fragment Analyzer (Agilent). Library preparation and PacBio HiFi sequencing was conducted by the Norwegian Sequencing Centre (Oslo, Norway) or Discovery Life Sciences (Huntsville, AL, USA) on a Sequel II (Pacific Biosciences) plaqorm. This included shearing using G-Tubes, cleaning up with either Ampure XP beads and/or PacBio Short Read Eliminator kit and the standard input protocol. The procedures followed generally the one by Roberts et al. (2024)^45^, with the exception of the amplification of *G. armata.*, which incorporated a modified version of the Repli-G amplification protocol (See supplemental information). All samples are available under BioProject accession number PRJNA1471681.

### Genome Assembly

First, bam files were converted to fastq files using BEDTools 2.29.2^46^. The distribution of read lengths was assessed using custom bash and R scripts. The kmer distribution was determined using Jellyfish v.2.3.0^47^ up to coverage of 10.000 and 1 million and then GenomeScope2.0^48^ to obtain kmer-based genome size and other parameters with default settings for a diploid genome. Ploidy levels were estimated using Smudgeplot v.0.2.4^48^ with default settings and the counts from the Jellyfish analysis. The genome was assembled using hifiasm v.0.25.0^49–51^ with default settings. Next, applying the BlobToolKit v. 4.4.6^52^ filter option only contigs of the primary assemblies were kept fulfilling certain criteria. The selection of the criteria was guided by allowing only slight decreases in the BUSCO score. For *G. armata*, the criteria were a GC content between 0.38 and 0.41, a coverage between 130x and 300x, and a length above 500,000 bp. For *E. meneta*, contigs with a GC content between 0.24 and 0.30 and coverage between 20x and 200x were kept, while contigs matching against non-metazoan taxa except for matches against Ascomycota and all in the category “Others” were removed. For *P. laevis*, all contigs with a match against non-metazoan taxa except for the ones in category “Others” were removed as well as all that did not have a GC content between 0.28 and 0.40, a coverage between 80x and 200x, or a length above 50,000 bp. For *S. kappae*, contigs with a GC content between 0.38 and 0.50, a coverage between 50x and 400x and a length above 40,000 bp were kept, while contigs matching against non-metazoan taxa except for matches against Ascomycota and all in the category “Others” were removed.

After the filtering, Purge_Dups v.1.0.1^53^ with an expected identity ≥95% and keeping secondary and supplementary alignments was applied to purge potential duplicates. Then, the assemblies were further investigated using BlobToolKit as well as custom scripts for BLAST searches to detect any remaining non-metazoan and possible host contamination in the case of *P. laevis*. A detailed description of our assembly and quality assessments specific to each species, alongside the quality metrics for each genome, is available in supplemental information.

## Dataset Assembly

In addition to the four new genomes generated for this study, we downloaded all available Chaetognathifera genomes available from NCBI as of March 17^th^ 2026. We then conducted a BUSCO 6.0.0^54^ search against the lophotrochozoa_odb12 dataset against each of the genomes. We first excluded all taxa from the dataset with less than 50% BUSCO completeness. This threshold was chosen as a balance between assembly quality and ensuring the inclusion of the sole seisonid genome, *Seison nebaliae*, with 51.3% completeness, and at least two acanthocephalan genomes, as the second-most complete genome from the phylum, our assembly of *P. laevis*, has a BUSCO completeness of 59.6%. This resulted in the exclusion of four genomes and a dataset size of 110 ingroup genomes, split across 29 species.

We then excluded all BUSCO genes with less than 50% occupancy across these genomes - those present in fewer than 55 taxa. This produced an initial 50% occupancy dataset, comprising 646 sequences, which were then each independently aligned using MAFFT v7.511^55^ and assessed for entropic site saturation (as determined by the DE-Score^56^), compositional heterogeneity (as determined by nRCFV^57^) and branch length heterogeneity (as determined by the LB-Score^58^), and ranked based on their performance in this analysis. Of these 646 genes, any genes ranked in the bottom third (<431^st^ position) of the dataset by any of these metrics were then discarded, resulting in a selection of 138 well-performing genes across the taxa.

Taxa were then further selected based on their performance within this dataset. Over the 138 genes, each taxon was ranked based on the following criteria:

- The number of genes with a DE-Score below the critical DE-Score.
- The number of genes with a LB-Score greater than two standard deviations from the mean.
- The number of genes with a taxon-specific nRCFV greater than two standard deviations from the mean.

For each of the 29 species, the genome with the lowest sum of these “red flags” was retained. In the case of a tie between genomes, the genome with the highest single copy BUSCO score was selected. In the case of *Brachionus asplanchnoidis*, no individual genome recovered fewer than 10 “red flags” over the three individuals assessed. Considering the known levels of intraspecific genome size variation in this species, the two genomes with the fewest number of red flags were selected^59^. This resulted in a complete ingroup of 30 genomes.

Our outgroup was assembled in reference to our 646 gene dataset. Of the 406 chromosome-level available non-chaetognathiferan genomes we downloaded from NCBI, we first removed those with less than 80% occupancy across the previously selected 646 genes, resulting in the exclusion of 26 genomes. Then, each gene in the resulting 380-genome dataset was aligned using MAFFT v7.511^55^ and assessed using the same criteria as the ingroup taxa:

- The number of genes with a DE-Score below the critical DE-Score.
- The number of genes with a LB-Score greater than two standard deviations from the mean.
- The number of genes with a taxon-specific nRCFV greater than two standard deviations from the mean.

10 Mollusca, 10 Annelida, two Nemertea and one Brachiopoda genomes were then retained as the least incongruent genomes in each phyla across the sampled range of outgroup taxa. This curated outgroup was then concatenated to each of the selected test ingroup datasets, to form a final full dataset of 53 genomes. These genes were then also aligned using MAFFT v7.511^55^.

### Phylogenetic Assessment: Canary Sequence Methodology

To assess the effect of potentially problematic sequences on phylogenetic reconstruction, we adopted the Canary Sequence Methodology to identify genomes that might be the cause of topological incongruence in our dataset^40^. To first identify the sequences of interest, we assessed our final, 138 gene alignment using the LB-Score, nRCFV and DE-Score, and identified taxa that had a DE-Score below the critical DE-Score, or an LB-Score or taxon-specific nRCFV greater than two standard deviations from the mean of the dataset^56–58^.

The seven sequences identified this way were selected as sequences of interest and removed from the full dataset. This allowed us to produce a base dataset of 46 genomes (53 genomes minus the seven sequences of interest) and seven checking datasets of 47 genomes. The full dataset, base dataset and all seven checking datasets were then assessed under the LG+F model in IQTree v2.4.0^60^. The resulting topologies were then examined to determine which sequences were potentially problematic, and which were potentially canary sequences, which resulted in a final dataset of 50 genomes, as no genomes were found to be topologically problematic, but three sequences of interest were found to be significantly affected by compositional heterogeneity (Supplemental Data). The topology for this final dataset was also constructed in IQ-Tree2 v2.4.0^60^ using the LG+F model, with 5000 UF bootstrap replicates.

### Phylogenetic Assessment: CAT-PMSF

We then assessed the plausibility of each of the 10 topological possibilities using CAT-PMSF, under the testing protocol established by Giacomelli et al 2025^61^, and using the full 53 taxa dataset (as the CAT model is specifically designed to account for compositional heterogeneity)^41,61,62^. Using pb_mpi and the CAT-Poisson model^41,63^, 10 input datasets were constructed, using a combination of the 138-gene concatenated dataset on a fixed topology corresponding to Chaetognatha-first, Gnathostomulida-first, Rotifera (Rotifera+Acanthocephala), Hemirotifera (Seisonidea+Bdelloidea as sister to Acanthocephala), Lemniscea (Bdelloidea+Acanthocephala), Monogonota-sister (Monogononta+Acanthocephala), Pararotatoria (Acanthocephala + Seisonidea), and three topologies that fixed Seisonidea as the sister to Acanthocephala and the remaining classes of Rotifera: Seisonidea-First with Monogononta-sister (Acanthocephala+Bdelloidea), Seisonidea-First with Bdelloidea-sister (Acanthocephala+Monogononta) and Seisonidea-First with Acanthocephala-sister (Bdelloidea+Monogononta).

In topologies which interrogated the earliest diverging phylum within Chaetognathifera, Pararotatoria was fixed, as it was the topology recovered by the final tree dataset. Similarly, in topologies which interrogated the position of Acanthocephala with respect to “Rotifera”, Chaetognatha-first was fixed for the same reason. As a result, both the Chaetognathifera-first and Pararotatoria datasets are identical, and in turn identical to the final tree recovered by the canary sequence methodology in the prior step.

Once each PMSF chain reached an effective sample size greater than 300 when considered with a 33% burnin, with a minimum of 15,000 generations, as established by tracecomp, the chain was terminated, and site profile files were generated using read_pbmpi. These siteprofile files were converted to IQ-Tree format site frequency input files^41^. A phylogenetic tree for each hypothesis, using these site frequency input files, was then constructed in IQ-Tree v2.4.0^60^ under the Poisson model, alongside 5000 UF bootstrap replicates.

### Phylogenetic Assessment: Gene Tree Concordance (DiscoVista)

To assess gene tree concordance for the phylogenetic relationships of Chaetognathifera, independent gene trees were constructed for each of the 646 genes that formed the 50% occupancy matrix. These trees were constructed in IQ-Tree v2.4.0^60^ under the best fitting model for that gene as determined by ModelFinder. To test the identity of the earliest diverging phylum within Chaetognathifera, this dataset was then filtered to include only those trees with a representative from both Chaetognatha and Gnathifera. The resulting 350 trees were then analysed under DiscoVista^42^, and the number of trees that strongly supported and rejected each hypothesis were noted and compared. Following this, to examine the effect of methodological incongruence on the dataset, the same strategy was applied to the gene trees recovered from genes in the top 138 genes according to our earlier analysis. This resulted in a filtered dataset of 83 gene trees, which were then analysed under DiscoVista, and the number of trees that strongly supported and rejected each hypothesis were noted and compared.

To test the placement of Acanthocephala within Syndermata, the 646 gene trees were filtered to include only those trees that included at least one member of each class of “Rotifera”, and at least one acanthocephalan. The resulting 294 trees were then analysed under DiscoVista^42^, and the number of trees that strongly supported and rejected each hypothesis were noted and compared. DiscoVista is unable to assess discordance within complex constraints, as it looks only for a monophyletic clade comprising all taxa within the input clades (including polyphyly). As a result of this, it lacks a way to distinguish between (Seisonidea,(Bdelloidea,Acanthocephala)), and (Bdelloidea,(Seisonidea,Acanthocephala)) if we are also interested in the relationship of these clades to Monogononta. As such, we used monophyletic constraints for each pair of classes: Bdelloidea+Seisonidea (Hemirotatoria), Bdelloidea+Acanthocephala (Lemniscea), Seisonidea+Acanthocephala (Pararotatoria), Acanthocephala+ Monogononta (AcanthoMono), Bdelloidea+Monogononta (BdelloMono), Monogononta+Seisonidea (MonoSeison), and two larger constraints: Rotifera (Monogononta+Bdelloidea+Seisonidea) and SeisonFirst (Monogononta+Bdelloidea+Acanthocephala).

Following this, to examine the effect of methodological incongruence on the dataset, the same strategy was applied to the gene trees recovered from genes in the top 138 genes according to our earlier analysis. This resulted in a filtered dataset of 70 gene trees, which were then analysed under DiscoVista, and the number of trees that strongly supported and rejected each hypothesis were noted and compared.

### Phylogenetic Assessment: Macrosyntenic Analysis and “The Macrosyntenic Jackknife”

To assess the evolution of macrosynteny within Chaetognathifera, we analysed two publicly available chromosome-scale genomes representing Chateognatha and Syndermata: *Paraspadella gotoi* and *Adineta vaga*. We supplemented these publicly available genomes with our own assembly of *Gnathostomula armata*, and selected the publicly available genomes of *Branchiostoma floridae* and *Mizuhopecten yessoensis* as outgroups.

We first obtained the conserved metazoan ancestral linkage groups using the default MacrosyntR comparison of *Branchiostoma* and *Mizuhopecten*^43^. We used the pair comparison table derived from this analysis to derive the conserved metazoan ancestral linkage groups in each of our three study taxa, and then conducted paired comparisons of all five taxa to one another. This resulted in 10 datasets, referred to as Conserved Linkage Group (CLG) datasets, and detailed in Table 1. We then independently compared the macrosyntenic arrangements within Chaetognathifera using rbhXpress to establish whether novel Chaetognathifera linkage groups could be determined, resulting in a further three “Independent” datasets (Table 1).

**Table 1:** Describing the size of each subsampled macrosyntenic input dataset, and the sampling regime that derived the original ortholog pair file. CLG = *Branchiostoma x Mizuhopecten* conserved linkage groups; Independent = Directly inferred linkage groups.

| Taxa 1 | Taxa 2 | Comparison Regime | Subsample Input Dataset Size |
| --- | --- | --- | --- |
| *Branchiostoma* | *Mizuhopecten* | CLG (control) | 1807 |
| *Branchiostoma* | *Adineta* | CLG | 1224 |
| *Branchiostoma* | *Gnathostomula* | CLG | 1053 |
| *Branchiostoma* | *Paraspadella* | CLG | 1346 |
| *Mizuhopecten* | *Adineta* | CLG | 1224 |
| *Mizuhopecten* | *Gnathostomula* | CLG | 1053 |
| *Mizuhopecten* | *Paraspadella* | CLG | 1346 |
| *Adineta* | *Gnathostomula* | CLG | 919 |
| *Adineta* | *Paraspadella* | CLG | 1063 |
| *Gnathostomula* | *Paraspadella* | CLG | 929 |
| *Adineta* | *Gnathostomula* | Independent | 1838 |
| *Adineta* | *Paraspadella* | Independent | 2126 |
| *Gnathostomula* | *Paraspadella* | Independent | 1857 |

To establish robusticity in our macrosyntenic findings, we implemented a jackknife-approximate subsampling regime on our comparison taxa pairs. We randomly sampled pairs of orthologs from the source dataset’s ortholog pair input table to generate 100 subsampled pseudoreplicate datasets that comprise some of the original dataset’s taxa pairs, whilst randomly excluding others. For each original dataset comprised of *Branchiostoma x Mizuhopecten* CLGs, the pseudoreplicate size was equal to 50% of the original source dataset. In contrast, for each of the three datasets comprised of novel inferred linkage groups, the size of these subsampled datasets was equal to the size of the dataset inferred under the more strongly limited *Branchiostoma x Mizuhopecten* CLG sampling regime. Then the MacroSyntR analysis was rerun for each taxa pair using subsampled datasets as input data. The scripts used to accomplish this can be found at the Github associated with this manuscript (https://github.com/JFFleming/MacrosyntenicJackknife), and in our supplemental information.

For each taxa pair, the results of the 100 subsamples were summarized by counting the number of appearances of each chromosome (or chromosome/contig) pair as well as the number of times that pair’s appearance was ranked as significant by the Fisher’s exact test used by MacrosyntR to determine the linkage group. As the Fisher’s exact test is set at a significance threshold of *p=*0.05, chromosome pairs that were recovered as significant in more than 90 of the 100 subsampled datasets were deemed to be robustly significant.

## Results

### Phylogenetic Assessment: Canary Sequence Methodology

A comprehensive analysis of the LB-Score, DE-Score and nRCFV of the full, concatenated dataset (138 genes across 53 tips) identified seven taxa as potentially problematic - *Brachionus angularis*, *G. armata, P. laevis, P. laevis* (this study)*, Paraspadella gotoi, Seison nebaliae* and *S. kappae*. These taxa were all identified as either possessing an LB-Score greater than two standard deviations from the mean of the dataset, or an nRCFV greater than two standard deviations from the mean of the dataset (all sequences were tested for entropic site saturation by the DE-Score, but none were found to be saturated) (Supplemental Table 2). The sequences of these taxa were thus identified as the sequences of interest for the Canary Sequence Methodology.

Following the construction of the base dataset (46 tips across 138 genes), each sequences of each taxon was independently added to form seven checking datasets. Within these checking datasets, all seven taxa were found to be stable - the tree produced by the checking dataset was isomorphic with the base dataset, and the corresponding taxon was found in the same position in both the checking tree and the full tree. As no canary sequences were identified, all seven taxa were identified as not potentially problematic to topological reconstruction given the dataset, despite their long branches and compositional heterogeneity (Supplemental Table 2). However, this means that the sequences of three taxa that were identified as sequences of interest due to their high compositional heterogeneity could not be tested as ambiguous sequences, and so are excluded from the final tree as per methodology^40^.

Within the final 50 taxa tree constructed by the Canary Sequence Methodology, Chaetognatha represent the earliest diverging clade of Chaetognathifera, as sister to Gnathostomulia+Syndermata, whilst Acanthocephala appear as a divergent clade within Syndermata, in a position that is consistent with Pararotatoria hypothesis. Both clades are also recovered quite robustly by this dataset, with 100% bootstrap support for Acanthocephala+Seisonidea, and 99% bootstrap support for Gnathostomulida+Syndermata (Figure 2).

**Figure 2:**
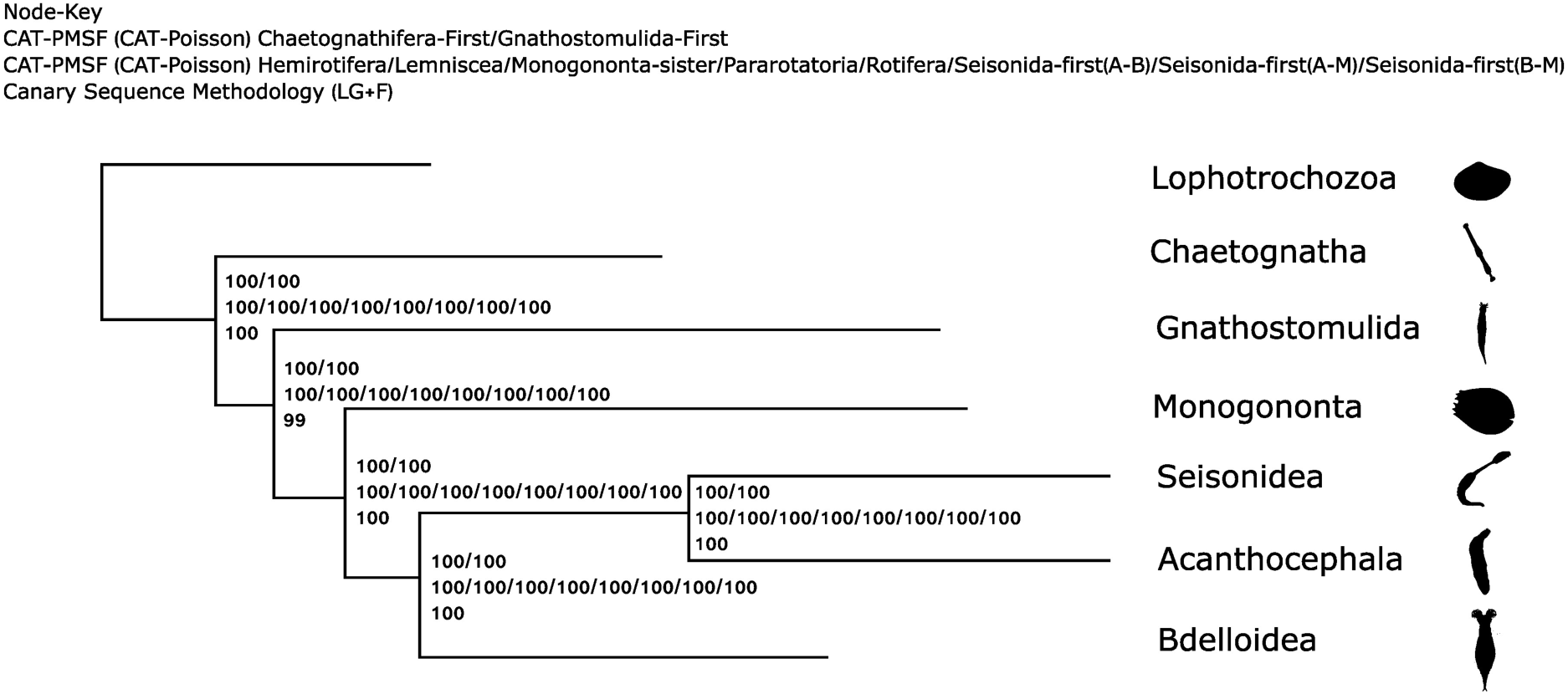
A comparative cladogram of the phylogenies generated by each of the methodologies used to assess the relationships of phyla within Chaetognathifera.

### Phylogenetic Assessment: CAT-PMSF

To further establish the reliability of our taxonomic inferences using molecular data, we used the CAT-PMSF methodology established by Giacomelli et al (2025)^61^. Here, we tested 10 conditions across the two key phylogenetic questions, obtaining the site profiles for the CAT-Poisson model first assuming Chaetognatha-first and Gnathostomulida-first, and then testing Rotifera, Pararotatoria, Hemirotifera, Monogononta-sister, Lemniscea and each of the three Seisonidea hypotheses (Figure 1).

We found that, under site profiles generated under both Chaetognatha-first and Gnathostomulida-first fixed topologies, that Chaetognatha-first was robustly recovered in both scenarios. Under the CAT-PMSF model constructed under Chaetognatha-first conditions, the Chaetognatha first topology was favoured with a bootstrap score of 100 at both the node distinguishing Chaetognatha from Gnathostomulida+Syndermata and the node distinguishing Gnathostomulida from Syndermata. Meanwhile, in the alternative scenario, the same topology was recovered, with the same support at both nodes (Figure 2, Supplemental Figure 6-7).

For our analyses testing the plausible position of Acanthocephala within Syndermata, thereby, we elected to fix our topologies under a Chaetognatha-first scenario, varying only the position of Acanthocephala with respect to the three classes of “Rotifera”. Of the eight scenarios used to generate new site profiles, all eight ultimately recovered Pararotatoria under our final analyses, with bootstrap replicates recovering overwhelmingly robust support, with 100 ultrafast bootstrap support regardless of the topology under which the site profile was generated (Figure 2, Supplemental Figures 6-14). This was even true of the three scenarios testing the plausible position of Seisonidea as the sister group to the remaining Syndermata, where all three recovered Pararotatoria (and Chaetognatha-first) with 100 ultrafast bootstrap support regardless of the generating topology.

### Phylogenetic Assessment: Gene Tree Concordance (DiscoVista)

Our gene tree concordance analyses show a clear preference for the Chaetognatha-first hypothesis in comparison to Gnathostomulida-first. The Chaetognatha-first is strongly supported in 36.86% of the gene trees in which it appears, as opposed to the 7.14% of the gene trees that show strong support for Gnathostomulida-first. Moreover, the Gnathostomulida-first topology is strongly rejected by 64.86% of gene trees in which its component clades appear, in comparison to the strong rejection of Chaetognatha-first in only 29.43% of constituent trees. When the dataset is limited to only the 138 best performing genes according to our analysis of methodological incongruence, this trend remains, with 28.92% of these genes showing strong support for Chaetognatha-first, and 26.51% strongly rejecting the hypothesis in comparison to only 8.43% showing support for Gnathostomulida-first, with 56.62% strongly rejecting the hypothesis (Figure 3).

**Figure 3:**
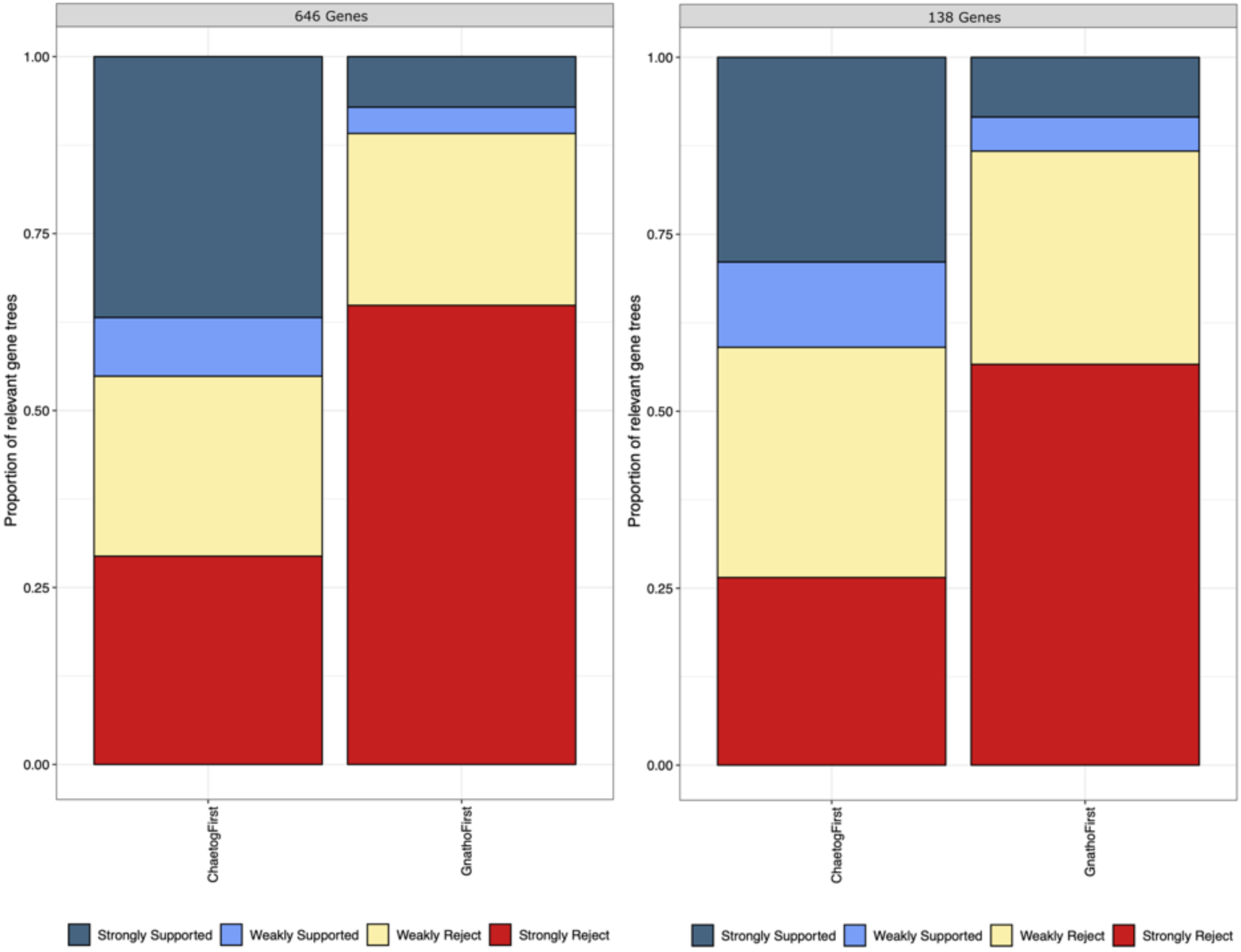
A DiscoVista plot showing gene tree support for Chaetognatha-First and Gnathostomulida-First hypotheses.

For our assessment of Syndermata, however, the situation appears more complex. Across the complete dataset, Lemniscea is mildly favoured, with 23.47% strongly supporting the hypothesis, and 53.4% of trees strongly rejecting it (remember that Lemniscea is here strictly defined as Acanthocephala+Bdelloidea). Second-favoured is Pararotatoria, with 21.43% of trees strongly supporting the hypothesis, and 59.86% strongly rejecting it. The other hypotheses show minimal amounts of support - Acanthocephala+Monogononta is supported by 6.12% of trees, Bdelloidea+Monogononta by 12.93%, Monogononta+Seisonidea by 8.84% and Hemirotifera by 4.75% (remember that Hemirotifera is strictly applied as Seisonidea+Bdelloidea). When the dataset is limited to only the 138 best performing genes according to our analysis of methodological incongruence, however, Pararotatoria becomes mildly favoured over Lemniscea, with 27.14% of gene trees in strong support of Pararotatoria, as opposed to 22.86% in strong support of Lemniscea. The same percentage of gene trees - 54.29% - strongly reject both hypotheses (Figure 4).

**Figure 4:**
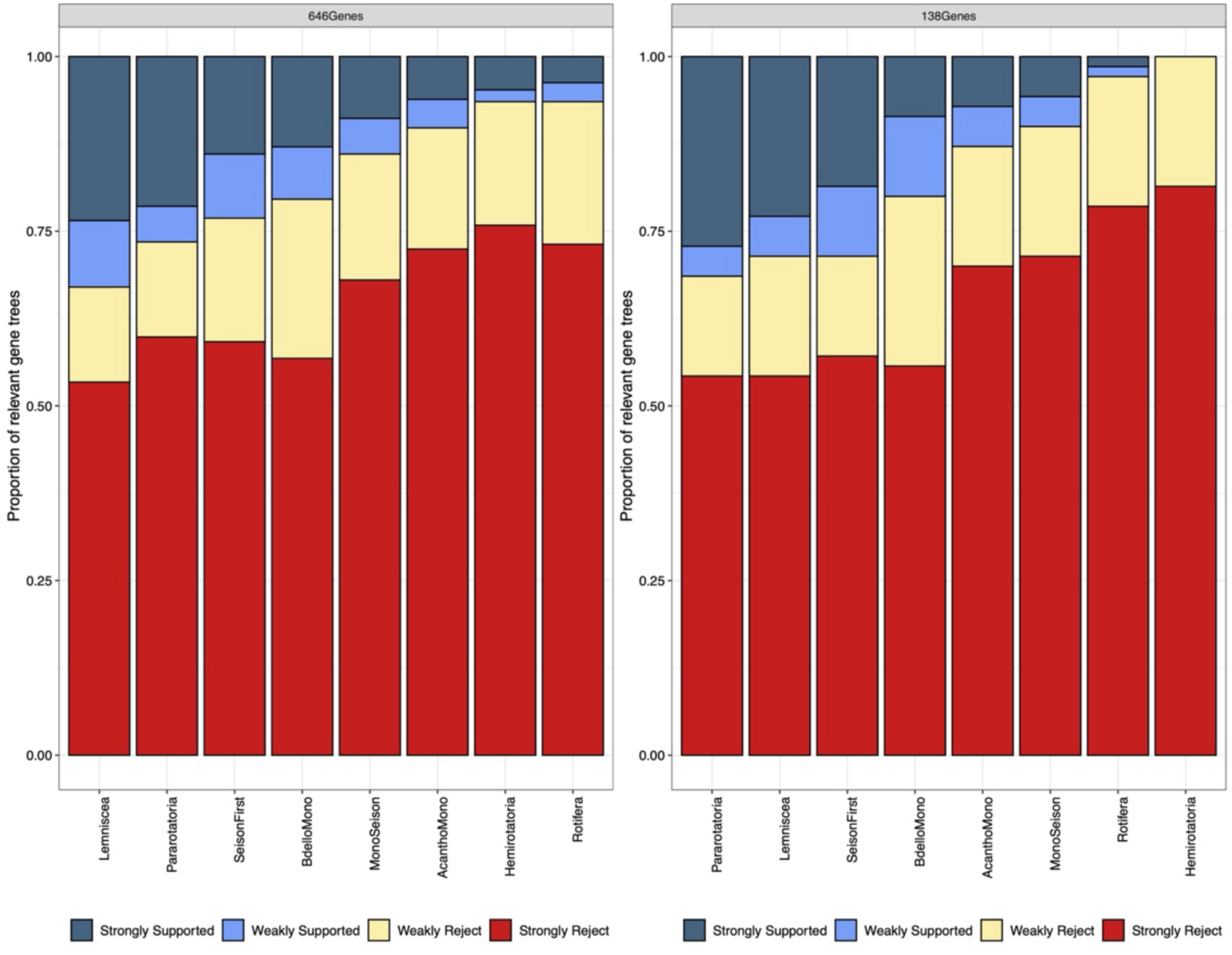
A DiscoVista plot showing gene tree support for the eight Rotifera hypotheses.

For the two hypotheses attempting to assess whether Seisonidea or Acanthocephala represent a sister taxon to the remaining Syndermata, Rotifera (Acanthocephala-first) is strongly supported by only 3.73% and strongly rejected by 73.13%, while Seisonidea-first is strongly supported by only 13.95% and strongly rejected by 59.18%. Under the stricter 138 best performing genes, strong support for Rotifera (Acanthocephala-first) increases to 14.28%, and for Seisonidea-first, strong support increases to 18.57%. While Seisonidea-first is the third-most favoured hypothesis overall of the eight constraints tested under DiscoVista, neither it nor Rotifera+Acanthocephala ever become hypotheses that were favoured over Lemniscea (Acanthocephala+Bdelloidea) or Pararotatoria (Acanthocephala+Seisonidea) (Figure 4).

### Phylogenetic Assessment: Macrosyntenic Analysis and “The Macrosyntenic Jackknife”

Our initial macrosyntenic analyses found no ALGs linking all five test species (*Branchiostoma floridae*, *Mizuhopecten yessoensis*, *Adineta vaga*, *Gnathostomula armata*, and *Paraspadella gotoi*) when using the conserved metazoan ancestral linkage groups derived from *Branchiostoma* and *Mizuhopecten*^43^. However, between smaller scale comparisons, some linkage groups could be located (Table 2). As expected, there are 23 significant chromosome pairs in our control comparison of *Branchiostoma* and *Mizuhopecten* reflecting the metazoan ALGs (Supplemental Figure 15). In our pairwise comparisons including Chaetognathifera species, significant macrosyntenic similarities could be found between *Branchiostoma* and both *Paraspadella* and *Gnathostomula* (Supplemental Figure 15), and between *Mizuhopecten* and both *Paraspadella* and *Gnathostomula*, though no significant similarities could be found in the Oxford dot plots between *Adineta* and any of the test species (Supplemental Figure 15). With respect to metazoan ALGs, given the significant pairs, the following ALGs can tentatively assigned. For *Paraspadella*, ALG D is assigned to chromosome 4 in comparisons to both *Branchiostoma* and *Mizuhopecten*. Additionally, B2 (*Branchiostoma*) to chromosome 4, B3 (*Branchiostoma*) and R (*Mizuhopecten*) to chromosome 2 and I (*Mizuhopecten*) to chromosome 1. For *Gnathostomula*, ALG D is assigned to contig 3 and L to contig 19 in both comparisons.

**Table 2:** Number of Significant Hits recovered from Oxford dot plot computations of macrosyntenic linkages, inferred from the shared metazoan ALGs.

| Test Species | *Branchiostoma* | *Mizuhopecten* | *Paraspadella* | *Gnathostomula* | *Adineta* |
| --- | --- | --- | --- | --- | --- |
| *Branchiostoma* | N/A | 23 | 3 | 2 | 0 |
| *Mizuhopecten* | 23 | N/A | 3 | 4 | 0 |
| *Paraspadella* | 3 | 3 | N/A | 0 | 0 |
| *Gnathostomula* | 2 | 4 | 0 | N/A | 0 |
| *Adineta* | 0 | 0 | 0 | 0 | N/A |

When compared to *Mizuhopecten*, J2 is assigned to contig 19, R to contig 12 and A1 to contig 36. However, while both D and R were assigned in comparisons of both *Paraspadella* and *Gnathostomula* to *Mizuhopecten*, their corresponding pairs (chromosome 4 & contig 3 and chromosome 2 & contig 12) are not significant when they are compared to one another.

Moreover, no new linkage groups could be detected between the chaetognathiferan taxa, which is in contrast to the situation in some other lophotrochozoan taxa with a breakdown of the metazoan ALGS; for example, in Clitellata and Hirudinea^31,34,36,37,64,65^.

However, we also followed up on this with a *de novo* approach using rbhXpress^43^ to directly infer ortholog pairs, and then use this information to obtain possible linkage groups specific to Chaetognathifera. Interestingly, this revealed numerous significant pairs from *Adineta* to *Gnathostomula* and to *Paraspadella*, but not a single significant pair between *Gnathostomula* and *Paraspadella* (Table 3, Supplemental Figure 16). Notably, in the case of the *Adineta*-*Paraspadella* comparison only one chromosome from each species is not assigned to a linkage group (Supplemental Figure 16). It is surprising that such apparently strong associations are not present in the “*Branchiostoma* and *Mizuhopecten”* subset, but can be so extensively discovered in the independent comparison. Additionally, while *Adineta* shares linkage groups with both, these linkage groups are not recovered when comparing *Gnathostomula* to *Paraspadella*. For example, *Adineta* CP075496.1 is in a linkage group with both *Paraspadella* chromosome 5 and *Gnathostomula* contig 19, but while the pair *Paraspadella* chromosome 5 and *Gnathostomula* contig 19 is present and shares many genes, it is not considered to be significant (Supplemental Figure 16).

**Table 3:**
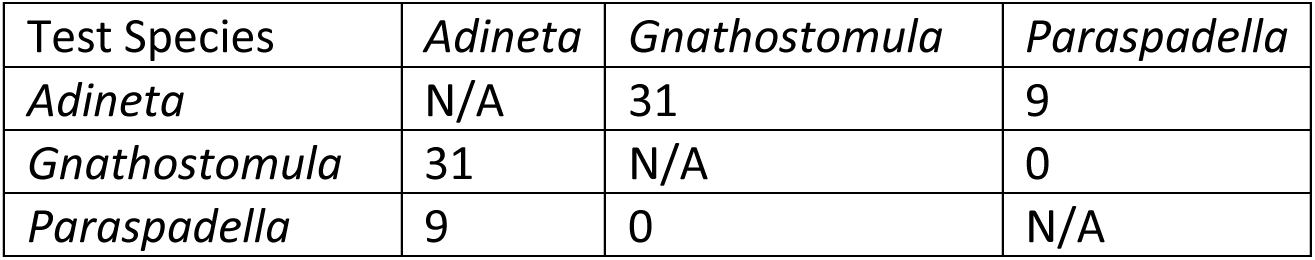
Number of significant hits recovered from Oxford dot plot computations of macrosyntenic linkages, inferred from the chaetognathiferan linkage groups.

| Test Species | *Adineta* | *Gnathostomula* | *Paraspadella* |
| --- | --- | --- | --- |
| *Adineta* | N/A | 31 | 9 |
| *Gnathostomula* | 31 | N/A | 0 |
| *Paraspadella* | 9 | 0 | N/A |

When one compares the significant pairs to other pairs for the same chromosomes, which are not significant, then it is not apparent that the number of genes is substantially different. Moreover, all chromosomes share a substantial number of genes in these three comparisons with other individual chromosomes in a non-significant way. This is substantially different from the situation found in the comparison of *Branchiostoma* and *Mizuhopecten* (Supplemental Figure 15). These substantial differences can also be seen when one compares the plots of gene positions on the chromosomes to each other (Figure 5). For example, in the comparison of *Branchiostoma* and *Mizuhopecten*, clusters of shared genes are easily recognizable, while no substantial difference can be seen in comparisons of *Adineta* to *Paraspadella*, independent of the subset of genes used. In this view, the latter two comparisons appear more like those which lack any linkage groups. Hence, despite the significant results it seems dubious that these results are robust in showing ancestral linkage groups.

**Figure 5:**
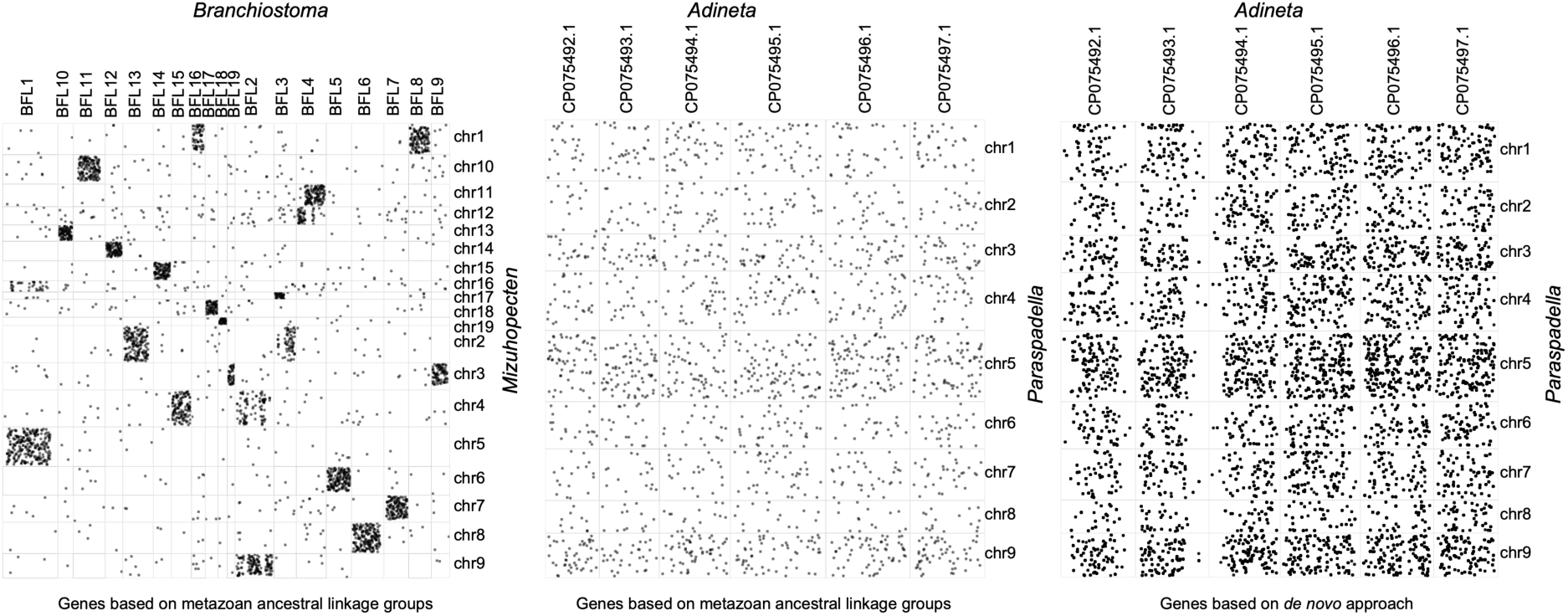
The distribution of genes based on ancestral linkage groups between *Branchiostoma* and *Mizuhopecten* (based on the conserved metazoan ancestral linkage groups), *Adineta* and *Paraspadella* (based on the conserved metazoan ancestral linkage groups) and *Adineta* and *Paraspadella* (based on the independent inferences between the two genomes). Note the more even distribution of genes across “linkage groups” in *Adineta* and *Paraspadella* in comparison to *Branchiostoma* and *Mizuhopecten*.

To establish the robusticity of these macrosyntenic inferences, we adopted a subsampling approach known as jackknifing. For each of our 13 datasets, we established the number of pairs that had been recovered as significant at least once in the 100 subsamples of each dataset, and then the number of pairs that were recovered as significant in 90% or more of the subsample datasets (and so might be considered robust inferences). While the ALGs inferred between *Branchiostoma* and *Mizuhopecten* were all robustly recovered across the subsample datasets (Figure 6), all our test conditions demonstrated significant uncertainty in linkage group inference for chaetognathiferan taxa, whether using the conserved metazoan linkage groups, or directly inferring linkage groups from the data (Figures 6 & 7, Tables 4 & 5). In most cases, none of the originally significant pairs were recovered robustly, with the sole exceptions being *Mizuhopecten* x *Gnathostomula* under the shared metazoan linkage group regime (with one significant pair recovered robustly, Table 4), and *Adineta* x *Gnathostomula* under the directly inferred linkage group regime (with two significant pairs recovered robustly, Table 5, Figure 7). For the *Mizuhopecten* x *Gnathostomula* comparison, chromosome 4 pairs with contig 19 and chromosome 4 harbors the metazoan ALGs Js and L (Supplemental Figure 15). For the other comparison, contig 30 pairs with CP075493.1 of *Adineta* and CP075494.1 with contig 25 of *Gnathostomula*.

**Figure 6:**
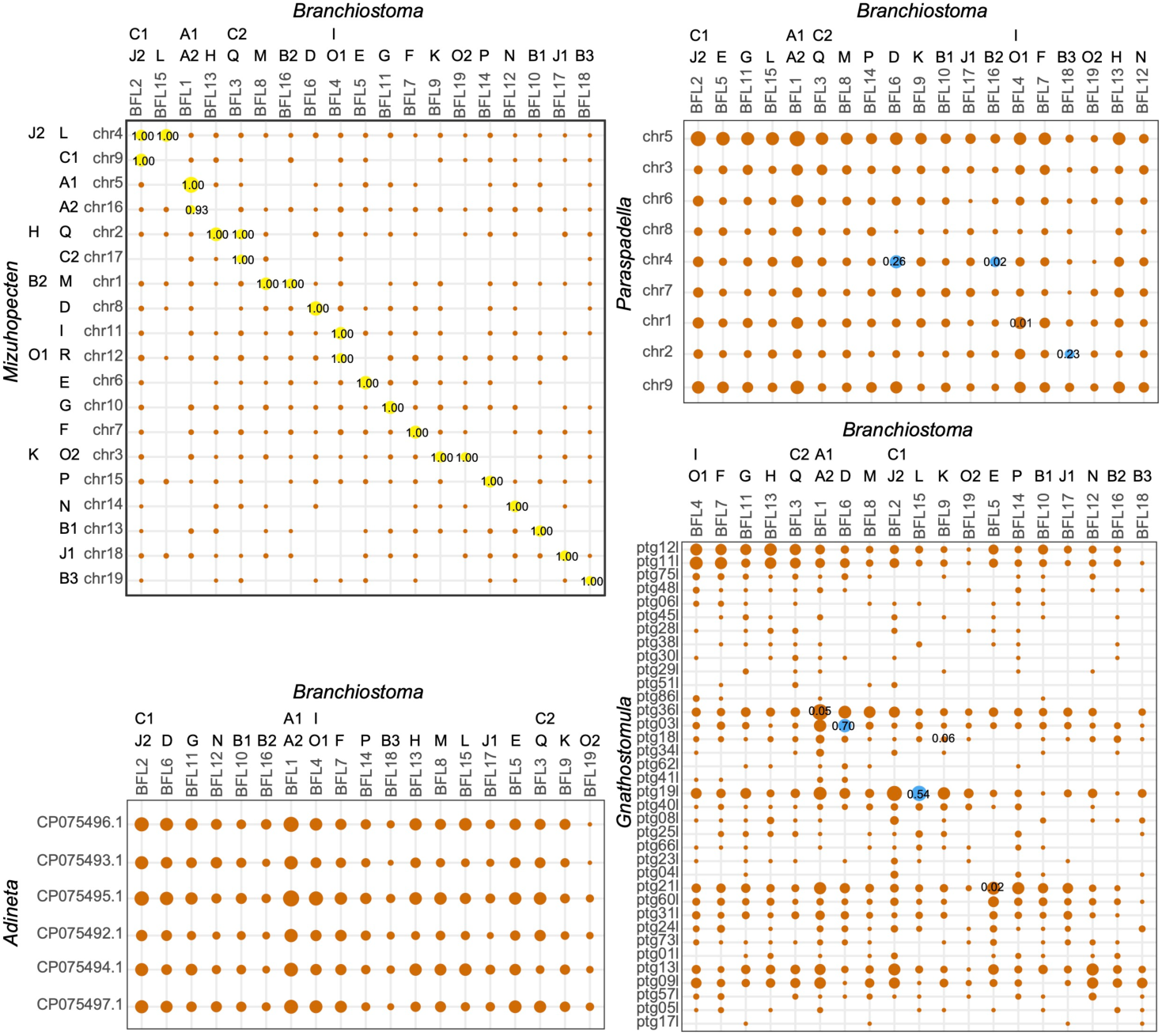
Four Oxford Dot plots with support values derived from a Jackknife subsampling approach. Each dot plot represents a comparison of *Branchiostoma floridae* with one of the four other species in the study. Yellow dots indicate a robust significant hit (a significant hit in the initial analysis that is recovered in 90% or more of subsamples). Blue dots indicate results that were recovered as significant in the initial analysis, but are not robust. Orange dots indicate results that were not recovered as significant in the initial analysis. The proportion of significant values across the 100 subsampled datasets are given when larger than 0.

**Figure 7:**
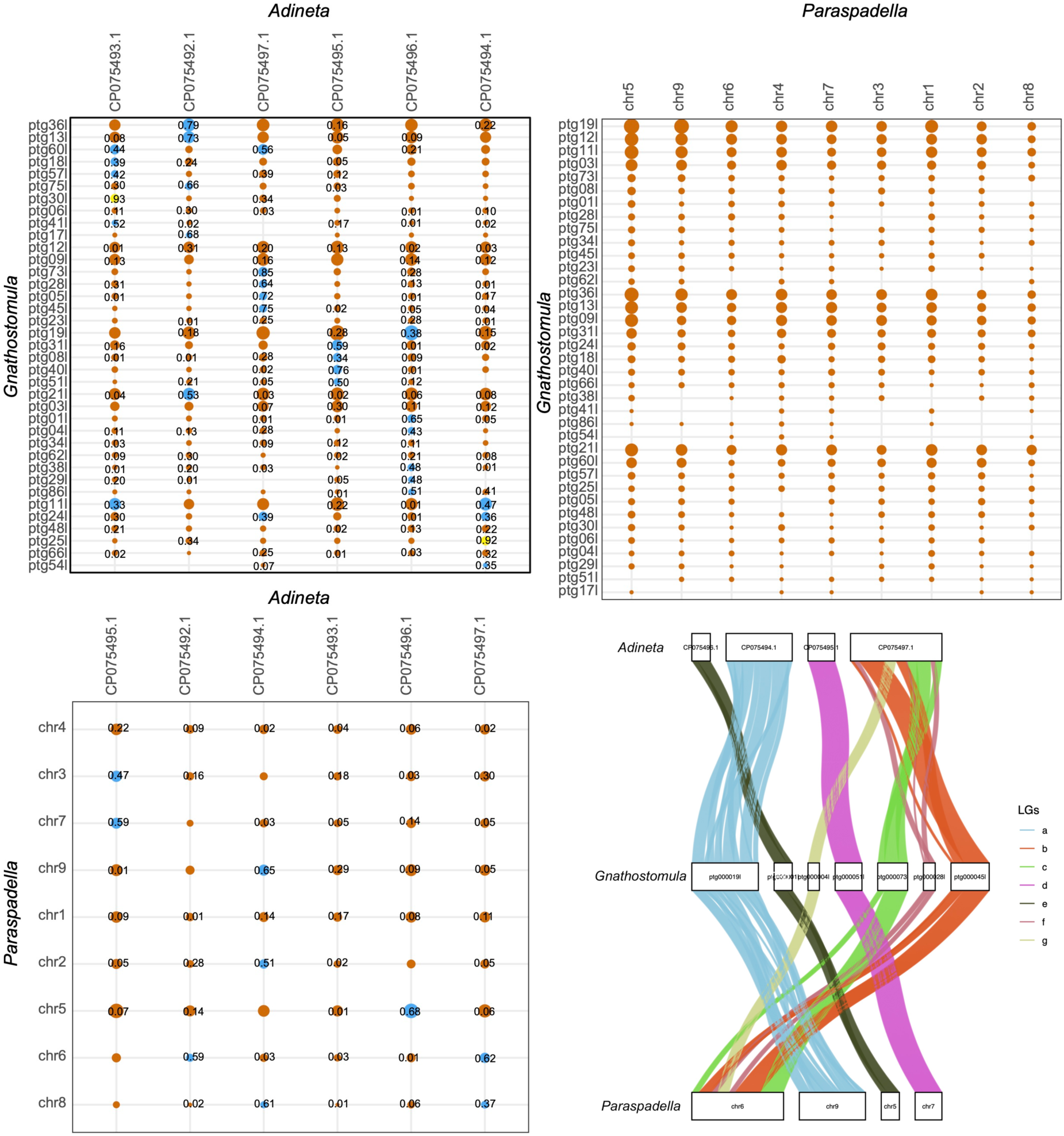
Three Oxford Dot plots with support values derived from a Jackknife subsampling approach. The bottom right panel shows the chord diagram derived from these plots, not taking robusticity into account. Note the neatness of the chord diagram despite the discordance in the accompanying dot plots. Each dot plot represents an independent comparison of linkage groups between the three Chaetognathifera in the study. Yellow dots indicate a robust significant hit (a significant hit in the initial analysis that is recovered in 90% or more of subsamples). Blue dots indicate results that were recovered as significant in the initial analysis, but are not robust. Orange dots indicate results that were not recovered as significant in the initial analysis. The proportion of significant values across the 100 subsampled datasets are given when larger than 0.

**Table 4:** The results of the macrosyntenic subsampling test on each pair of taxa, where the macrosyntenic pairs were originally derived from the shared metazoan ancestral linkage groups. All Oxford Dot plots for this table and the counts per pair are given in Supplementary Material:

| Taxa 1 | Taxa 2 | # Input Ortholog Pairs | # Significant Pairs in original dataset | # Robust Significant Pairs in subsampled datasets |
| --- | --- | --- | --- | --- |
| *Branchiostoma* | *Mizuhopecten* | 1807 | 23 | 23 |
| *Branchiostoma* | *Paraspadella* | 1346 | 3 | 0 |
| *Branchiostoma* | *Gnathostomula* | 1053 | 2 | 0 |
| *Branchiostoma* | *Adineta* | 1224 | 0 | 0 |
| *Mizuhopecten* | *Paraspadella* | 1346 | 3 | 0 |
| *Mizuhopecten* | *Gnathostomula* | 1053 | 4 | 1 |
| *Mizuhopecten* | *Adineta* | 1224 | 0 | 0 |
| *Gnathostomula* | *Paraspadella* | 929 | 0 | 0 |
| *Adineta* | *Paraspadella* | 1063 | 0 | 0 |
| *Adineta* | *Gnathostomula* | 919 | 0 | 0 |

**Table 5:** The results of the macrosyntenic subsampling test on each pair of taxa, where the macrosyntenic pairs were independently derived from the compared genomes. All Oxford Dot plots for this table and the counts per pair are given in Supplementary Material:

| Taxa 1 | Taxa 2 | # Input Ortholog Pairs | # Significant Pairs in original dataset | # Robust Significant Pairs in subsampled datasets |
| --- | --- | --- | --- | --- |
| *Adineta* | *Gnathostomula* | 1838 | 31 | 2 |
| *Adineta* | *Paraspadella* | 2126 | 9 | 0 |
| *Gnathostomula* | *Paraspadella* | 1857 | 0 | 0 |

Summarizing these results, our macrosynteny analyses show that there are very few - or zero - metazoan ALGs that could possibly be assigned to a chaetognathiferan genome. When considering only the significance level, five metazoan ALGs were recovered from *Paraspadella* (B2, B3, D, R and I), and *Gnathostomula* (A1, D, J2, L, R). While two of these linkage groups (D and R) were recovered from both species, the only robust inference was found between *Gnathostomula* x *Mizuhopecten*’s contig 19 x chromosome 4. This represents the Ancestral Linkage Groups L and J2, which were both found only in *Gnathostomula*. When we attempted to independently infer linkage groups specific for Chaetognathifera we found several possible linkage groups between Syndermata and either Chaetognatha or Gnathostomulida when considering only significance levels. However, with the exception of two linkage groups, all others were not robust in the jackknifing approach. Moreover, a closer examination of the Oxford dot plots shows that the genes of each contig or chromosome of the chaetognathiferan genome were relatively evenly distributed across the chromosomes or contigs of the other chaetognathiferan and outgroup genomes examined. In contrast, all 23 linkage groups found in the comparison of *Branchiostoma* and *Mizuhopecten* were both significant and robust using our new jackknifing approach, and the Oxford Dot plots show clear aggregations in chromosome pairs that are easily visible.

## Discussion

### Macrosynteny in deep time? The need for caution and thorough testing of robusticity

Our macrosyntenic results very strongly evidence that no metazoan ALGs are significantly conserved in the genomes of Chaetognatha, Gnathostomulida and Syndermata, with the sole exception of maybe J2 and/or L in Gnathostomulida. The same is the case for possible chaetognathiferan linkage groups, of which none were determined to be significantly present in the study. The findings herein are in agreement with previous results in Syndermata, but in contrast to those in Chaetognatha, that detected 20 out of the 24 metazoan ALGs confined uniquely to specific chromosomes^39^. However, this conclusion has already been questioned, as the Oxford Dot plots show a more or less even distribution of gene pairs across the entire grid - as can also be observed in this study (Figure 5)^18^. The distribution of the genes in the Oxford Dots plots for chaetognathiferan taxa strongly resembles the distributions that can be seen when comparing a clitellate taxon to a non-clitellate one or a hirudinean taxon to a non-hirudinean one^31,36,37^ - and both of these examples actually show stronger aggregations of genes than plots with chaetognathiferan taxa. The situation in Clitellata and Hirudinea has also been called “extensive genome-wide scrambling”^37^. Similarly, for Chaetognatha, Gnathostomulida and Clitellata, it must be concluded that each of these genomes have independently experienced massive chromosomal rearrangements and a genome-wide scrambling. For example, massive rearrangements of chromosomes have also been reported for bryozoans, coleoid cephalopods, platyhelminths, vertebrates, lepidopterans and tunicates^34,36,38,64–70^.

The significant linkage groups detected in this study, and in other analyses of Chaetognatha, could possibly be due to an overestimation of significance^71^, with only minimal fractional differences in the number of genes resulting in inferences of significant difference^18^. This problem is generally known as the large (N) problem in statistics concerning biological studies using large amounts of input data^72–74^. In this case, increasing N will increase the test power and decrease p-values “due to the inherent trade-off between type I and type II error probabilities in frequentist testing. This trade-off raises concerns about the reliability of declaring ‘statistical significance’ based on conventional significance levels”^72^. In essence, it means that minor differences between two populations of values can be statistically significant given standard significance levels, but meaningless to answer the question at hand.

In the case of macrosynteny, for example, the genes of an ALG could be relatively evenly distributed amongst the majority of chromosomes in a comparison of species, while being slightly more densely arranged in one of the chromosomes. This slight increase could result in an inference of significant difference, but in fact only indicate that there are significantly more genes of the ALG found in this chromosome than in the other chromosomes, not that genes belonging to this ALG do not reside elsewhere in the genome, or are found in only small numbers. Accordingly, it does not naturally follow that this slightly higher aggregation of genes in one pair of chromosomes is a result of shared ancestry, in which this linkage group was passed through the ages more or less unchanged. It could also be a chance accumulation of genes, or the result of a smaller subset of genes that are stable through time. In both cases, inferring that these chromosomes are homologous would be erroneous.

A basic assumption of the macrosyntenic approach in phylogenetic studies is that the ALGs are homologous units that can be reliably traced through time, and speciation. Due to this, fusion events coupled with mixing can indicate relatedness as they are comparable to Shannon entropy^33^. However, for this assumption to be reliable, the ALGs must be robustly established, with a large majority of the genes in the ALG confined to only the indicated chromosomes, and not several other non-indicated ones as well. If the other chromosomes harbor a larger number of genes of the ALG, this could be due to either translocations of small numbers of genes (or even single genes) from one chromosome to another, or the trace of a prior fusion event that was followed by fission events. The likelihood of the latter increases tremendously with the number of genes found outside the indicated chromosomes that belong to that ALG. Accordingly, these chromosomes would have to be counted as members of the ALG as well.

Given the problem of spurious or random statistical significance, it is important to establish that the detected linkage groups are not only statistically significant, but that the indicated linkage groups comprise the clear absolute majority of the genes of this linkage group. It has already been suggested that a more conservative approach should be taken to the assignment of ALGs to minimise the risk of calling false positives^71,75^. Herein, we show a subsampling strategy also known as jackknifing. Jackknifing procedures have a long tradition in phylogenetics, where they are used to assess the robustness of the support for a node in a reconstructed tree (though a related technique, bootstrapping, is more popular)^76,77^. The rationale is that a strong and robust signal should still be detectable and recoverable in a smaller random subset of the original sample. The higher the frequency of recovering the same result, the more robustly the original result can be regarded. In the context of macrosynteny, this result would be the detection of a significant pair.

Our subsampling procedure herein clearly evidenced the issues outlined above - when little similarity can be found between genomes at the chromosome scale, due to the high number of comparative tests made between orthologous groups, significance can be inferred due to random chance or minimal differences that are meaningless in the context of macrosynteny. Our subsampled comparison of *Mizuhopecten yessoensis* and *Branchiostoma floridae* (using the dataset provided as an example packaged with MacroSyntR) showed reasonably robust inference of ancestral linkage groups, with 22 recovered as significant across all 100 subsamples, and one recovered in 93/100, and our comparison of *Adineta vaga* and *Branchiostoma floridae* showed no linkage groups recovered as significant at all. However, our comparisons with other chaetognathiferan genomes were significantly different.

*Branchiostoma* x *Gnathostomula* showed a recovery of five linkage groups as significant across the 100 subsamples, but with a robusticity ranging from 2-70%, while *Branchiostoma* x *Paraspadella* recovered four linkage groups as significant with robusticity ranging from 1-26%. None of these inferences can be reasonably considered robust - assuming a normal distribution around the 5% significance threshold of Fisher’s exact test, to account for significance not being found by random chance, a robust significant hit is one that can be found in 90% or more of subsampled datasets.

When considering the evolution of Chaetognathifera through current macrosyntenic methods, our results are strikingly inconclusive. Between *Adineta* and *Gnathostmula*, two linkage groups are robustly recovered (with subsampled percentages of 93% and 92% respectively), whilst a further 138 pairs were recovered with lower robusticity, varying from 1%-85%. In this case, our lack of a truly chromosome-scale complete genome will inevitably lead to more (and less robust) inferences, but this trend is also carried to our comparison of *Adineta* and *Paraspadella*, two chromosome-scale genomes within the superphylum. Here, no linkage groups were robustly recovered, and 47 were recovered across the 100 subsampled datasets with robusticity ranging from 1%-68%. In our final comparison of *Gnathostomula* and *Paraspadella*, not a single ancestral linkage group was recovered across any of the subsamples. This again suggests that significant chromosome rearrangement has likely occurred multiple times within the clade. Whilst we may be able to tentatively say that the chromosome macrosynteny of *Adineta* and *Gnathostomula* appears more similar than either are to *Paraspadella* - thus further reinforcing support for the Chaetognatha-first hypothesis - this support is qualitative and not as robust as the evidences produced by more classical molecular phylogenetic methods, such as gene tree concordance or our CAT-PMSF assessments. Establishing the nature and timing of these chromosomal rearrangements will require significantly more chromosome-scale genomes from all the chaetognathiferan phyla (including Micrognathozoa) but could help us understand much more about the tempo and mode of macrosyntenic evolution.

Furthermore, linkage groups with low robusticity from our subsampling procedure can be misleading. Especially considering the growing popularity of macrosyntenic analysis, this discovery reinforces the concept that careful observation of Oxford dot plots is required to engage in any serious inferences about macrosyntenic evolution, to ensure that significant similarities found between genomes are truly robust. Accordingly, such plots should always be reported as part of macrosyntenic studies, and not just the ribbon plots generated from them, which can be misleading due to their reductive behavior. For example, our analysis to detect possible chaetognathiferan linkage groups generated a very conclusive and seemingly strong ribbon plot comprising at least some of the chromosomes and contigs (Figure 7, Supplemental Figure 17). However, it depicts direct connections between *Paraspadella* and *Gnathostomula* even though not a single significant pair was found in the pairwise comparison (Figure 7).

Moreover, it suppresses all the gene-gene connections that do not belong to the recognized linkage group, providing an impression of certainty and uniqueness that is not present. Finally, subsampling represents a quick way to actively test the robusticity of these inferences and numerically assign confidence to our understanding of chromosomal change over time. The scripts we used to accomplish this are available on the Github associated with this project, and are freely compatible with MacroSyntR^43^.

### The evolution of Chaetognathifera

Our phylogeny clearly demonstrates that Chaetognatha mark the earliest diverging group within Chaetognathifera, followed by Gnathostomulida. Across all our analyses, Gnathostomulida was consistently recovered as the sister group to Syndermata, with incredibly strong support. Notably, this support was not strongly affected by selecting genes within our gene tree discordance analysis, suggesting that the phylogenetic signal supporting this topology is particularly robust. Moreover, when selecting genes with low methodological incongruence, the relative support for this hypothesis only increases, suggesting quite strongly that Gnathostomula-first is an artifact driven by compositional and branch length heterogeneity^18,19,22^. This also means that the monophyly of a Gnathifera comprising Gnathostomulida, Micrognathozoa and Syndermata is supported. Although not included in our analysis, Micrognathozoa are consistently recovered as sister to Syndermata in recent phylogenomic studies^18^ and morphological data also supports this close relationship^11,12,20,21^.

Hence, the synapomorphy of the jaws found in these three phyla is supported. The jaw is predominantly used as a rasping organ in Gnathostomulida, Micrognathozoa and in many syndermatan species. It most likely emerged as a rasping organ for grazing (on, for example, bacterial mats) and only later evolved in adaptation to other dietary strategies such as predation. Moreover, all three phyla are ancestrally meiofaunal or interstitial benthic species of small body sizes. Hence, the gnathiferan ancestor was also most likely a meiofaunal or interstitial benthic species.

On the other hand, Chaetognatha are usually not considered as meiofaunal or interstitial but are larger in body size - with individuals in the cm-range who live as pelagic predators using spines protected by a hood at the head for grasping prey. Due to their sister-group relationship, it cannot be concluded with certainty that the pelagic lifestyle of larger Chaetognatha evolved from a smaller benthic ancestor or vice versa, and so the ancestral ecology of the shared ancestor of all Chaetognatha cannot be immediately deduced. As the relationships of Platytrochozoa – the sister-group of Chaetognathifera – are still unresolved^18^, the outgroup also cannot be used to contextualise the common ancestor. Another topic of consideration is whether the grasping spines of Chaetognatha and the gnathiferan jaws are truly homologous. Support in favor of a homology stems from their shared “composite organization at the ultrastructural level with alternating layers of material that is opaque or dense to electrons disposed in tubular fashion”^3^. Additionally, the fossil species *Amiskwia sagittiformis* appears to possess an intermediate morphology between Gnathifera and Chaetognatha by the possession of both a paired pharyngeal jaw and a general body organization resembling chaetognaths, including a hooded head^6,78,79^. Hence, given all evidence, it seems likely that the chaetognath spines evolved from a gnathiferan-like jaw. Due to this, while it does not strictly follow that the shared ancestor also possessed a gnathiferan-like lifestyle, it seems more likely than a chaetognathan-like ecology.

Within Syndermata, Monogononta represent the earliest diverging group, followed by Bdelloidea, and then Acanthocephala+Seisonidea. Notably, our gene tree discordance analysis of Syndermata, combined with the results of our CAT-PMSF analysis, suggests that support for the Lemniscea hypothesis is likely driven by compositional biases and branch length heterogeneity within the dataset (Figures 2-4). We found no significant evidence of entropic site saturation across the 646 genes, but significant variation in compositional and branch length heterogeneity - using models such as CAT-PMSF that are better able to account for this variation^41^, or simply removing them from the dataset as we did in our selective gene tree analyses, appears to strongly increase support for Pararotatoria over Lemniscea. This further support for the Pararotatoria hypothesis across all our analyses is consistent with more recent molecular studies, and presents a growing consensus from multiple data sources^25–27,29^, and multiple analytical methods, that this may represent a more reliable, modern understanding of the evolution of this clade. This is also notably consistent with some observations of both the ecology and morphology of these clades - Seisonidea comprises a number of epibiotic species, and it is plausible that the shared ancestor of Seisonidea and Acanthocephala possessed a similar ecology prior to specialising as an endoparasite^24,28,29^. Both Acanthocephala and Seisonidea lack a corona, and both have modified their proximal appendages for latching onto a host organism, further corroborating this molecular argument^8^. As a consequence, the lemnisci and epidermal intrusions in acanthocephalans and bdelloids, respectively, are either not homologous structures, and represent two independent evolutions of the ability to retract the anterior end of the organism^80^, or represent a homologous structure that has been independently lost in Seisonidea due to the organization of the neck in pseudosegments.

### Strong amounts of change, without saturation

During this analysis, the Dayhoff Exchange Score (DE-Score) was used to assess the entropic saturation of the 646 genes that comprised the dataset^56^. Notably, not a single gene was found to fall below the critical DE-Score for that gene. Despite this, several genes were susceptible to long branch attraction artifacts and compositional heterogeneity, two other violations of the SRH assumptions, which were measured using nRCFV and the LB-Score^57,58^. Thanks to new ways to assess entropic saturation without bias driven by these other forms of incongruence, we can now more clearly consider the model insufficiencies driving not just controversies and incongruences within Chaetognathifera, but more broadly within Lophotrochozoa-Spiralia.

This same pattern was also visible in an earlier reassessment of the Kocot et al (2017) all-Spiralia dataset^2,56^. This lack of entropic saturation, especially in the presence of high amounts of branch length heterogeneity and compositional heterogeneity in both datasets suggests that Lophotrochozoa-Spiralia as a superphylum more generally is one dominated by directed change, or unsampled change - this could indicate rapid diversifications in the common shared ancestor of Lophotrochozoa-Spiralia, or that the modern extant phyla represent a depauperation of a much greater extinct diversity^18,19^. The latter inference is hard to test - reservoirs of spiralian diversity may still be present in the form of cryptic species, or inhabiting challenging environments and refugia, but even then, the extinct diversity of a predominantly soft-bodied superphylum is challenging to assess or approximate^81^.

However, it remains notable that amino acid change within Chaetognathifera, and Lophotrochozoa-Spiralia as a whole, appears to be directed, rather than random - and that it is compositional signal, and a degree of directed change, that is challenging our models in this case. It represents a stark contrast from controversies in early Metazoa, which often appear to be driven by a combination of all three - and where entropic saturation appears to drive substantial topological incongruence^82–84^.

## Conclusions

The growing quantity and quality of genomic data, particularly from historically understudied and controversial phyla allows us to understand the history of the diversity of life like never before. However, it is also clear that, with this data, understanding methodological approaches, and selecting good data with representative models, will become ever more important. By carefully considering methodological incongruence in our data, and with the inclusion of novel data from these understudied clades, we can firmly resolve two controversies in invertebrate systematics. Understanding the natural history of Chaetognathifera more clearly also grants us the firm foundations to consider the relationships between other clades of Lophotrochozoa-Spiralia. Beyond this, a more complete understanding of novel genome-scale data such as this also helps inform our understanding of new methods, such as macrosyntenic analysis, leading us to methods that allow us to make more robust inferences. A thorough assessment of the support and robusticity of possible linkage groups revealed that they are lacking in the chaetognathiferan taxa and massive chromosomal rearrangements must have taken place. Most importantly, this study clearly revealed the necessity of careful examination of possible linkage groups in macrosyntenic studies beyond assessments of just the significance level. Future studies of macrosynteny should also assess robusticity in addition to significance using, for example, a jackknifing approach like herein. Only linkage groups that are both significant and robust should be considered as true linkage groups, to minimize the risk of misleading false positive results.

