## Supplemental Methods, Tables and Figures for "The phylogenetic affinities of Chaetognathifera, with considerations of systematic error and the robusticity of macrosyntenic results"

#### Genome assembly, repeat masking and annotation

First, bam files were converted to fastq files using BEDTools 2.29.2[1]. The distribution of read lengths was assessed using custom bash and R scripts. The kmer distribution was determined using Jellyfish v.2.3.0[2] up to coverage of 10.000 and 1 million and then GenomeScope2.0[3] to obtain kmer-based genome size and other parameters with default settings for a diploid genome. Ploidy levels were estimated using Smudgeplot v.0.2.4[3] with default settings and the counts from the Jellyfish analysis. The genome was assembled using hifiasm v.0.25.0[4-6] with default settings. Next, applying the BlobToolKit v. 4.4.6[7] filter option only contigs of the primary assemblies were kept fulfilling certain criteria. The selection of the criteria was guided by allowing only slight decreases in the BUSCO score. For *G. armata*, the criteria were a GC content between 0.38 and 0.41, a coverage between 130x and 300x, and a length above 500,000 bp. For *E. meneta*, contigs with a GC content between 0.24 and 0.30 and coverage between 20x and 200x were kept, while contigs matching against non-metazoan taxa except for matches against Ascomycota and all in the category "Others" were removed. For *P. laevis*, all contigs with a match against non-metazoan taxa except for the ones in category "Others" were removed as well as all that did not have a GC content between 0.28 and 0.40, a coverage between 80x and 200x, or a length above 50,000 bp. For *S. kappae*, contigs with a GC content between 0.38 and 0.50, a coverage between 50x and 400x and a length above 40,000 bp were kept, while contigs matching against non-metazoan taxa except for matches against Ascomycota and all in the category "Others" were removed.

After the filtering, Purge_Dups v.1.0.1[8] with an expected identity ≥95% and keeping secondary and supplementary alignments was applied to purge potential duplicates. Then, the assemblies were investigated again using BlobToolKit. For *E. meneta*, all remaining contigs matching a non-metazoan taxon and all in category "Others" were removed. For *P. laevis* (*sensu lato*), all remaining contigs matching a non-metazoan taxon were removed. For *S. kappae*, all contigs not matching a metazoan taxon, except for the ones matching Ascomycota, were removed as well as the ones with a length below 40,000 bp, a GC content not between 0.39 and 0.5, or a coverage not between 60x and 530x. For *G. armata*, no contigs were removed. For *G. armata* and *S. kappae*, a custom script was used for an additional test for contigs matching against non-metazoan taxa. The contigs were split into 20 equal-sized parts of at least 1,000 bp length. For *G. armata*, each part was checked using a blastn search of BLAST+ v.2.16.0[9] was conducted against a local copy of the nt database as off November 2025 and the 20 best hits kept. For *S. kappae*, a blastx search DIAMOND v.2.0.7[10] against the uniprot_swissprot database as of February 2026 was conducted. For each hit, the corresponding taxon was determined. Finally, the proportion of hits against a metazoan taxon was determined. All contigs with a proportion of less than 60% were removed from the assembly. For *G. armata*, none of the two contigs had a proportion below 60%. For *S. kappae*, 202 contigs had proportions below 60%. These were removed using seqkit v.2.12.0[11]. Finally, the same script was used to check the remaining contigs of *P. laevis* matching against Actinopteri for cross-contamination by the fish host (chub, *Leuciscus cephalus*, and European eel, *Anguilla anguillae*). The maximum proportion of hits against Actinopteri in this check was 15% for any of the contigs. Accordingly, none of these contigs showed proportions greater than 15%, only one other contig had a proportion of hits of 11.1% and all others were lower than 10%. Hence, none of these contigs were considered cross-contamination.

At different steps, quality assessments of the assemblies included QUAST v.5.2.0[12], merqury v.1.3[13], BUSCO v.5.4.7[14], and BlobToolKit v. 4.4.6[7]. The merqury analysis was based on a meryl database of the reads with a kmer size of 21. The metazoa_odb10 dataset was checked in the BUSCO analysis. Besides the BUSCO results, the BlobToolKit analysis also aggregated the coverage based on minimap2 v.2.29[15,16] and SAMtools v.1.21[17] and a taxonomic assignment. For *G. armata*, this was based on a megablast search against the local nt database, an e value of $1e^{-20}$, a maximum of 1 target sequence and a maximum of 1 high-scoring segment pair. For the other three species, this was done by blastx search with DIAMOND v.2.0.7 against the uniprot_swissprot database with the sensitive setting, a maximum target sequence of 1 and e value of $1e^{-20}$. Additionally, all mitochondrial genomes in the assembly were retrieved by using a tblastn search of the protein-coding genes of the mitochondrial genome of *Platynereis dumerlii* (NP_009239) as query against the assembly and an e-value of $1e^{-20}$. The species of the retrieved contigs were then identified using a blastn search against the local nt database and keeping only the best hit. The taxonomic path of each unique TaxonID was retrieved from the NCBI taxonomy database.

For repeat annotation and masking, first, repeat models were generated using RepeatModeler v.2.0.6[18] with two additional rounds and the structural identification of long terminal repeat (LTR) retrotransposons. Using RepeatMasker v.4.1.7[18], simple repeats are annotated and masked, followed by repeat models sourced from the Dfam database "Metazoa", and then based on the species-specific repeat models generated before. The results of the three annotation steps were combined into one and different summary data and graphics generated using tools incorporated in RepeatMasker v.4.1.7 and R v.4.2.1 in a custom script.

The annotation of protein-coding genes was conducted generally following the EBP-Nor annotation pipeline[19]. First, protein alignments were generated using miniprot v.0.13[20] based independently on three different resources. For all four species, alignments were generated based on protein models from the uniprot_swissprot database and the Spiralia database of OrthoDB_v12. For *G. armata*, protein models were retrieved from transcriptomic data of *Austrognatharia medusifera*, *Haplognathia ruberrima*, and four unidentified species of Bursovaginoidea (N. Roberts, unpublished results). For the other three publicly available resources from annotated genomes have been used (i.e., E. meneta: *Adineta ricciae* GCA_905250025.1, *A. steineri* GCA_905250065.1, *A. vaga* GCA_021613535.1, *Brachionus calyciflorus* GCA_905250105.1, *B. plicatilis* GCA_003710015.1, *Didymodactylos carnosus* GCA_905250885.1, *Rotaria magnacalcarata* GCA_965140935.1, *R. socialis* GCA_965120325.2, *R. sordida* GCA_905251635.1, *Rotaria* sp. Silwood1 GCA_905250075.1, *Rotaria* sp. Silwood2 GCA_905329745.1, *Seison nebaliae* GCA_023231475.1; *P. laevis*: *Neoechinorhynchus agilis* GCA_051530055.1, *P. laevis* GCA_012934845.2; *S. kappae*: *Paraspadella gotoi* GCA_051294375.1). At different steps, protein and mRNA sequences were extracted using agat v.1.4.2[21] and GFF3 files generated using evidencemodeler v.1.1.1[22]. Using galba v.1.0.11[23], *ab* *initio* protein-model were predicted based on the protein models specifically used for each species and setting “stopCodonExcludedFromCDS” to false. The four GFF3 files (from species-specific source data, uniprot, orthodb and galba) were combined and gene models predicted using evidencemodeler v.1.1.1. As part of this, the weight of the *ab initio* prediction by Augustus v.3.5.0[24] was 1 and of the miniprot prediction 4 and, additionally, options -m and -i specified as 10 and 1,500 bp. Next, low-quality gene predictions were removed with agat by detecting repeat-like proteins using diamond v.2.1.10[10] against the repeat database of funannotate v.1.8.17[25], an e value of $1e^{-10}$ and keeping only the best hit, by detecting proteins with "X" (e.g., translation across gaps) and by detecting protein sequences shorter than 50 amino acids. The structural annotation was followed by a functional annotation. Using InterProScan v.5.62-94.0[26], the proteins were annotated with known protein domains, GO terms, and family classifications. Additionally, another functional annotation was done by using diamond searches against the UniProt database, the sensitive setting, a maximum target sequence of 1 and e value of $1e^{-25}$. The annotations of both approaches were merged and exported as a GFF3 file using agat. Finally, summary statistics of the functionally annotated genes were generated using agat. The final set of protein and mRNA sequences was also extracted. The quality of the protein sequences was also checked using BUSCO and the metazoa_odb10 database.

### Supplemental Results

#### Genome assembly and annotation

The assembly was based on about 1.7 to 5.6 million PacBio Hifi reads with a mean read length between 6,673 to 13,144 bp and a median one between 5,873 to 12,720 bp. The vast majority of reads were recovered at around 10 kbp, with the exception of *P. laevis*, where the majority of reads were recovered at around 5 kbp (Supplemental Figure 1). However, the lengths of some reads were above 20 kbp and, in the case of *S. kappae*, even above 50 kbp. Models of kmer distribution did not converge on meaningful results for *E. meneta* and *S. kappae* (Supplemental Figure 2). For *G. armata*, it indicated a genome size of 99.6 Mbp, a uniqueness of 35.3%, a heterozygosity of 6.4% and a kmer coverage of 126x. For *P. laevis*, the numbers were 328.8 Mbp, 32.2%, 1.1% and 56x, respectively. Similarly, in the SmudgePlot analysis[3], *S. kappae* failed to converge on a reliable result, while for the other three species a diploid status is predominantly proposed (Supplemental Figure 3). However, for *G. armata*, a low level of triploidy might be present.

The sizes of the final assembled genomes after all filtering steps ranged from 91.4 Mbp in *E. meneta* to 278.3 Mbp in *P. laevis* with the ones for *G. armata* and *S. kappae* having 128.6 Mbp and 192.3 Mbp (Supplemental Table 1, Supplemental Figure 4). During the filtering steps, substantial numbers of contigs were removed from the first assembly due to either duplicated contigs (possibly due to higher levels of heterozygosity) or contamination with non-target DNA. It has been shown that whole genome amplification can also results in the predominant amplification of non-target DNA[27]. On the other hand, even some contigs of the final assembly have the best hit against a non-metazoan taxon (Supplemental Figure 5)[7]. However, these hits are short stretches (< about 2 kbp) of large contigs, and are usually weak hits. The additional check using different parts of these contigs showed that for these the majority of hits (> 60%) resolved against metazoan taxa. As a consequence of the sometimes strong filtering regime, our completeness scores are especially low for *E. meneta* and lower for *P. laevis* and *G. armata* (Supplemental Table 1). On the other hand, QV values are all above 40 as required by the EBP standard[28] and the assembly contains fewer than 2 errors per 10 kbp, or <0.002%. Similarly, there are also fewer than 2 N's per 10 kbp. The GC content ranges from 26.5% to 44.3% across the four species.

The genome of *G. armata* is highly contiguous, with just 37 contigs, an L50 of 9 and an N50 of 4.6 Mbp (Supplemental Table 1, Supplemental Figures 4-5). At 9.6 Mbp, it also has the largest contig of all four species. Hence, even though we did not apply a scaffolding method, this genome can be regarded as almost chromosome-level. The other three genomes are more discontinuous. *P. laevis* is the second best with an N50 of 533.3 kbp and L50 of 172 over a total of 724 contigs, followed by *S. kappae* with a N50 of 182, kbp and L50 of 311 (of 1,397). Both also have contigs larger than 1 Mbp. The rotifer *E. meneta* has a largest contig of about 0.5 Mbp and accordingly a N50 of 114.5 kbp with a L50 of 251 (Supplemental Table 1, Supplemental Figures 4-5).

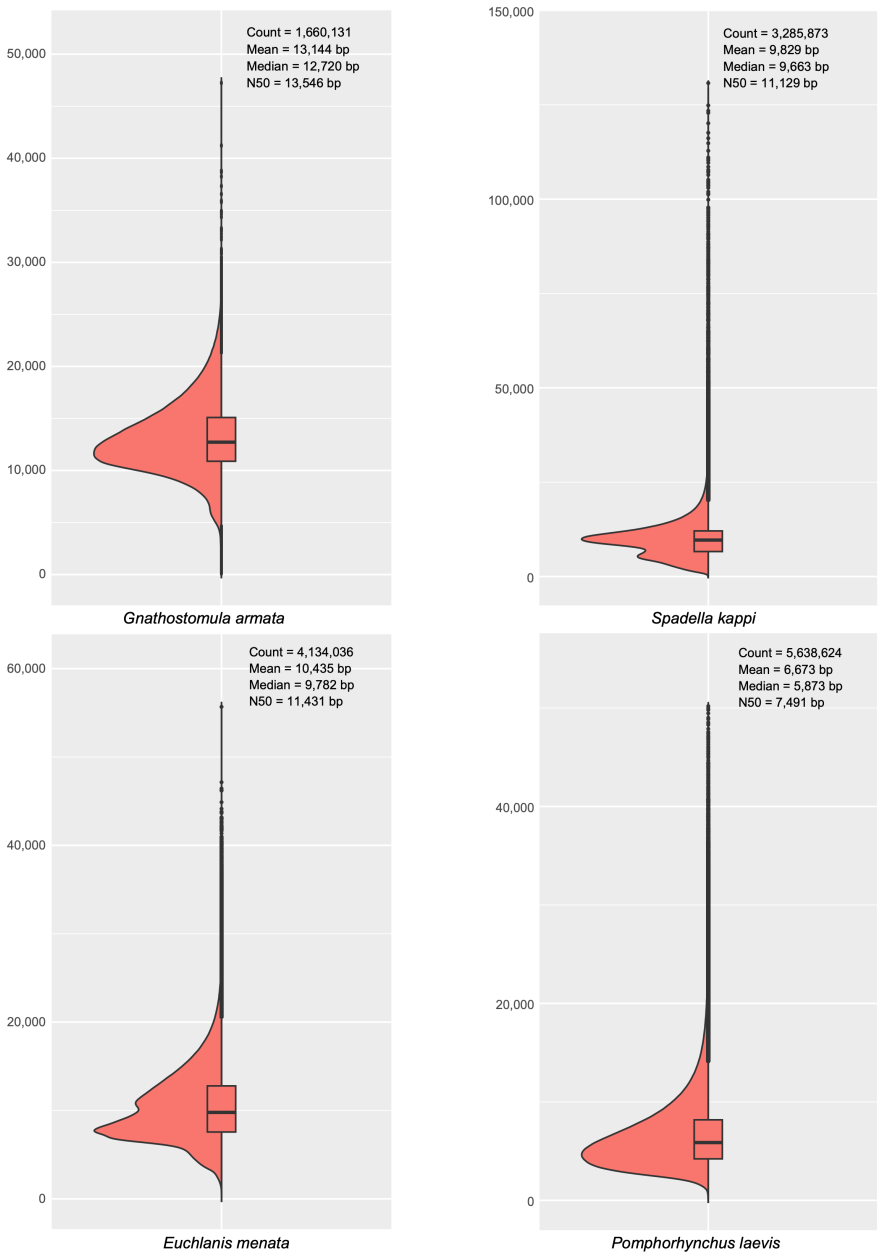


Supplemental Figure 1: Read length distribution of PacBio Hifi data for the four species shown as violin and box plots. In the upper right corner, the number of reads and the mean, median and N50 read length are shown.

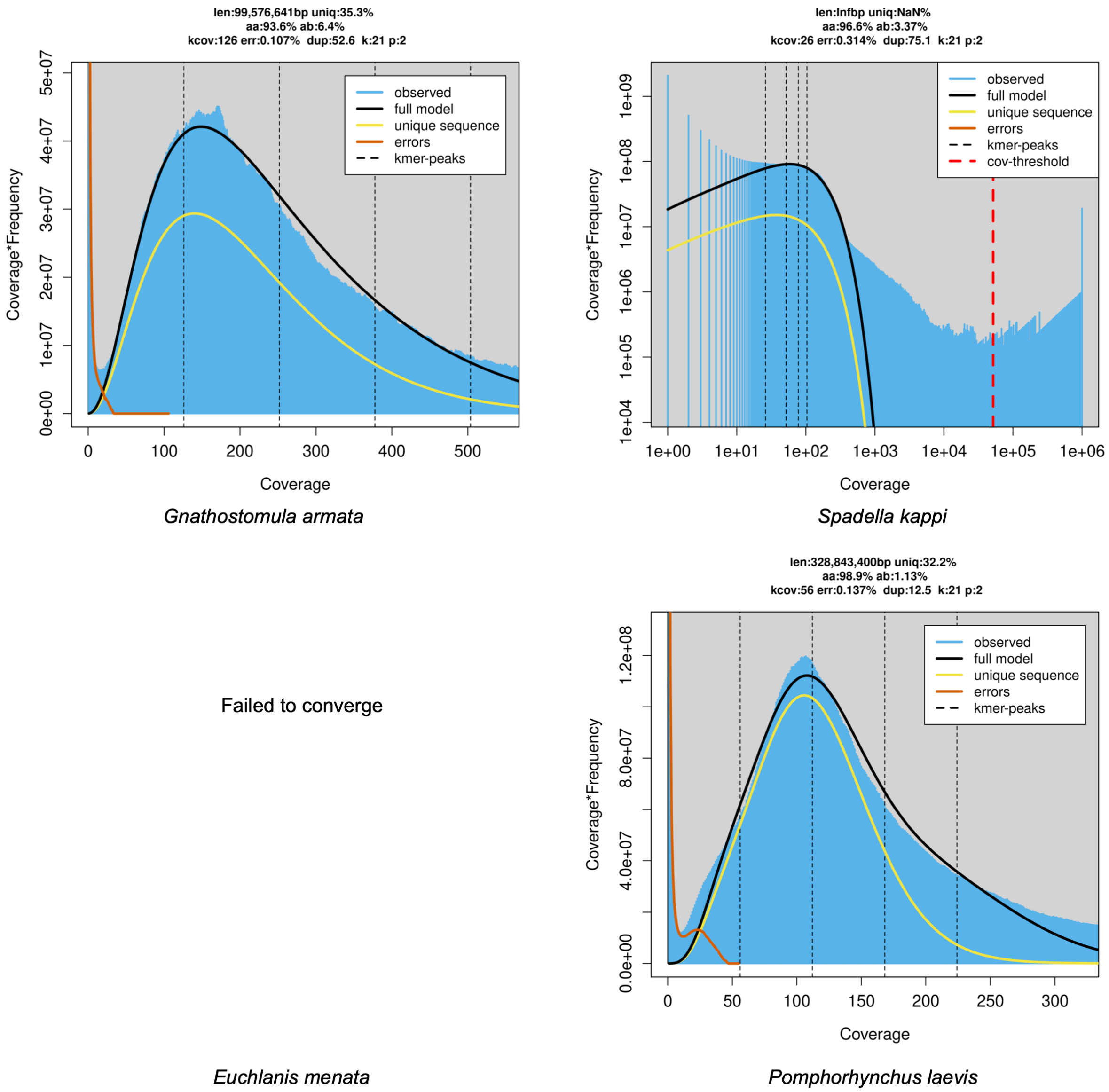


Supplemental Figure 2: Log-transformed kmer of the PacBio Hifi reads for the four species. The model failed to coverge for E. meneta. On top of each distribution, optimal model parameters such as kmer-estimated genome size (len), proportion of unique kmer (uniq), homozygosity (aa), heterozygosity (ab) and error (err) as well as kmer coverage (kcov) and read duplication rate (dup) are given as well as the provided paramters kmer size (k) and ploidy level (p).

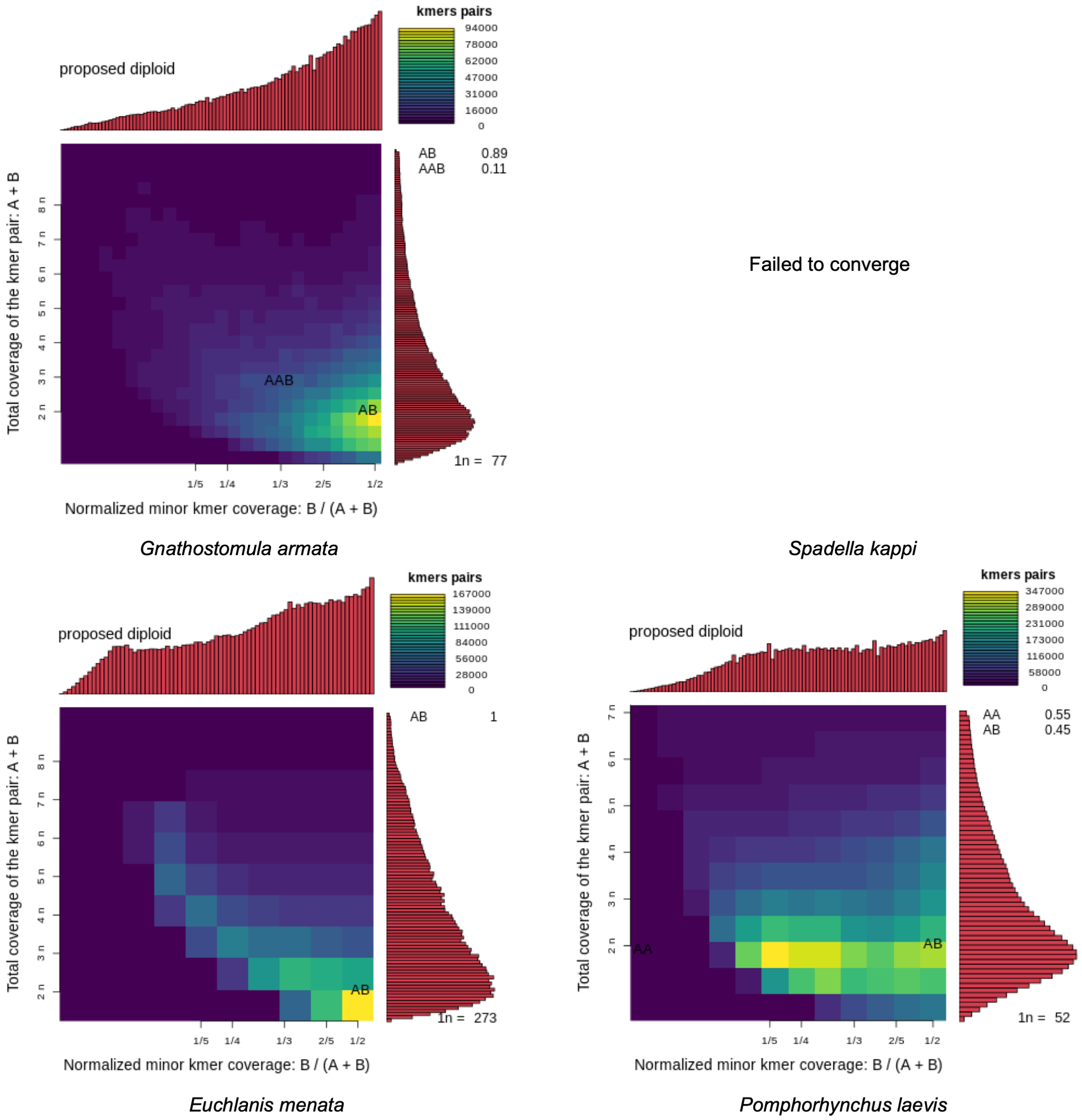


Supplemental Figure 3: SmudgePlot of kmer distribution of the PacBio Hifi reads for the four species. The model failed to coverge for *S. kappae*.
In top right corner, the legend of the heatmap of the number of reads and the proportion ploidy levels given the model.

Supplemental Table 1: The Genome Assembly and Annotation Statistics for each of the four novel genomes in this study as well the best representative genome publicly available for Chaetognatha and Monogononata (with respect to the N50 value) and the other *P. laevis* genome available. Sources: a = https://goat.genomehubs.org/[27,29]; b = this study; c = Guiglielmoni et al. 2025[30]; d = Kim et al. 2022[31]; e = Mauer et al., 2020[32].

| | **Gnathostomulida** | **Chaetognatha** | | | **Monogononta** | | | **Acanthocephala** | | |
|---|---|---|---|---|---|---|---|---|---|---|
| | *Gnathostomula armata* | *Spadella kappae* | *Flaccisagitta enflata* | | *Euchlanis meneta* | *Brachionus manjavacas* | | *Pomphorhynchus laevis* | *P. laevis* | |
| ***Assembly*** | this study | | GCA_965153355.1 | Source | this study | GCA_019054805.1 | Source | this study | GCA_012934845.2 | Source |
| Genome size (Mbp) | 128.6 | 192.3 | 794.2 | a | 91.4 | 114.2 | a | 278.3 | 253.3 | a |
| # contigs | 37 | 1,397 | 1,485 | a | 1,061 | 63 | a | 724 | 3,864 | a |
| Largest contig (kbp) | 9,578.6 | 1,025.1 | | | 527.0 | 12,101.6 | d | 2,469.4 | | |
| N50 (kbp) | 4,618.4 | 182.1 | 1,023.7 | a | 114.5 | 6,365.4 | a | 533.3 | 126.6 | a |
| L50 | 9 | 311 | 238 | a | 251 | 7 | a | 172 | 615 | a |
| GC content (%) | 39.7 | 44.3 | 41.0 | a | 26.5 | 28.0 | a | 31.9 | 33.0 | a |
| # N's per 100 kbp | 0.7 | 0.7 | | | 0.6 | | | 1.6 | | |
| Completeness Merqury (%) | 83.6 | 91.8 | 100.0 | a | 25.5 | | | 75.2 | | |
| QV Merqury | 60.6 | 49.1 | | | 47.7 | | | 55.4 | | |
| Error rate per 100 kbp | 0.1 | 1.2 | | | 1.7 | | | 0.3 | | |
| *BUSCO Metazoa_obd10* | | | | | | | | | | |
| Complete (%) | 72.9 | 87.4 | 91.3 | c | 63.9 | 85.7 | a | 44.9 | 43.1 | a |
| Duplicated (%) | 22.3 | 14.5 | 11.1 | c | 8.3 | 1.7 | a | 8.7 | 2.7 | a |
| Fragmented (%) | 5.1 | 3.6 | | | 6.3 | 4.7 | a | 6.8 | 10.1 | a |
| Missing (%) | 22.0 | 9.0 | | | 29.8 | 9.5 | a | 48.3 | 46.9 | a |

| | | | | | | | | | | |
|---|---|---|---|---|---|---|---|---|---|---|
| *BUSCO Lophotrochozoa_obd12* | | | | | | | | | | |
| Complete (%) | 71.6 | 85.5 | 91.5 | b | 64.3 | 85.9 | b | 59.6 | 62 | b |
| Duplicated (%) | 21.9 | 20.1 | 16.1 | b | 8.6 | 3.5 | b | 10.9 | 2.4 | b |
| Fragmented (%) | 3.9 | 3.0 | 2.0 | b | 3.3 | 3.0 | b | 5.1 | 5.9 | b |
| Missing (%) | 24.4 | 11.4 | 6.5 | b | 32.4 | 11.1 | b | 35.3 | 32.1 | b |
| ***Repetitiveness*** | | | | | | | | | | |
| Retroelements (%) | 8.9 | 14.9 | | | 2.9 | 25.7 | d | 40.4 | 41.9 | e |
| DNA transposons (%) | 7.1 | 1.5 | | | 1.1 | 11.5 | d | 1.1 | 2.9 | e |
| Rolling-circles (%) | 0.2 | 0.1 | | | 0.1 | 1.4 | d | 0.0 | | |
| Unclassified (%) | 25.2 | 16.1 | | | 7.4 | 0.2 | d | 11.1 | 16.2 | e |
| Small RNA (%) | 0.1 | 1.1 | | | 0.1 | | | 0.0 | | |
| Satellites (%) | 0.0 | 0.1 | | | 0.1 | 0.2 | d | 0.0 | | |
| Simple repeats (%) | 2.0 | 6.0 | | | 3.4 | 2.0 | d | 3.3 | | |
| Low complexity (%) | 0.4 | 1.0 | | | 1.1 | | | 0.6 | | |
| Total (%) | 43.8 | 40.7 | 64.2 | c | 16.2 | 41.0 | d | 56.6 | 63.0 | e |
| ***Annotation*** | | | | | | | | | | |
| # genes | 16,699 | 14,852 | 35,833 | c | 12,846 | 18,527 | d | 9,447 | 12,073 | a |
| mean gene length (bp) | 2,206 | 6,542 | | | 2,677 | | | 9,539 | | |
| mean cds length (bp) | 1,434 | 1,655 | | | 1,593 | 1,529 | d | 1,167 | | |
| mean exons per cds | 2.9 | 6.7 | | | 5.1 | | | 4.8 | | |
| *BUSCO Metazoa_obd10* | | | | | | | | | | |
| Complete (%) | 65.4 | 83.6 | 87.8 | c | 65.4 | | | 54.6 | | |

| | | | | | | |
|---|---|---|---|---|---|---|
| Duplicated (%) | 16.4 | 13.7 | 10.8 | c | 7.2 | 9.7 |
| Fragmented (%) | 6.4 | 1.4 | | | 3.6 | 3.8 |
| Missing (%) | 28.2 | 15.0 | | | 31.0 | 41.6 |

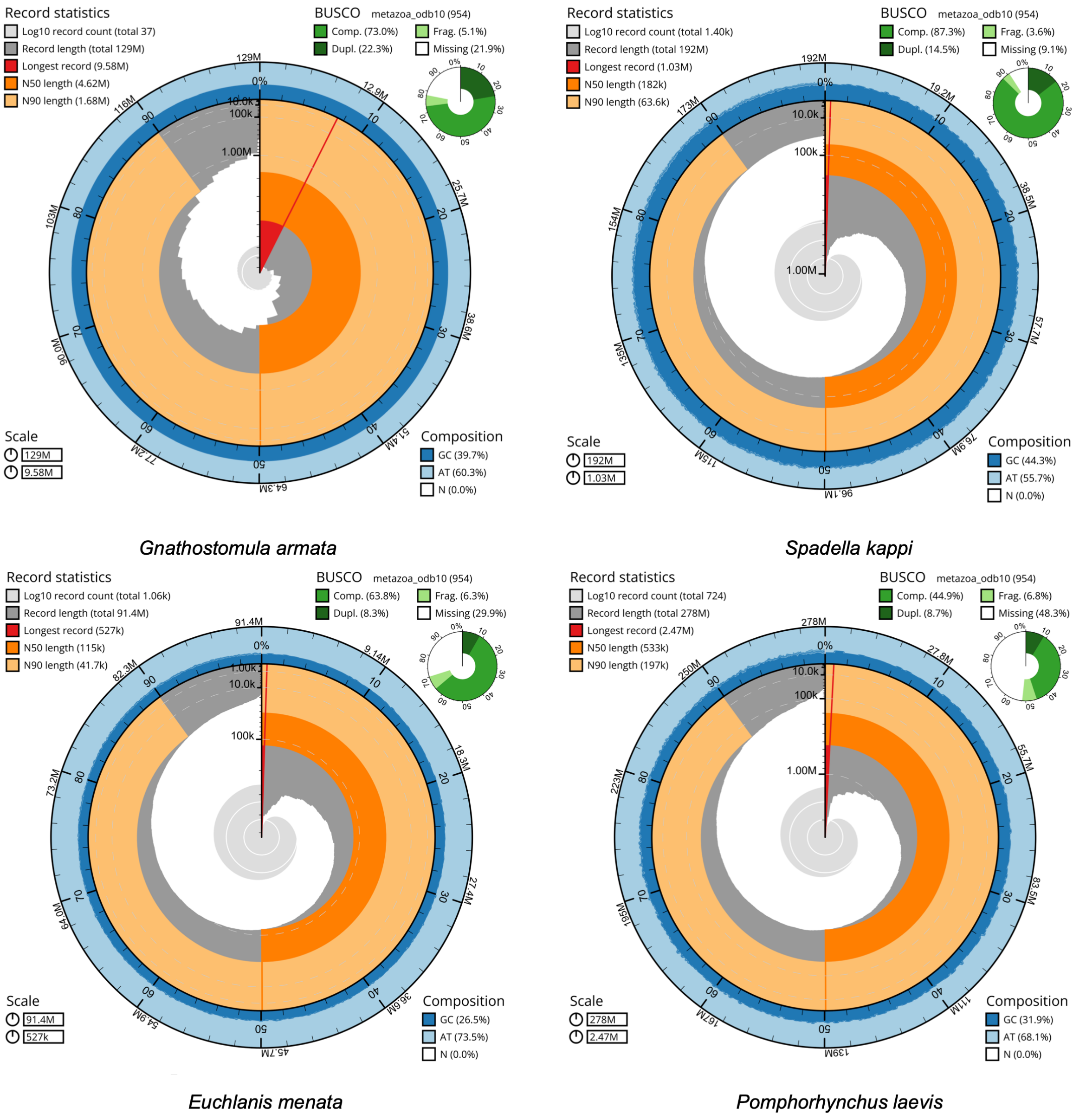


Supplemental Figure 4: Four snail plots of the novel genomes assembled as part of this study.

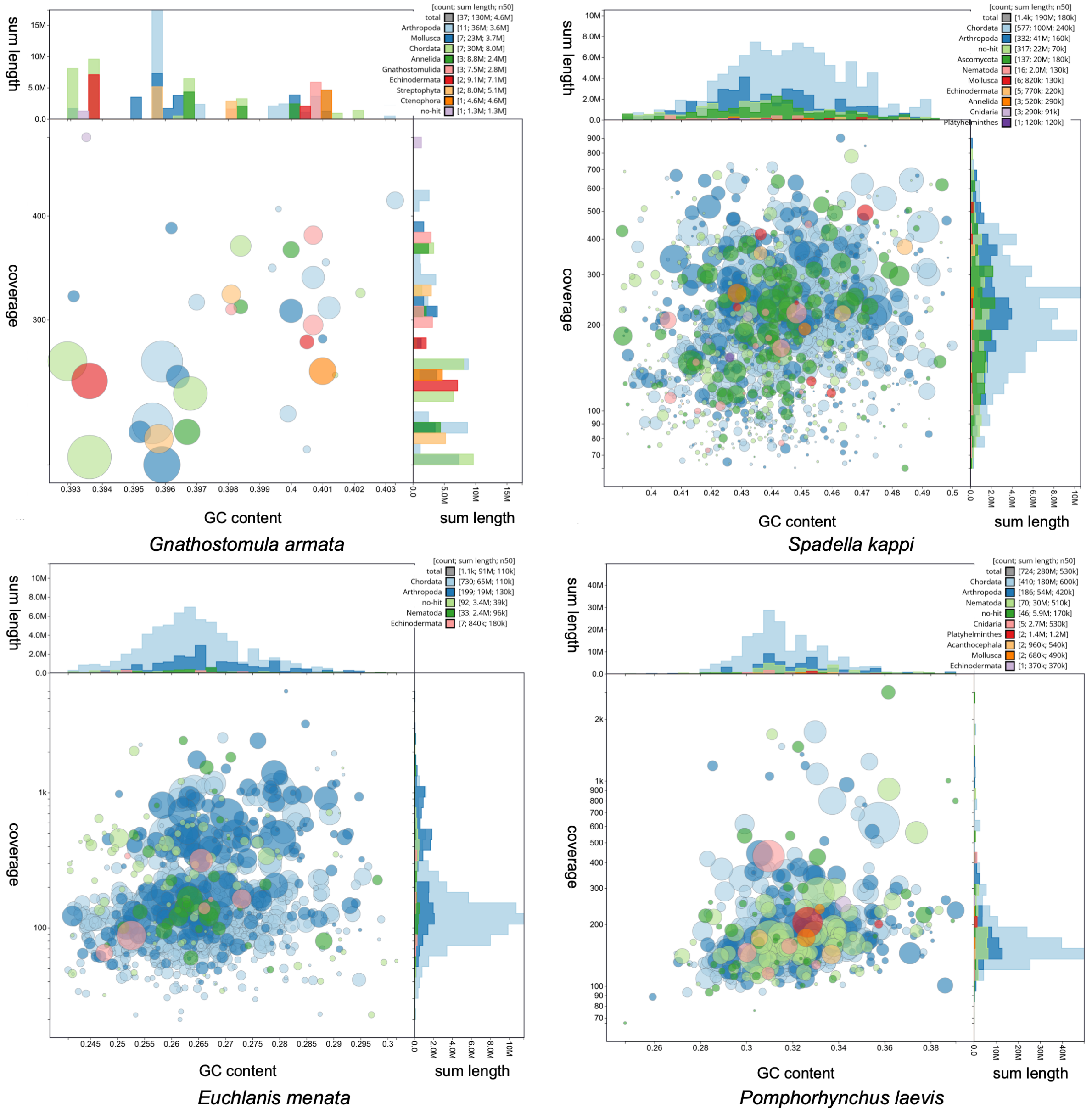


Supplemental Figure 5: BlobPlot of the four final assembled genomes. The hits to different phyla are shown in the top right corner with the number of contigs, length and N50 value. The size of the blob of each contig scales logarithmically to the length and its position to the GC content and coverage.

The complete BUSCO scores range from 44.9% to 87.4% and 59.6% to 85.5% for Metazoa_odb10 and Lophotrochozoa_odb12, respectively (Supplemental Table 1, Supplemental Figure 4). All have also relatively high duplicated BUSCO scores ranging from 8.3% to 22.3% and from 8.6% to 21.9%. The level of fragmented genes is comparatively low, ranging from 3.6% to 6.8% and from 3.0% to 5.9%. Hence, while there is a low degree of missing genes for *S. kappae* for both datasets, the percentage increases from a bit over 20% in *G. armata* to around 30% in *E. meneta* and finally, to almost 50% and 35% in *P. laevis*.

Interestingly, repetitiveness is the highest in *P. laevis* with 56.6% (Supplemental Table 1). Especially prominent here are retroelements such as LINEs, SINEs and LTRs, alongside additional unclassified repeats. The next most repetitive genome is *G. armata* with 43.8%, mostly consisting of unclassified repeats, retroelements and DNA transposons in equal proportions. The chaetognath *S. kappae* has a slightly lower proportion with 40.7%. Again, retroelements and unclassified repeats are most prominent, but simple repeats also have a larger proportion. Finally, *E. meneta* has only 16.2% repetitiveness and like for *S. kappae*, these are unclassified repeats, retroelements and simple repeats.

The number of genes ranges from 9,447 in *P. laevis* to 16,699 in *G. armata* (Supplemental Table 1). While average gene length varies strongly, the average length of the coding sequence (CDS) is quite similar across all four species. On the other hand, on average *G. armata* has clearly fewer exons per gene than the other three. Finally, the BUSCO scores for Metazoa_odb10 are usually smaller than for the assembly, but generally similar. However, for the two syndermatans (*E. meneta* and *P. laevis*), these BUSCO scores actually increase, especially for *P. laevis* with 54.6%.

**Phylogenetic Assessment: Canary Sequence Methodology**

Supplemental Table 2: the methodological incongruence that resulted in the selection of each of the sequences of interest. As all sequences were found to be stable, the "result" is not included in this table.

| Taxa name | Methodological incongruence | Result |
|---|---|---|
| *Brachionus angularis* | nRCFV (0.0163) | Stable, but excluded due to compositional heterogeneity |
| *Gnathostomula armata* | LB-Score (22.545) | Stable |
| *Pomphorhynchus laevis* | LB-Score (38.022) | Stable |
| *Pomphorhynchus laevis* (this study) | LB-Score (35.657) | Stable |
| *Paraspadella gotoi* | nRCFV (0.0159) | Stable, but excluded due to compositional heterogeneity |
| *Seison nabaliae* | LB-Score (33.714) | Stable |
| *Spadella kappae* | nRCFV (0.0182) | Stable, but excluded due to compositional heterogeneity |

**Phylogenetic Assessment: CAT-PMSF**

Supplemental Figures 6-14: The finale CAT-PMSF topologies recovered for the CAT-PMSF models generated under each constraint.

Supplemental Figure 6: Chaetognatha-first

Supplemental Figure 7: Gnathostomulida-first (note: this is also Pararotatoria, as both were simultaneously recovered in the initial phylogeny)

Supplemental Figure 8: Eurotatoria

Supplemental Figure 9: Hemirotifera

Supplemental Figure 10: Lemniscea

Supplemental Figure 11: Monogononta-sister

Supplemental Figure 12: Acanthocephala-Bdelloidea Sister and Seisonidea as sister to all Rotifera

Supplemental Figure 13: Acanthocephala-Monogononta Sister and Seisonidea as sister to all Rotifera

Supplemental Figure 14: Bdelloidea-Monogononta Sister and Seisonidea as sister to all Rotifera

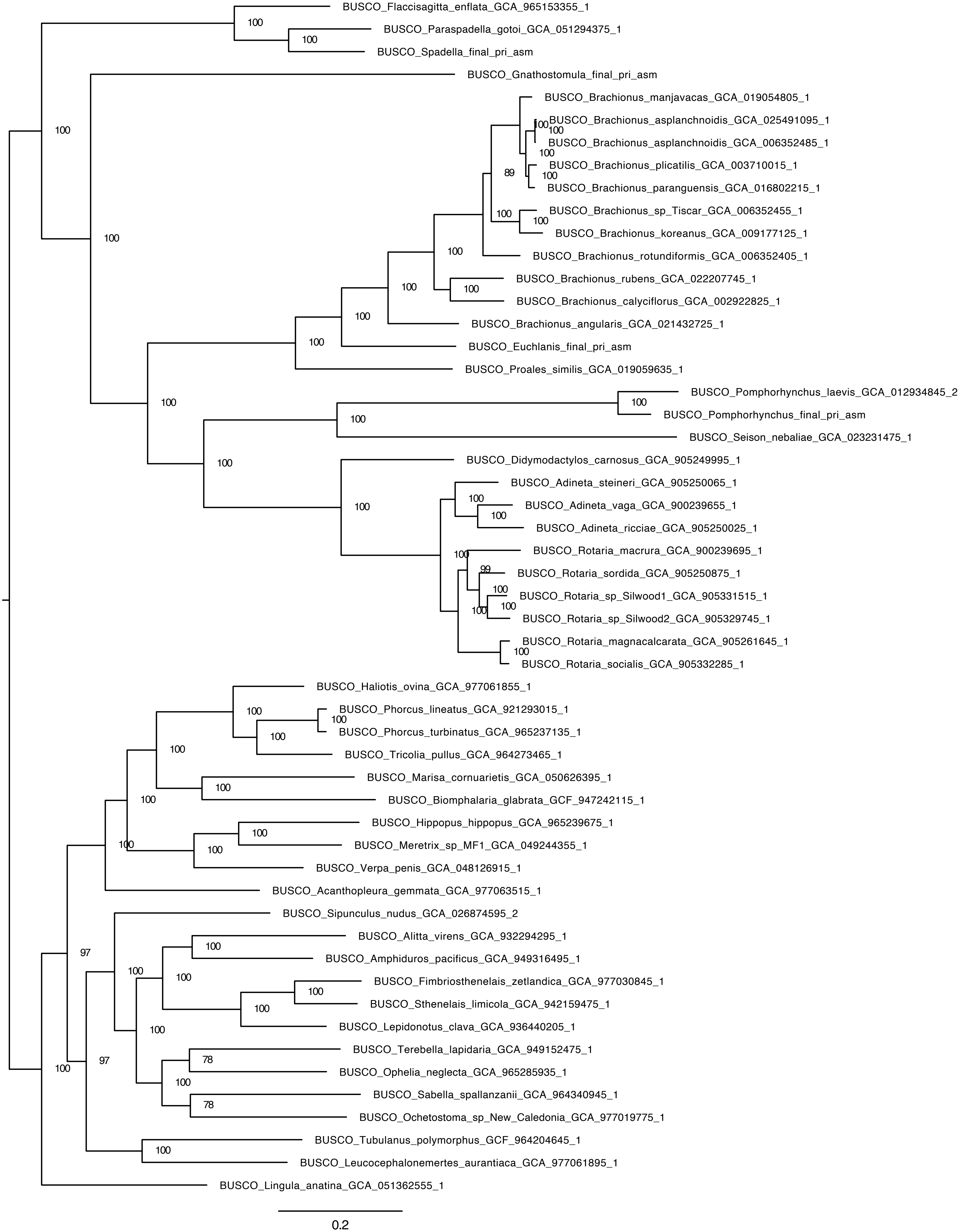
BUSCO_Flaccisagitta_enflata_GCA_965153355_1
BUSCO_Paraspadella_gotoi_GCA_051294375_1
BUSCO_Spadella_final_pri_asm
BUSCO_Gnathostomula_final_pri_asm
BUSCO_Brachionus_manjavacas_GCA_019054805_1
BUSCO_Brachionus_asplanchnoidis_GCA_025491095_1
BUSCO_Brachionus_asplanchnoidis_GCA_006352485_1
BUSCO_Brachionus_plicatilis_GCA_003710015_1
BUSCO_Brachionus_paranguensis_GCA_016802215_1
BUSCO_Brachionus_sp_Tiscar_GCA_006352455_1
BUSCO_Brachionus_koreanus_GCA_009177125_1
BUSCO_Brachionus_rotundiformis_GCA_006352405_1
BUSCO_Brachionus_rubens_GCA_022207745_1
BUSCO_Brachionus_calyciflorus_GCA_002922825_1
BUSCO_Brachionus_angularis_GCA_021432725_1
BUSCO_Euchlanis_final_pri_asm
BUSCO_Proales_similis_GCA_019059635_1
BUSCO_Pomphorhynchus_laevis_GCA_012934845_2
BUSCO_Pomphorhynchus_final_pri_asm
BUSCO_Seison_nebaliae_GCA_023231475_1
BUSCO_Didymodactylos_carnosus_GCA_905249995_1
BUSCO_Adineta_steineri_GCA_905250065_1
BUSCO_Adineta_vaga_GCA_900239655_1
BUSCO_Adineta_ricciae_GCA_905250025_1
BUSCO_Rotaria_macrura_GCA_900239695_1
BUSCO_Rotaria_sordida_GCA_905250875_1
BUSCO_Rotaria_sp_Silwood1_GCA_905331515_1
BUSCO_Rotaria_sp_Silwood2_GCA_905329745_1
BUSCO_Rotaria_magnacalcarata_GCA_905261645_1
BUSCO_Rotaria_socialis_GCA_905332285_1
BUSCO_Haliotis_ovina_GCA_977061855_1
BUSCO_Phorcus_lineatus_GCA_921293015_1
BUSCO_Phorcus_turbinatus_GCA_965237135_1
BUSCO_Tricolia_pullus_GCA_964273465_1
BUSCO_Marisa_cornuarietis_GCA_050626395_1
BUSCO_Biomphalaria_glabrata_GCF_947242115_1
BUSCO_Hippopus_hippopus_GCA_965239675_1
BUSCO_Meretrix_sp_MF1_GCA_049244355_1
BUSCO_Verpa_penis_GCA_048126915_1
BUSCO_Acanthopleura_gemmata_GCA_977063515_1
BUSCO_Sipunculus_nudus_GCA_026874595_2
BUSCO_Alitta_virens_GCA_932294295_1
BUSCO_Amphiduros_pacificus_GCA_949316495_1
BUSCO_Fimbriosthenelais_zetlandica_GCA_977030845_1
BUSCO_Sthenelais_limicola_GCA_942159475_1
BUSCO_Lepidonotus_clava_GCA_936440205_1
BUSCO_Terebella_lapidaria_GCA_949152475_1
BUSCO_Ophelia_neglecta_GCA_965285935_1
BUSCO_Sabella_spallanzanii_GCA_964340945_1
BUSCO_Ochetostoma_sp_New_Caledonia_GCA_977019775_1
BUSCO_Tubulanus_polymorphus_GCF_964204645_1
BUSCO_Leucocephalonemertes_aurantiaca_GCA_977061895_1
BUSCO_Lingula_anatina_GCA_051362555_1
0.2

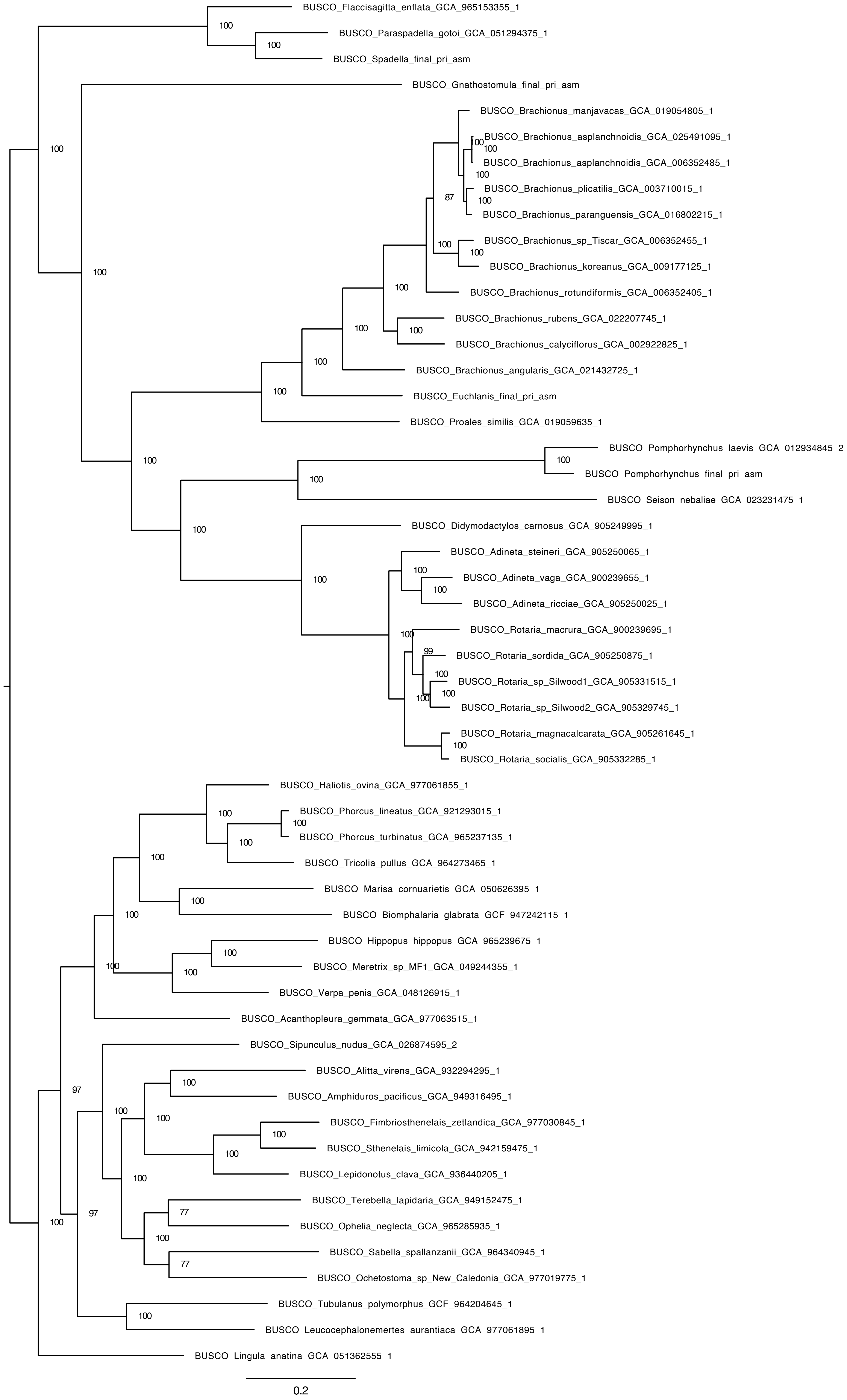
BUSCO_Flaccisagitta_enflata_GCA_965153355_1
BUSCO_Paraspadella_gotoi_GCA_051294375_1
BUSCO_Spadella_final_pri_asm
BUSCO_Gnathostomula_final_pri_asm
BUSCO_Brachionus_manjavacas_GCA_019054805_1
BUSCO_Brachionus_asplanchnoidis_GCA_025491095_1
BUSCO_Brachionus_asplanchnoidis_GCA_006352485_1
BUSCO_Brachionus_plicatilis_GCA_003710015_1
BUSCO_Brachionus_paranguensis_GCA_016802215_1
BUSCO_Brachionus_sp_Tiscar_GCA_006352455_1
BUSCO_Brachionus_koreanus_GCA_009177125_1
BUSCO_Brachionus_rotundiformis_GCA_006352405_1
BUSCO_Brachionus_rubens_GCA_022207745_1
BUSCO_Brachionus_calyciflorus_GCA_002922825_1
BUSCO_Brachionus_angularis_GCA_021432725_1
BUSCO_Euchlanis_final_pri_asm
BUSCO_Proales_similis_GCA_019059635_1
BUSCO_Pomphorhynchus_laevis_GCA_012934845_2
BUSCO_Pomphorhynchus_final_pri_asm
BUSCO_Seison_nebaliae_GCA_023231475_1
BUSCO_Didymodactylos_carnosus_GCA_905249995_1
BUSCO_Adineta_steineri_GCA_905250065_1
BUSCO_Adineta_vaga_GCA_900239655_1
BUSCO_Adineta_ricciae_GCA_905250025_1
BUSCO_Rotaria_macrura_GCA_900239695_1
BUSCO_Rotaria_sordida_GCA_905250875_1
BUSCO_Rotaria_sp_Silwood1_GCA_905331515_1
BUSCO_Rotaria_sp_Silwood2_GCA_905329745_1
BUSCO_Rotaria_magnacalcarata_GCA_905261645_1
BUSCO_Rotaria_socialis_GCA_905332285_1
BUSCO_Haliotis_ovina_GCA_977061855_1
BUSCO_Phorcus_lineatus_GCA_921293015_1
BUSCO_Phorcus_turbinatus_GCA_965237135_1
BUSCO_Tricolia_pullus_GCA_964273465_1
BUSCO_Marisa_cornuarietis_GCA_050626395_1
BUSCO_Biomphalaria_glabrata_GCF_947242115_1
BUSCO_Hippopus_hippopus_GCA_965239675_1
BUSCO_Meretrix_sp_MF1_GCA_049244355_1
BUSCO_Verpa_penis_GCA_048126915_1
BUSCO_Acanthopleura_gemmata_GCA_977063515_1
BUSCO_Sipunculus_nudus_GCA_026874595_2
BUSCO_Alitta_virens_GCA_932294295_1
BUSCO_Amphiduros_pacificus_GCA_949316495_1
BUSCO_Fimbriosthenelais_zetlandica_GCA_977030845_1
BUSCO_Sthenelais_limicola_GCA_942159475_1
BUSCO_Lepidonotus_clava_GCA_936440205_1
BUSCO_Terebella_lapidaria_GCA_949152475_1
BUSCO_Ophelia_neglecta_GCA_965285935_1
BUSCO_Sabella_spallanzanii_GCA_964340945_1
BUSCO_Ochetostoma_sp_New_Caledonia_GCA_977019775_1
BUSCO_Tubulanus_polymorphus_GCF_964204645_1
BUSCO_Leucocephalonemertes_aurantiaca_GCA_977061895_1
BUSCO_Lingula_anatina_GCA_051362555_1
0.2

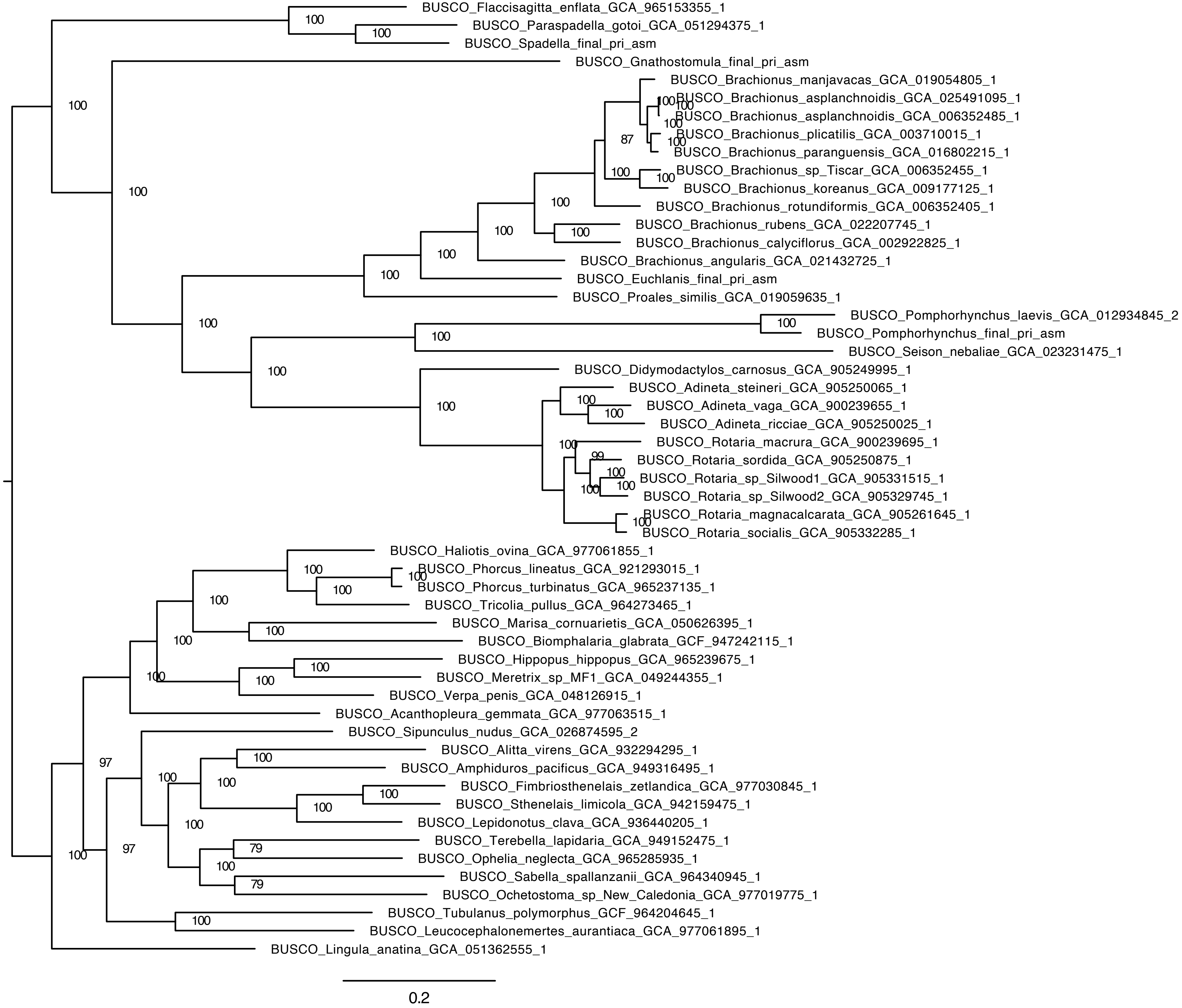
BUSCO_Flaccisagitta_enflata_GCA_965153355_1
BUSCO_Paraspadella_gotoi_GCA_051294375_1
BUSCO_Spadella_final_pri_asm
BUSCO_Gnathostomula_final_pri_asm
BUSCO_Brachionus_manjavacas_GCA_019054805_1
BUSCO_Brachionus_asplanchnoidis_GCA_025491095_1
BUSCO_Brachionus_asplanchnoidis_GCA_006352485_1
BUSCO_Brachionus_plicatilis_GCA_003710015_1
BUSCO_Brachionus_paranguensis_GCA_016802215_1
BUSCO_Brachionus_sp_Tiscar_GCA_006352455_1
BUSCO_Brachionus_koreanus_GCA_009177125_1
BUSCO_Brachionus_rotundiformis_GCA_006352405_1
BUSCO_Brachionus_rubens_GCA_022207745_1
BUSCO_Brachionus_calyciflorus_GCA_002922825_1
BUSCO_Brachionus_angularis_GCA_021432725_1
BUSCO_Euchlanis_final_pri_asm
BUSCO_Proales_similis_GCA_019059635_1
BUSCO_Pomphorhynchus_laevis_GCA_012934845_2
BUSCO_Pomphorhynchus_final_pri_asm
BUSCO_Seison_nebaliae_GCA_023231475_1
BUSCO_Didymodactylos_carnosus_GCA_905249995_1
BUSCO_Adineta_steineri_GCA_905250065_1
BUSCO_Adineta_vaga_GCA_900239655_1
BUSCO_Adineta_ricciae_GCA_905250025_1
BUSCO_Rotaria_macrura_GCA_900239695_1
BUSCO_Rotaria_sordida_GCA_905250875_1
BUSCO_Rotaria_sp_Silwood1_GCA_905331515_1
BUSCO_Rotaria_sp_Silwood2_GCA_905329745_1
BUSCO_Rotaria_magnacalcarata_GCA_905261645_1
BUSCO_Rotaria_socialis_GCA_905332285_1
BUSCO_Haliotis_ovina_GCA_977061855_1
BUSCO_Phorcus_lineatus_GCA_921293015_1
BUSCO_Phorcus_turbinatus_GCA_965237135_1
BUSCO_Tricolia_pullus_GCA_964273465_1
BUSCO_Marisa_cornuarietis_GCA_050626395_1
BUSCO_Biomphalaria_glabrata_GCF_947242115_1
BUSCO_Hippopus_hippopus_GCA_965239675_1
BUSCO_Meretrix_sp_MF1_GCA_049244355_1
BUSCO_Verpa_penis_GCA_048126915_1
BUSCO_Acanthopleura_gemmata_GCA_977063515_1
BUSCO_Sipunculus_nudus_GCA_026874595_2
BUSCO_Alitta_virens_GCA_932294295_1
BUSCO_Amphiduros_pacificus_GCA_949316495_1
BUSCO_Fimbriosthenelais_zetlandica_GCA_977030845_1
BUSCO_Sthenelais_limicola_GCA_942159475_1
BUSCO_Lepidonotus_clava_GCA_936440205_1
BUSCO_Terebella_lapidaria_GCA_949152475_1
BUSCO_Ophelia_neglecta_GCA_965285935_1
BUSCO_Sabella_spallanzanii_GCA_964340945_1
BUSCO_Ochetostoma_sp_New_Caledonia_GCA_977019775_1
BUSCO_Tubulanus_polymorphus_GCF_964204645_1
BUSCO_Leucocephalonemertes_aurantiaca_GCA_977061895_1
BUSCO_Lingula_anatina_GCA_051362555_1
0.2

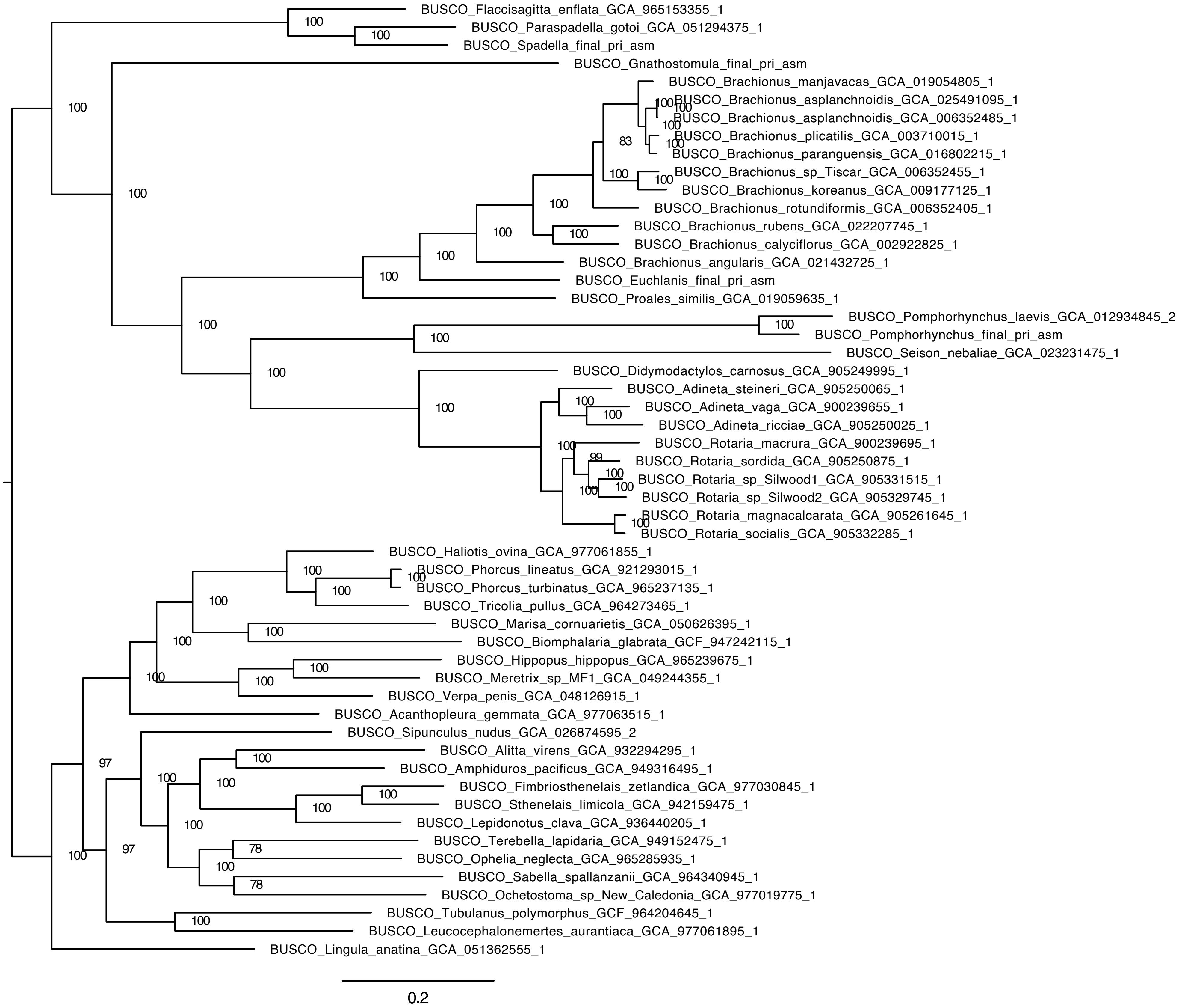
BUSCO_Flaccisagitta_enflata_GCA_965153355_1
BUSCO_Paraspadella_gotoi_GCA_051294375_1
BUSCO_Spadella_final_pri_asm
BUSCO_Gnathostomula_final_pri_asm
BUSCO_Brachionus_manjavacas_GCA_019054805_1
BUSCO_Brachionus_asplanchnoidis_GCA_025491095_1
BUSCO_Brachionus_asplanchnoidis_GCA_006352485_1
BUSCO_Brachionus_plicatilis_GCA_003710015_1
BUSCO_Brachionus_paranguensis_GCA_016802215_1
BUSCO_Brachionus_sp_Tiscar_GCA_006352455_1
BUSCO_Brachionus_koreanus_GCA_009177125_1
BUSCO_Brachionus_rotundiformis_GCA_006352405_1
BUSCO_Brachionus_rubens_GCA_022207745_1
BUSCO_Brachionus_calyciflorus_GCA_002922825_1
BUSCO_Brachionus_angularis_GCA_021432725_1
BUSCO_Euchlanis_final_pri_asm
BUSCO_Proales_similis_GCA_019059635_1
BUSCO_Pomphorhynchus_laevis_GCA_012934845_2
BUSCO_Pomphorhynchus_final_pri_asm
BUSCO_Seison_nebaliae_GCA_023231475_1
BUSCO_Didymodactylos_carnosus_GCA_905249995_1
BUSCO_Adineta_steineri_GCA_905250065_1
BUSCO_Adineta_vaga_GCA_900239655_1
BUSCO_Adineta_ricciae_GCA_905250025_1
BUSCO_Rotaria_macrura_GCA_900239695_1
BUSCO_Rotaria_sordida_GCA_905250875_1
BUSCO_Rotaria_sp_Silwood1_GCA_905331515_1
BUSCO_Rotaria_sp_Silwood2_GCA_905329745_1
BUSCO_Rotaria_magnacalcarata_GCA_905261645_1
BUSCO_Rotaria_socialis_GCA_905332285_1
BUSCO_Haliotis_ovina_GCA_977061855_1
BUSCO_Phorcus_lineatus_GCA_921293015_1
BUSCO_Phorcus_turbinatus_GCA_965237135_1
BUSCO_Tricolia_pullus_GCA_964273465_1
BUSCO_Marisa_cornuarietis_GCA_050626395_1
BUSCO_Biomphalaria_glabrata_GCF_947242115_1
BUSCO_Hippopus_hippopus_GCA_965239675_1
BUSCO_Meretrix_sp_MF1_GCA_049244355_1
BUSCO_Verpa_penis_GCA_048126915_1
BUSCO_Acanthopleura_gemmata_GCA_977063515_1
BUSCO_Sipunculus_nudus_GCA_026874595_2
BUSCO_Alitta_virens_GCA_932294295_1
BUSCO_Amphiduros_pacificus_GCA_949316495_1
BUSCO_Fimbriosthenelais_zetlandica_GCA_977030845_1
BUSCO_Sthenelais_limicola_GCA_942159475_1
BUSCO_Lepidonotus_clava_GCA_936440205_1
BUSCO_Terebella_lapidaria_GCA_949152475_1
BUSCO_Ophelia_neglecta_GCA_965285935_1
BUSCO_Sabella_spallanzanii_GCA_964340945_1
BUSCO_Ochetostoma_sp_New_Caledonia_GCA_977019775_1
BUSCO_Tubulanus_polymorphus_GCF_964204645_1
BUSCO_Leucocephalonemertes_aurantiaca_GCA_977061895_1
BUSCO_Lingula_anatina_GCA_051362555_1
0.2

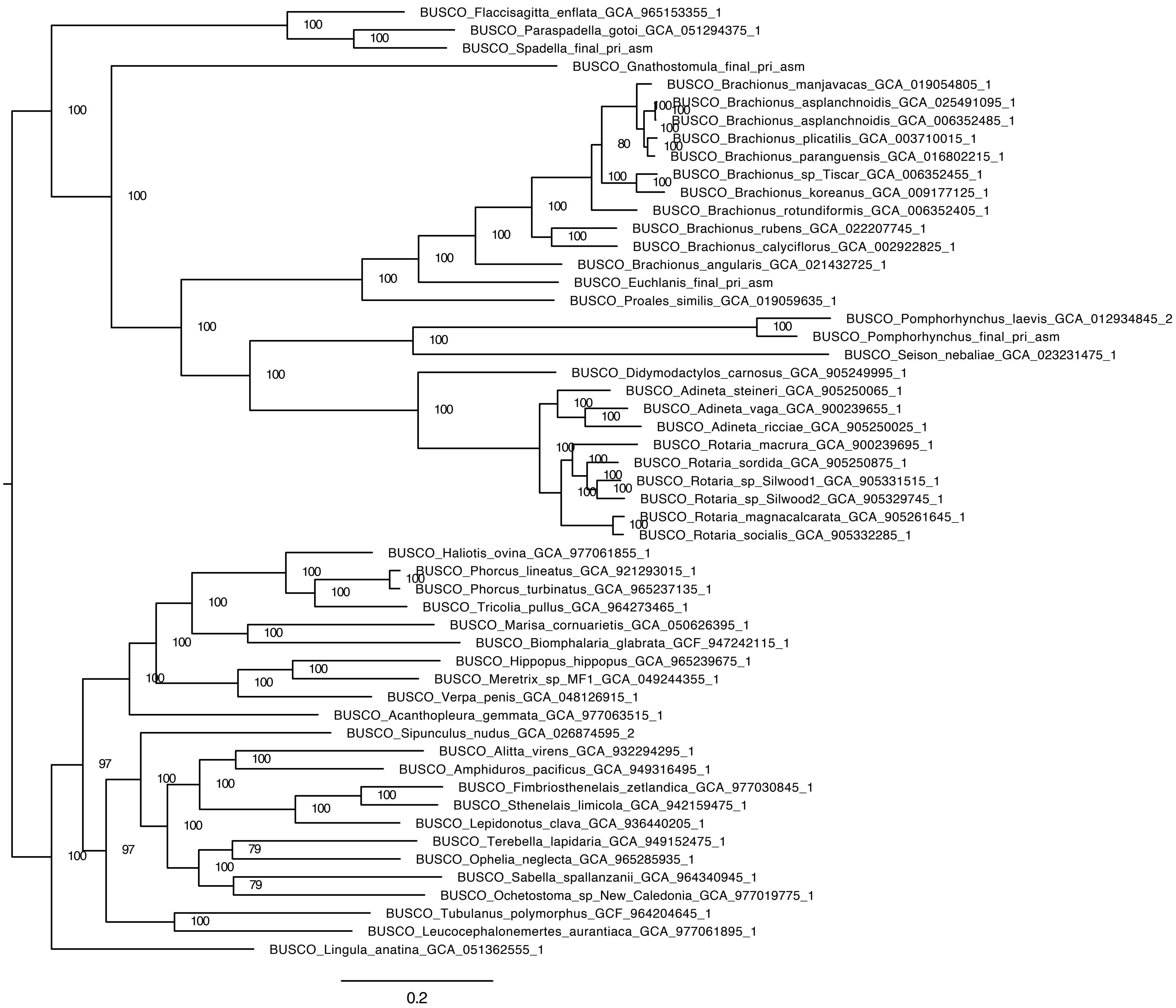
BUSCO_Flaccisagitta_enflata_GCA_965153355_1
BUSCO_Paraspadella_gotoi_GCA_051294375_1
BUSCO_Spadella_final_pri_asm
BUSCO_Gnathostomula_final_pri_asm
BUSCO_Brachionus_manjavacas_GCA_019054805_1
BUSCO_Brachionus_asplanchnoidis_GCA_025491095_1
BUSCO_Brachionus_asplanchnoidis_GCA_006352485_1
BUSCO_Brachionus_plicatilis_GCA_003710015_1
BUSCO_Brachionus_paranguensis_GCA_016802215_1
BUSCO_Brachionus_sp_Tiscar_GCA_006352455_1
BUSCO_Brachionus_koreanus_GCA_009177125_1
BUSCO_Brachionus_rotundiformis_GCA_006352405_1
BUSCO_Brachionus_rubens_GCA_022207745_1
BUSCO_Brachionus_calyciflorus_GCA_002922825_1
BUSCO_Brachionus_angularis_GCA_021432725_1
BUSCO_Euchlanis_final_pri_asm
BUSCO_Proales_similis_GCA_019059635_1
BUSCO_Pomphorhynchus_laevis_GCA_012934845_2
BUSCO_Pomphorhynchus_final_pri_asm
BUSCO_Seison_nebaliae_GCA_023231475_1
BUSCO_Didymodactylos_carnosus_GCA_905249995_1
BUSCO_Adineta_steineri_GCA_905250065_1
BUSCO_Adineta_vaga_GCA_900239655_1
BUSCO_Adineta_ricciae_GCA_905250025_1
BUSCO_Rotaria_macrura_GCA_900239695_1
BUSCO_Rotaria_sordida_GCA_905250875_1
BUSCO_Rotaria_sp_Silwood1_GCA_905331515_1
BUSCO_Rotaria_sp_Silwood2_GCA_905329745_1
BUSCO_Rotaria_magnacalcarata_GCA_905261645_1
BUSCO_Rotaria_socialis_GCA_905332285_1
BUSCO_Haliotis_ovina_GCA_977061855_1
BUSCO_Phorcus_lineatus_GCA_921293015_1
BUSCO_Phorcus_turbinatus_GCA_965237135_1
BUSCO_Tricolia_pullus_GCA_964273465_1
BUSCO_Marisa_cornuarietis_GCA_050626395_1
BUSCO_Biomphalaria_glabrata_GCF_947242115_1
BUSCO_Hippopus_hippopus_GCA_965239675_1
BUSCO_Meretrix_sp_MF1_GCA_049244355_1
BUSCO_Verpa_penis_GCA_048126915_1
BUSCO_Acanthopleura_gemmata_GCA_977063515_1
BUSCO_Sipunculus_nudus_GCA_026874595_2
BUSCO_Alitta_virens_GCA_932294295_1
BUSCO_Amphiduros_pacificus_GCA_949316495_1
BUSCO_Fimbriosthenelais_zetlandica_GCA_977030845_1
BUSCO_Sthenelais_limicola_GCA_942159475_1
BUSCO_Lepidonotus_clava_GCA_936440205_1
BUSCO_Terebella_lapidaria_GCA_949152475_1
BUSCO_Ophelia_neglecta_GCA_965285935_1
BUSCO_Sabella_spallanzanii_GCA_964340945_1
BUSCO_Ochetostoma_sp_New_Caledonia_GCA_977019775_1
BUSCO_Tubulanus_polymorphus_GCF_964204645_1
BUSCO_Leucocephalonemertes_aurantiaca_GCA_977061895_1
BUSCO_Lingula_anatina_GCA_051362555_1
0.2

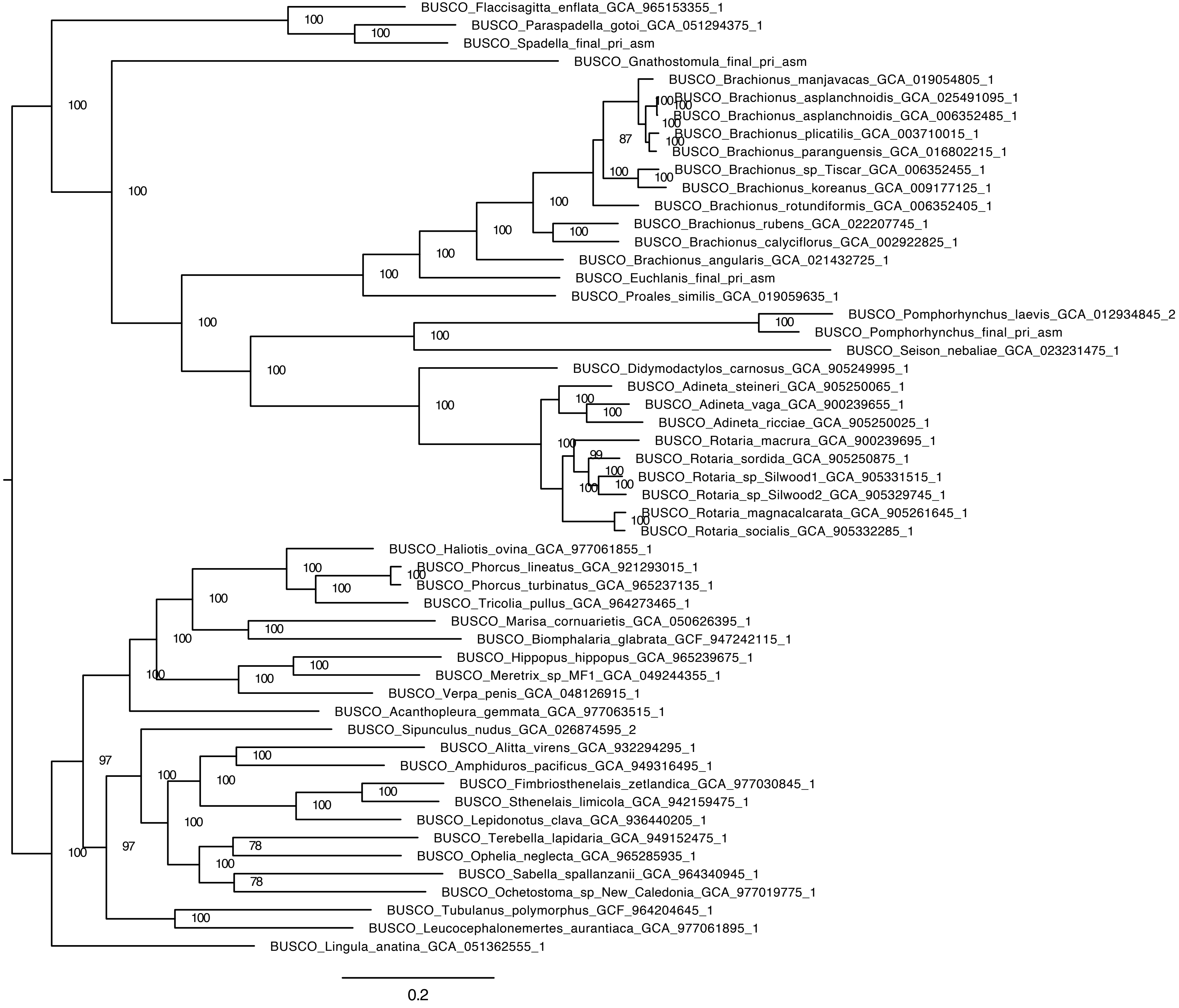
BUSCO_Flaccisagitta_enflata_GCA_965153355_1
BUSCO_Paraspadella_gotoi_GCA_051294375_1
BUSCO_Spadella_final_pri_asm
BUSCO_Gnathostomula_final_pri_asm
BUSCO_Brachionus_manjavacas_GCA_019054805_1
BUSCO_Brachionus_asplanchnoidis_GCA_025491095_1
BUSCO_Brachionus_asplanchnoidis_GCA_006352485_1
BUSCO_Brachionus_plicatilis_GCA_003710015_1
BUSCO_Brachionus_paranguensis_GCA_016802215_1
BUSCO_Brachionus_sp_Tiscar_GCA_006352455_1
BUSCO_Brachionus_koreanus_GCA_009177125_1
BUSCO_Brachionus_rotundiformis_GCA_006352405_1
BUSCO_Brachionus_rubens_GCA_022207745_1
BUSCO_Brachionus_calyciflorus_GCA_002922825_1
BUSCO_Brachionus_angularis_GCA_021432725_1
BUSCO_Euchlanis_final_pri_asm
BUSCO_Proales_similis_GCA_019059635_1
BUSCO_Pomphorhynchus_laevis_GCA_012934845_2
BUSCO_Pomphorhynchus_final_pri_asm
BUSCO_Seison_nebaliae_GCA_023231475_1
BUSCO_Didymodactylos_carnosus_GCA_905249995_1
BUSCO_Adineta_steineri_GCA_905250065_1
BUSCO_Adineta_vaga_GCA_900239655_1
BUSCO_Adineta_ricciae_GCA_905250025_1
BUSCO_Rotaria_macrura_GCA_900239695_1
BUSCO_Rotaria_sordida_GCA_905250875_1
BUSCO_Rotaria_sp_Silwood1_GCA_905331515_1
BUSCO_Rotaria_sp_Silwood2_GCA_905329745_1
BUSCO_Rotaria_magnacalcarata_GCA_905261645_1
BUSCO_Rotaria_socialis_GCA_905332285_1
BUSCO_Haliotis_ovina_GCA_977061855_1
BUSCO_Phorcus_lineatus_GCA_921293015_1
BUSCO_Phorcus_turbinatus_GCA_965237135_1
BUSCO_Tricolia_pullus_GCA_964273465_1
BUSCO_Marisa_cornuarietis_GCA_050626395_1
BUSCO_Biomphalaria_glabrata_GCF_947242115_1
BUSCO_Hippopus_hippopus_GCA_965239675_1
BUSCO_Meretrix_sp_MF1_GCA_049244355_1
BUSCO_Verpa_penis_GCA_048126915_1
BUSCO_Acanthopleura_gemmata_GCA_977063515_1
BUSCO_Sipunculus_nudus_GCA_026874595_2
BUSCO_Alitta_virens_GCA_932294295_1
BUSCO_Amphiduros_pacificus_GCA_949316495_1
BUSCO_Fimbriosthenelais_zetlandica_GCA_977030845_1
BUSCO_Sthenelais_limicola_GCA_942159475_1
BUSCO_Lepidonotus_clava_GCA_936440205_1
BUSCO_Terebella_lapidaria_GCA_949152475_1
BUSCO_Ophelia_neglecta_GCA_965285935_1
BUSCO_Sabella_spallanzanii_GCA_964340945_1
BUSCO_Ochetostoma_sp_New_Caledonia_GCA_977019775_1
BUSCO_Tubulanus_polymorphus_GCF_964204645_1
BUSCO_Leucocephalonemertes_aurantiaca_GCA_977061895_1
BUSCO_Lingula_anatina_GCA_051362555_1
0.2

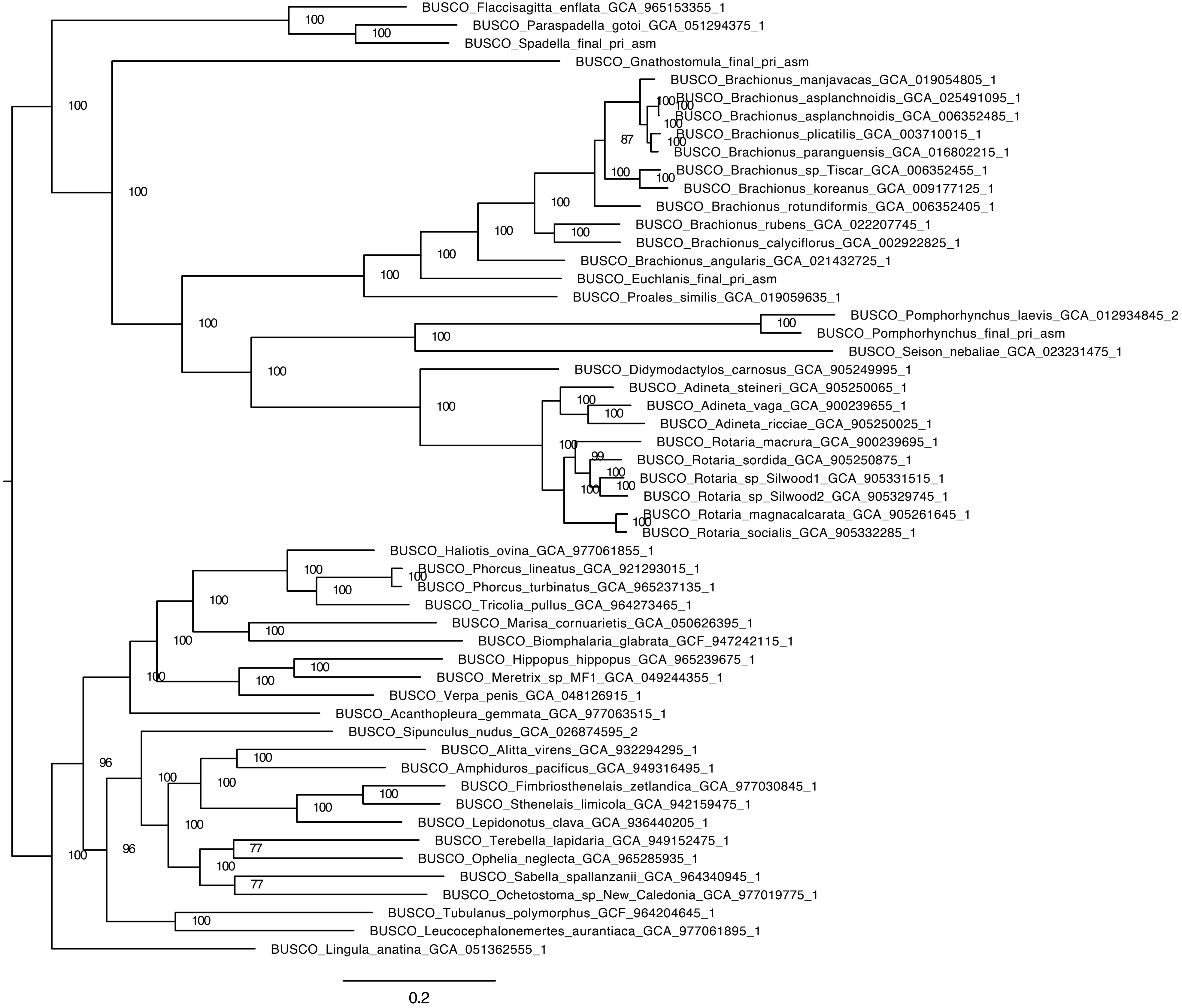
BUSCO_Flaccisagitta_enflata_GCA_965153355_1
BUSCO_Paraspadella_gotoi_GCA_051294375_1
BUSCO_Spadella_final_pri_asm
BUSCO_Gnathostomula_final_pri_asm
BUSCO_Brachionus_manjavacas_GCA_019054805_1
BUSCO_Brachionus_asplanchnoidis_GCA_025491095_1
BUSCO_Brachionus_asplanchnoidis_GCA_006352485_1
BUSCO_Brachionus_plicatilis_GCA_003710015_1
BUSCO_Brachionus_paranguensis_GCA_016802215_1
BUSCO_Brachionus_sp_Tiscar_GCA_006352455_1
BUSCO_Brachionus_koreanus_GCA_009177125_1
BUSCO_Brachionus_rotundiformis_GCA_006352405_1
BUSCO_Brachionus_rubens_GCA_022207745_1
BUSCO_Brachionus_calyciflorus_GCA_002922825_1
BUSCO_Brachionus_angularis_GCA_021432725_1
BUSCO_Euchlanis_final_pri_asm
BUSCO_Proales_similis_GCA_019059635_1
BUSCO_Pomphorhynchus_laevis_GCA_012934845_2
BUSCO_Pomphorhynchus_final_pri_asm
BUSCO_Seison_nebaliae_GCA_023231475_1
BUSCO_Didymodactylos_carnosus_GCA_905249995_1
BUSCO_Adineta_steineri_GCA_905250065_1
BUSCO_Adineta_vaga_GCA_900239655_1
BUSCO_Adineta_ricciae_GCA_905250025_1
BUSCO_Rotaria_macrura_GCA_900239695_1
BUSCO_Rotaria_sordida_GCA_905250875_1
BUSCO_Rotaria_sp_Silwood1_GCA_905331515_1
BUSCO_Rotaria_sp_Silwood2_GCA_905329745_1
BUSCO_Rotaria_magnacalcarata_GCA_905261645_1
BUSCO_Rotaria_socialis_GCA_905332285_1
BUSCO_Haliotis_ovina_GCA_977061855_1
BUSCO_Phorcus_lineatus_GCA_921293015_1
BUSCO_Phorcus_turbinatus_GCA_965237135_1
BUSCO_Tricolia_pullus_GCA_964273465_1
BUSCO_Marisa_cornuarietis_GCA_050626395_1
BUSCO_Biomphalaria_glabrata_GCF_947242115_1
BUSCO_Hippopus_hippopus_GCA_965239675_1
BUSCO_Meretrix_sp_MF1_GCA_049244355_1
BUSCO_Verpa_penis_GCA_048126915_1
BUSCO_Acanthopleura_gemmata_GCA_977063515_1
BUSCO_Sipunculus_nudus_GCA_026874595_2
BUSCO_Alitta_virens_GCA_932294295_1
BUSCO_Amphiduros_pacificus_GCA_949316495_1
BUSCO_Fimbriosthenelais_zetlandica_GCA_977030845_1
BUSCO_Sthenelais_limicola_GCA_942159475_1
BUSCO_Lepidonotus_clava_GCA_936440205_1
BUSCO_Terebella_lapidaria_GCA_949152475_1
BUSCO_Ophelia_neglecta_GCA_965285935_1
BUSCO_Sabella_spallanzanii_GCA_964340945_1
BUSCO_Ochetostoma_sp_New_Caledonia_GCA_977019775_1
BUSCO_Tubulanus_polymorphus_GCF_964204645_1
BUSCO_Leucocephalonemertes_aurantiaca_GCA_977061895_1
BUSCO_Lingula_anatina_GCA_051362555_1
0.2

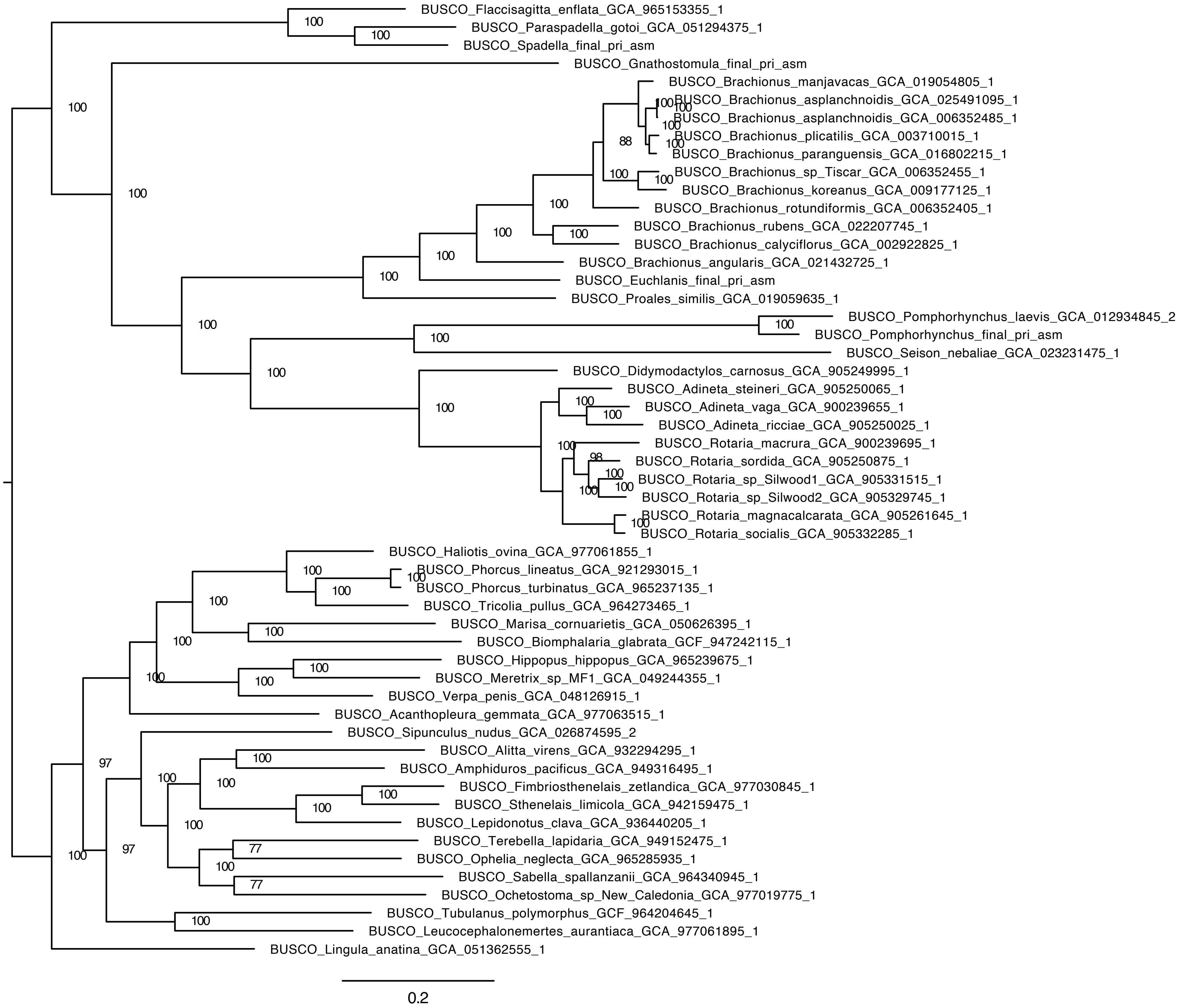
BUSCO_Flaccisagitta_enflata_GCA_965153355_1
BUSCO_Paraspadella_gotoi_GCA_051294375_1
BUSCO_Spadella_final_pri_asm
BUSCO_Gnathostomula_final_pri_asm
BUSCO_Brachionus_manjavacas_GCA_019054805_1
BUSCO_Brachionus_asplanchnoidis_GCA_025491095_1
BUSCO_Brachionus_asplanchnoidis_GCA_006352485_1
BUSCO_Brachionus_plicatilis_GCA_003710015_1
BUSCO_Brachionus_paranguensis_GCA_016802215_1
BUSCO_Brachionus_sp_Tiscar_GCA_006352455_1
BUSCO_Brachionus_koreanus_GCA_009177125_1
BUSCO_Brachionus_rotundiformis_GCA_006352405_1
BUSCO_Brachionus_rubens_GCA_022207745_1
BUSCO_Brachionus_calyciflorus_GCA_002922825_1
BUSCO_Brachionus_angularis_GCA_021432725_1
BUSCO_Euchlanis_final_pri_asm
BUSCO_Proales_similis_GCA_019059635_1
BUSCO_Pomphorhynchus_laevis_GCA_012934845_2
BUSCO_Pomphorhynchus_final_pri_asm
BUSCO_Seison_nebaliae_GCA_023231475_1
BUSCO_Didymodactylos_carnosus_GCA_905249995_1
BUSCO_Adineta_steineri_GCA_905250065_1
BUSCO_Adineta_vaga_GCA_900239655_1
BUSCO_Adineta_ricciae_GCA_905250025_1
BUSCO_Rotaria_macrura_GCA_900239695_1
BUSCO_Rotaria_sordida_GCA_905250875_1
BUSCO_Rotaria_sp_Silwood1_GCA_905331515_1
BUSCO_Rotaria_sp_Silwood2_GCA_905329745_1
BUSCO_Rotaria_magnacalcarata_GCA_905261645_1
BUSCO_Rotaria_socialis_GCA_905332285_1
BUSCO_Haliotis_ovina_GCA_977061855_1
BUSCO_Phorcus_lineatus_GCA_921293015_1
BUSCO_Phorcus_turbinatus_GCA_965237135_1
BUSCO_Tricolia_pullus_GCA_964273465_1
BUSCO_Marisa_cornuarietis_GCA_050626395_1
BUSCO_Biomphalaria_glabrata_GCF_947242115_1
BUSCO_Hippopus_hippopus_GCA_965239675_1
BUSCO_Meretrix_sp_MF1_GCA_049244355_1
BUSCO_Verpa_penis_GCA_048126915_1
BUSCO_Acanthopleura_gemmata_GCA_977063515_1
BUSCO_Sipunculus_nudus_GCA_026874595_2
BUSCO_Alitta_virens_GCA_932294295_1
BUSCO_Amphiduros_pacificus_GCA_949316495_1
BUSCO_Fimbriosthenelais_zetlandica_GCA_977030845_1
BUSCO_Sthenelais_limicola_GCA_942159475_1
BUSCO_Lepidonotus_clava_GCA_936440205_1
BUSCO_Terebella_lapidaria_GCA_949152475_1
BUSCO_Ophelia_neglecta_GCA_965285935_1
BUSCO_Sabella_spallanzanii_GCA_964340945_1
BUSCO_Ochetostoma_sp_New_Caledonia_GCA_977019775_1
BUSCO_Tubulanus_polymorphus_GCF_964204645_1
BUSCO_Leucocephalonemertes_aurantiaca_GCA_977061895_1
BUSCO_Lingula_anatina_GCA_051362555_1
0.2

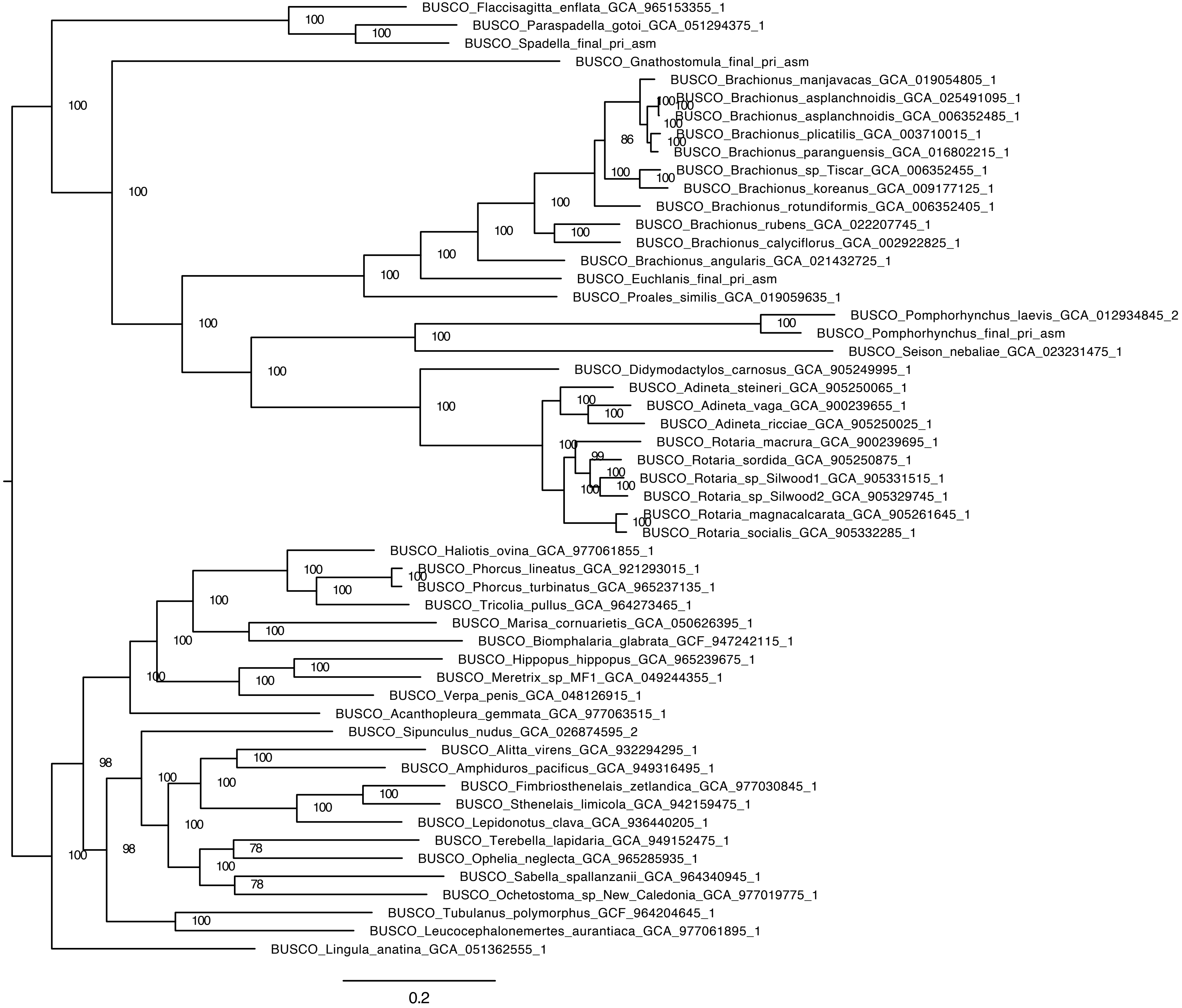
BUSCO_Flaccisagitta_enflata_GCA_965153355_1
BUSCO_Paraspadella_gotoi_GCA_051294375_1
BUSCO_Spadella_final_pri_asm
BUSCO_Gnathostomula_final_pri_asm
BUSCO_Brachionus_manjavacas_GCA_019054805_1
BUSCO_Brachionus_asplanchnoidis_GCA_025491095_1
BUSCO_Brachionus_asplanchnoidis_GCA_006352485_1
BUSCO_Brachionus_plicatilis_GCA_003710015_1
BUSCO_Brachionus_paranguensis_GCA_016802215_1
BUSCO_Brachionus_sp_Tiscar_GCA_006352455_1
BUSCO_Brachionus_koreanus_GCA_009177125_1
BUSCO_Brachionus_rotundiformis_GCA_006352405_1
BUSCO_Brachionus_rubens_GCA_022207745_1
BUSCO_Brachionus_calyciflorus_GCA_002922825_1
BUSCO_Brachionus_angularis_GCA_021432725_1
BUSCO_Euchlanis_final_pri_asm
BUSCO_Proales_similis_GCA_019059635_1
BUSCO_Pomphorhynchus_laevis_GCA_012934845_2
BUSCO_Pomphorhynchus_final_pri_asm
BUSCO_Seison_nebaliae_GCA_023231475_1
BUSCO_Didymodactylos_carnosus_GCA_905249995_1
BUSCO_Adineta_steineri_GCA_905250065_1
BUSCO_Adineta_vaga_GCA_900239655_1
BUSCO_Adineta_ricciae_GCA_905250025_1
BUSCO_Rotaria_macrura_GCA_900239695_1
BUSCO_Rotaria_sordida_GCA_905250875_1
BUSCO_Rotaria_sp_Silwood1_GCA_905331515_1
BUSCO_Rotaria_sp_Silwood2_GCA_905329745_1
BUSCO_Rotaria_magnacalcarata_GCA_905261645_1
BUSCO_Rotaria_socialis_GCA_905332285_1
BUSCO_Haliotis_ovina_GCA_977061855_1
BUSCO_Phorcus_lineatus_GCA_921293015_1
BUSCO_Phorcus_turbinatus_GCA_965237135_1
BUSCO_Tricolia_pullus_GCA_964273465_1
BUSCO_Marisa_cornuarietis_GCA_050626395_1
BUSCO_Biomphalaria_glabrata_GCF_947242115_1
BUSCO_Hippopus_hippopus_GCA_965239675_1
BUSCO_Meretrix_sp_MF1_GCA_049244355_1
BUSCO_Verpa_penis_GCA_048126915_1
BUSCO_Acanthopleura_gemmata_GCA_977063515_1
BUSCO_Sipunculus_nudus_GCA_026874595_2
BUSCO_Alitta_virens_GCA_932294295_1
BUSCO_Amphiduros_pacificus_GCA_949316495_1
BUSCO_Fimbriosthenelais_zetlandica_GCA_977030845_1
BUSCO_Sthenelais_limicola_GCA_942159475_1
BUSCO_Lepidonotus_clava_GCA_936440205_1
BUSCO_Terebella_lapidaria_GCA_949152475_1
BUSCO_Ophelia_neglecta_GCA_965285935_1
BUSCO_Sabella_spallanzanii_GCA_964340945_1
BUSCO_Ochetostoma_sp_New_Caledonia_GCA_977019775_1
BUSCO_Tubulanus_polymorphus_GCF_964204645_1
BUSCO_Leucocephalonemertes_aurantiaca_GCA_977061895_1
BUSCO_Lingula_anatina_GCA_051362555_1
0.2

**Phylogenetic Assessment: Macrosyntenic Analysis and "The Macrosyntenic Jackknife"**

Supplemental Figure 15: Oxford dot plots for *Branchiostoma* against the other four test species (*Adineta, Gnathostomula, Mizuhopecten* and *Paraspadella*), evidencing the strong significance in *Branchiostoma* vs. *Mizuhopecten*, and the more even distribution of dots across the remaining species comparisons. These comparisons were all derived from the conserved metazoan ALGS.

Supplemental Figure 16: Oxford dot plots for *Adineta* against *Gnathostomula* and *Paraspadella*, and *Paraspadella* against *Gnathostomula*. These comparisons were all derived directly from the data, in order to find Chaetognathifera-specific linkage groups. Note the relative even distribution across each graph.

Supplemental Figure 17: Supplemental Figure 16 with the addition of support values for each significant pair found within the jackknife analysis. Note that *Paraspadella* against *Gnathostomula* contains no significant pairs within the jackknife, and that of the 31 significant pairs found in the primary analysis between *Adineta* and *Gnathostomula*, only 2 are recovered robustly in the jackknife analysis.

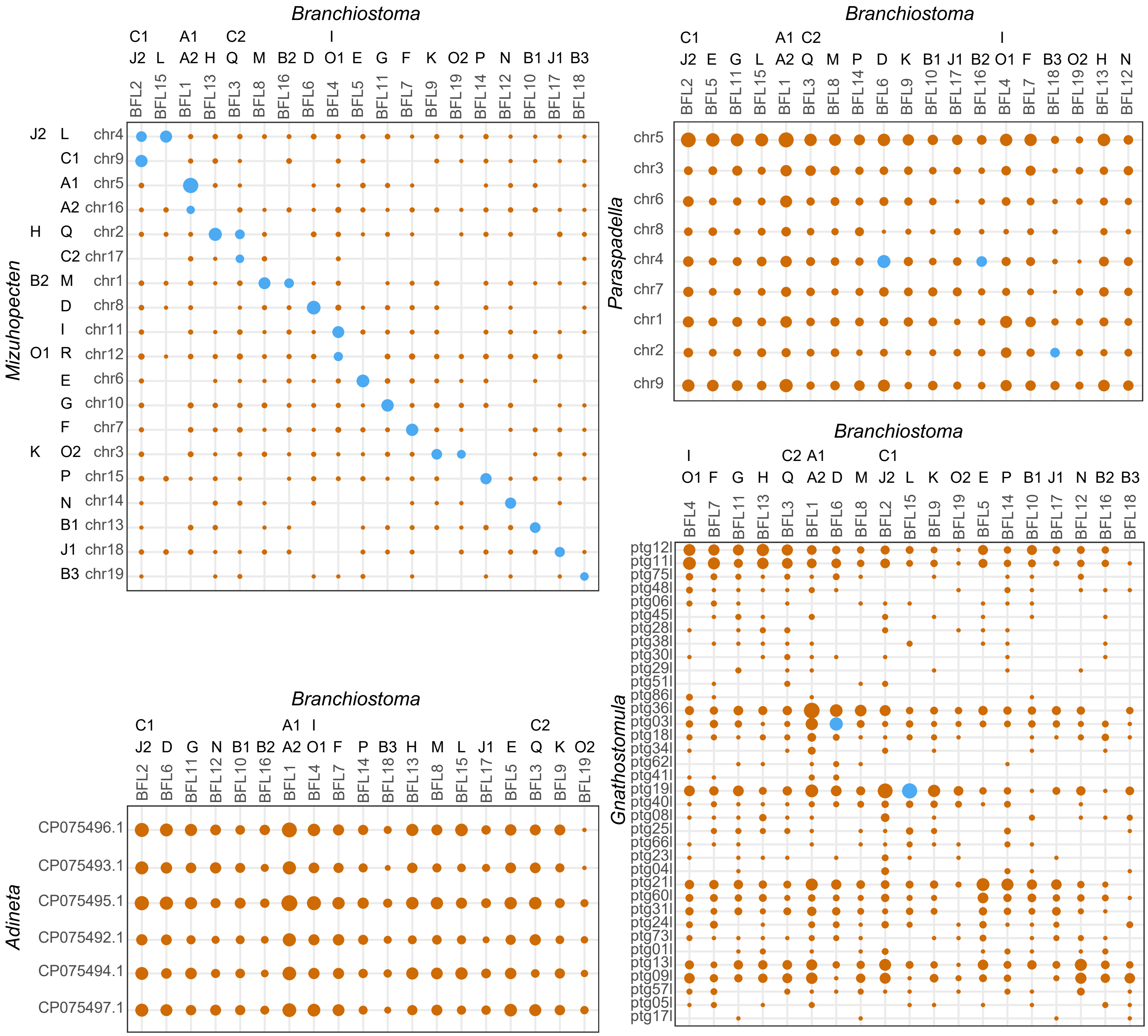

Branchiostoma
C1 A1 C2 I
J2 L A2 H Q M B2 D O1 E G F K O2 P N B1 J1 B3
BFL2 BFL15 BFL1 BFL13 BFL3 BFL8 BFL16 BFL6 BFL4 BFL5 BFL11 BFL7 BFL9 BFL19 BFL14 BFL12 BFL10 BFL17 BFL18
Mizuhopecten
J2 L chr4
C1 chr9
A1 chr5
A2 chr16
H Q chr2
C2 chr17
B2 M chr1
D chr8
I chr11
O1 R chr12
E chr6
G chr10
F chr7
K O2 chr3
P chr15
N chr14
B1 chr13
J1 chr18
B3 chr19
Branchiostoma
C1 A1 C2 I
J2 E G L A2 Q M P D K B1 J1 B2 O1 F B3 O2 H N
BFL2 BFL5 BFL11 BFL15 BFL1 BFL3 BFL8 BFL14 BFL6 BFL9 BFL10 BFL17 BFL16 BFL4 BFL7 BFL18 BFL19 BFL13 BFL12
Paraspadella
chr5
chr3
chr6
chr8
chr4
chr7
chr1
chr2
chr9
Branchiostoma
I C2 A1 C1
O1 F G H Q A2 D M J2 L K O2 E P B1 J1 N B2 B3
BFL4 BFL7 BFL11 BFL13 BFL3 BFL1 BFL6 BFL8 BFL2 BFL15 BFL9 BFL19 BFL5 BFL14 BFL10 BFL17 BFL12 BFL16 BFL18
Gnathostomula
ptg12l
ptg11l
ptg75l
ptg48l
ptg06l
ptg45l
ptg28l
ptg38l
ptg30l
ptg29l
ptg51l
ptg86l
ptg36l
ptg03l
ptg18l
ptg34l
ptg62l
ptg41l
ptg19l
ptg40l
ptg08l
ptg25l
ptg66l
ptg23l
ptg04l
ptg21l
ptg60l
ptg31l
ptg24l
ptg73l
ptg01l
ptg13l
ptg09l
ptg57l
ptg05l
ptg17l
Branchiostoma
C1 A1 I C2
J2 D G N B1 B2 A2 O1 F P B3 H M L J1 E Q K O2
BFL2 BFL6 BFL11 BFL12 BFL10 BFL16 BFL1 BFL4 BFL7 BFL14 BFL18 BFL13 BFL8 BFL15 BFL17 BFL5 BFL3 BFL9 BFL19
Adineta
CP075496.1
CP075493.1
CP075495.1
CP075492.1
CP075494.1
CP075497.1

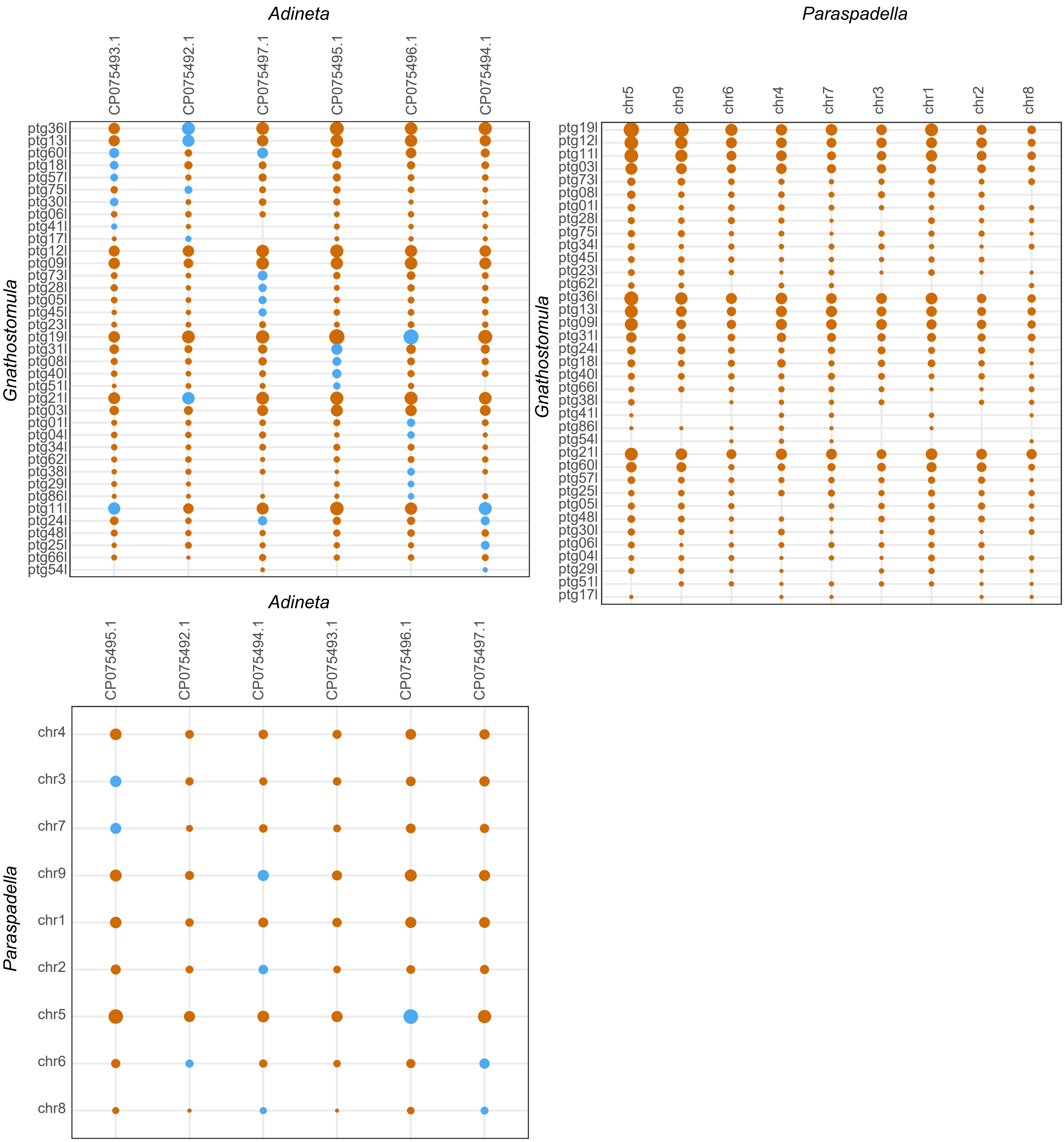
Adineta
CP075493.1
CP075492.1
CP075497.1
CP075495.1
CP075496.1
CP075494.1
Gnathostomula
ptg36l
ptg13l
ptg60l
ptg18l
ptg57l
ptg75l
ptg30l
ptg06l
ptg41l
ptg17l
ptg12l
ptg09l
ptg73l
ptg28l
ptg05l
ptg45l
ptg23l
ptg19l
ptg31l
ptg08l
ptg40l
ptg51l
ptg21l
ptg03l
ptg01l
ptg04l
ptg34l
ptg62l
ptg38l
ptg29l
ptg86l
ptg11l
ptg24l
ptg48l
ptg25l
ptg66l
ptg54l
Paraspadella
chr5
chr9
chr6
chr4
chr7
chr3
chr1
chr2
chr8
Gnathostomula
ptg19l
ptg12l
ptg11l
ptg03l
ptg73l
ptg08l
ptg01l
ptg28l
ptg75l
ptg34l
ptg45l
ptg23l
ptg62l
ptg36l
ptg13l
ptg09l
ptg31l
ptg24l
ptg18l
ptg40l
ptg66l
ptg38l
ptg41l
ptg86l
ptg54l
ptg21l
ptg60l
ptg57l
ptg25l
ptg05l
ptg48l
ptg30l
ptg06l
ptg04l
ptg29l
ptg51l
ptg17l
Adineta
CP075495.1
CP075492.1
CP075494.1
CP075493.1
CP075496.1
CP075497.1
Paraspadella
chr4
chr3
chr7
chr9
chr1
chr2
chr5
chr6
chr8

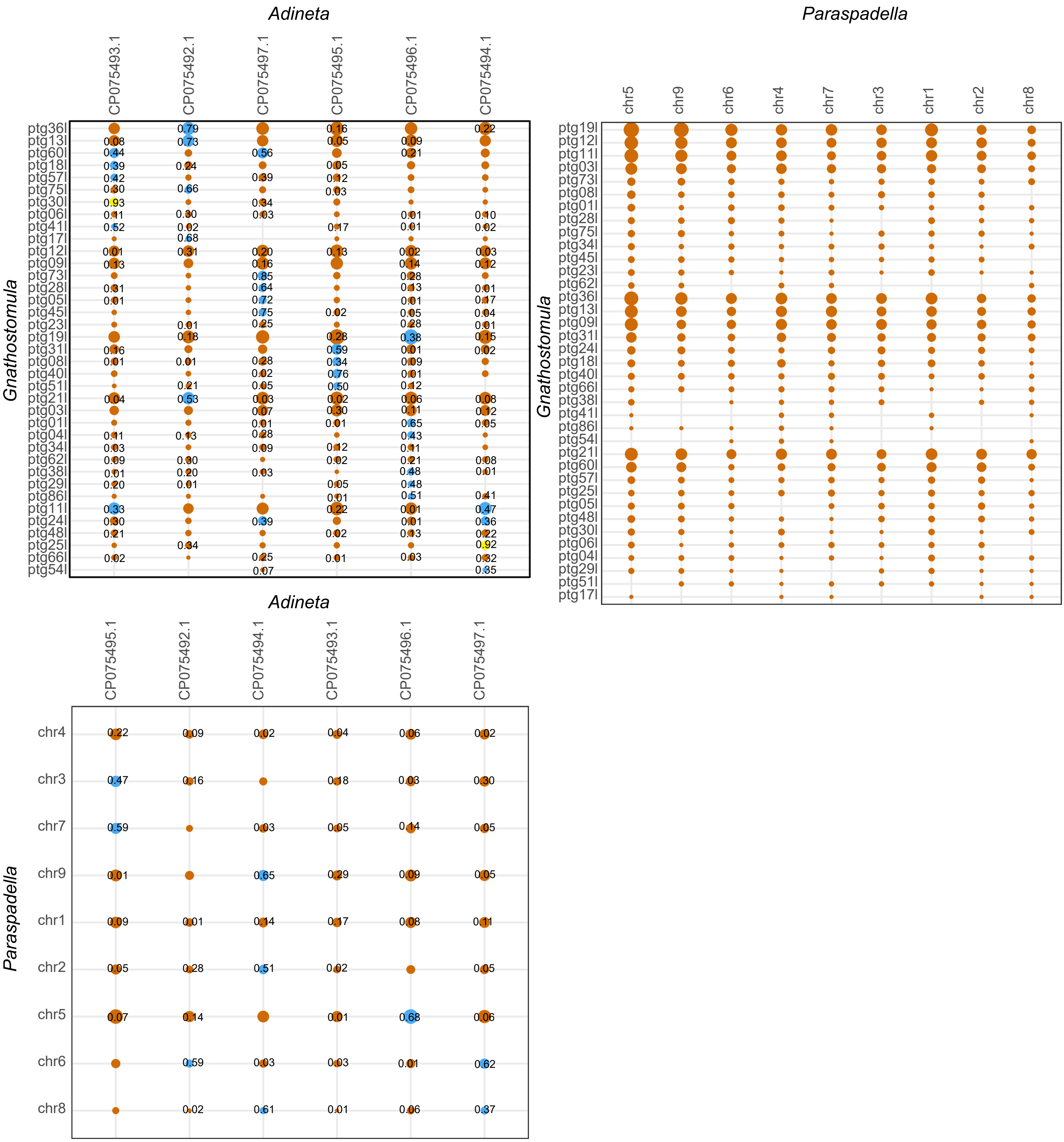
Adineta
CP075493.1
CP075492.1
CP075497.1
CP075495.1
CP075496.1
CP075494.1
Gnathostomula
ptg36l
ptg13l
ptg60l
ptg18l
ptg57l
ptg75l
ptg30l
ptg06l
ptg41l
ptg17l
ptg12l
ptg09l
ptg73l
ptg28l
ptg05l
ptg45l
ptg23l
ptg19l
ptg31l
ptg08l
ptg40l
ptg51l
ptg21l
ptg03l
ptg01l
ptg04l
ptg34l
ptg62l
ptg38l
ptg29l
ptg86l
ptg11l
ptg24l
ptg48l
ptg25l
ptg66l
ptg54l
Paraspadella
chr5
chr9
chr6
chr4
chr7
chr3
chr1
chr2
chr8
Gnathostomula
ptg19l
ptg12l
ptg11l
ptg03l
ptg73l
ptg08l
ptg01l
ptg28l
ptg75l
ptg34l
ptg45l
ptg23l
ptg62l
ptg36l
ptg13l
ptg09l
ptg31l
ptg24l
ptg18l
ptg40l
ptg66l
ptg38l
ptg41l
ptg86l
ptg54l
ptg21l
ptg60l
ptg57l
ptg25l
ptg05l
ptg48l
ptg30l
ptg06l
ptg04l
ptg29l
ptg51l
ptg17l
Adineta
CP075495.1
CP075492.1
CP075494.1
CP075493.1
CP075496.1
CP075497.1
Paraspadella
chr4
chr3
chr7
chr9
chr1
chr2
chr5
chr6
chr8
